# An interpretable machine learning algorithm to predict disordered protein phase separation based on biophysical interactions

**DOI:** 10.1101/2022.07.06.499043

**Authors:** Hao Cai, Robert M. Vernon, Julie D. Forman-Kay

**Author notes:** Shopify Inc., 150 Elgin Street 8th Floor, Ottawa, ON K2P 1L4, Canada. Amgen Inc. Research and Development, Burnaby, BC V5A 1V7, Canada.

## Abstract

Protein phase separation is increasingly understood to be an important mechanism of biological organization and biomaterial formation. Intrinsically disordered protein regions (IDRs) are often significant drivers of protein phase separation. A number of protein phase separation prediction algorithms are available, with many specific for particular classes of proteins and others providing results that are not amenable to interpretation of contributing biophysical interactions. Here we describe LLPhyScore, a new predictor of IDR-driven phase separation, based on a broad set of physical interactions or features. LLPhyScore uses sequence-based statistics from the RCSB PDB database of folded structures for these interactions, and is trained on a manually curated set of phase separation driver proteins with different negative training sets including the PDB and human proteome. Competitive training for a variety of physical chemical interactions shows the greatest importance of solvent contacts, disorder, hydrogen bonds, pi-pi contacts, and kinked-beta structure, with electrostatics, cation-pi, and absence of helical secondary structure also contributing. LLPhyScore has strong phase separation prediction recall statistics and enables a quantitative breakdown of the contribution from each physical feature to a sequence’s phase separation propensity. The tool should be a valuable resource for guiding experiment and providing hypotheses for protein function in normal and pathological states, as well as for understanding how specificity emerges in defining individual biomolecular condensates.

**Graphical Abstract:** 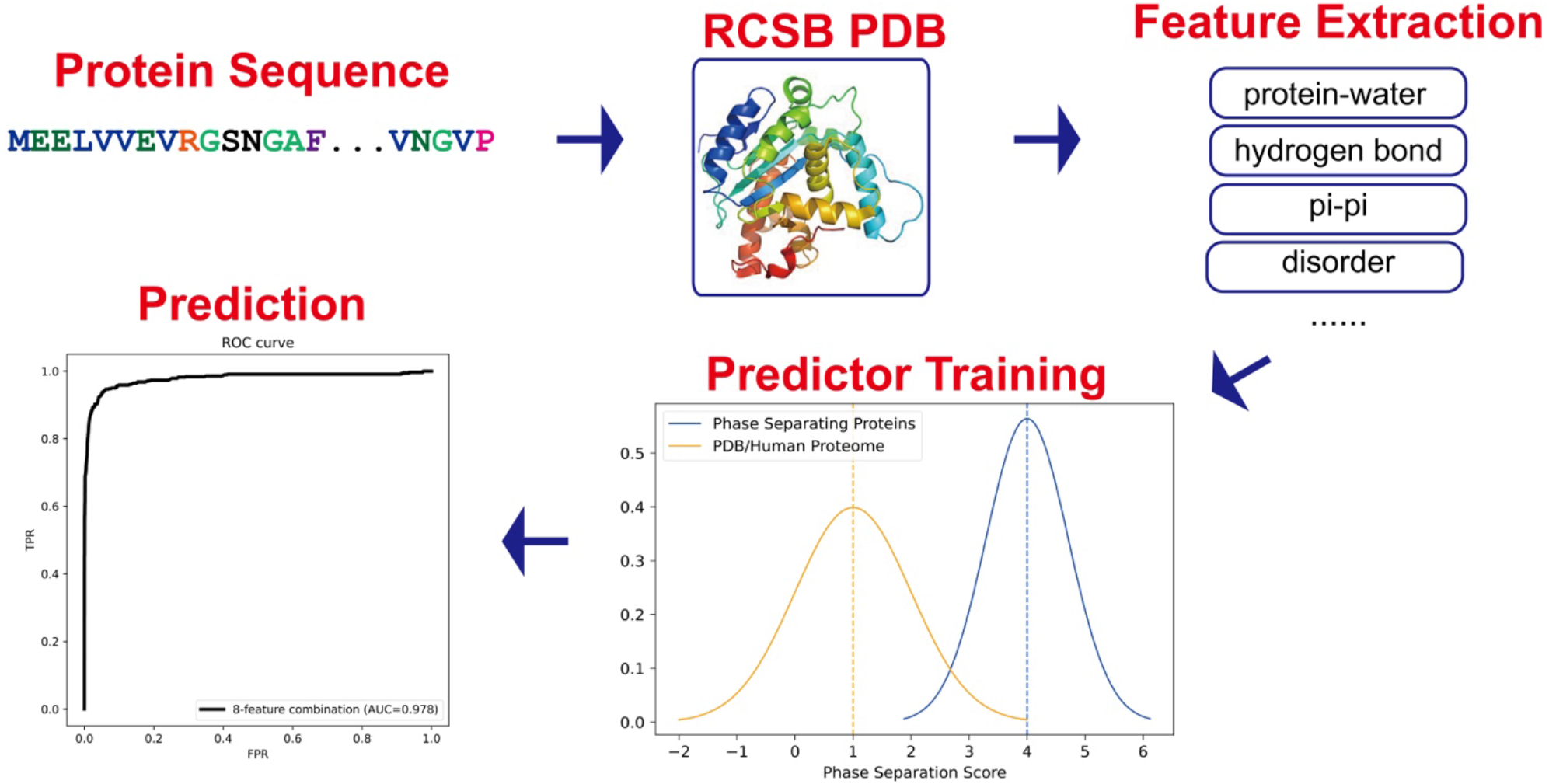

## Introduction

Protein phase separation has recently been recognized as an important mechanism of compartmentalization in cells contributing to formation of biomolecular condensates.[1, 2] Liquid-liquid phase separation (LLPS) is not the only physical phenomenon that can contribute to formation of these condensates, with these including sol-gel transitions and phase separation coupled to percolation (PSCP)[1, 3, 4]. Here we use the term “phase separation” as an imprecise shorthand for these mechanisms that rely on exchanging multi-valent interactions[5] that give rise to biomolecular condensates. Biomolecular condensates are found in a wide range of biological contexts, including intracellular condensates and membraneless organelles[6, 7] such as signaling puncta[8, 9], nuclear pores[10], transcription centers[11], and mRNA transport granules[12–14]; and extracellular biological materials such as in elastin[15–17], mussel foot[18, 19], and squid beak[20–23]. Biomolecular condensates are also implicated in pathological aggregation (e.g., ALS[24] and Alzheimer’s disease[25]).

Physical mechanistic understanding of protein phase separation in all its complexity is challenged due to the richness and versatility of its driving forces. Phase separation can be affected by a large set of sequence-dependent factors, with a significant role of intrinsically disordered protein regions (IDRs) in many cases. For phase separation driven by IDRs, numerous weak interaction forces have been highlighted as contributing, including electrostatic interactions[26–28], pi-pi stacking[29–31], cation-pi interactions[19, 26, 32], and hydrogen bonding[33, 34], with multiple forces often implicated as seen in low-complexity aromatic-rich kinked segments (LARKS)[33] which show kinked-beta backbone hydrogen bonding and aromatic sidechain interactions. In elastin and elastin-like peptides, the hydrophobic effect is important for phase separation [35, 36]. For phase separation driven by folded domains, specific sequence motifs, SLiMs[37], and their cognate folded binding domains are key; while these are an important driver of biological phase separation, our focus here is IDR-driven phase separation.

Since most of the physicochemical factors that facilitate phase separation are sequence-dependent, there have been numerous efforts to use statistical learning to draw physical insights from known phase-separating sequences, i.e., to predict whether a sequence will undergo phase separation by comparing against tested sequences, as previously summarized in a 2019 review[38]. However, the algorithms mentioned in that review focus on specific categories of condensates or biophysical features, and can only predict a subset of phase-separating proteins with high confidence. There is a high level of correlation among biophysical features, e.g., pi-pi and solvent interactions[29], electrostatic interactions and hydrophobic interactions[28, 39], but none of these algorithms can estimate phase-separation propensities based on all these physical forces, limiting the overall predictive capability of these “first-generation” predictors. In subsequent work[40], a machine-learning based prediction tool (PSPredictor) that uses word2vec sequence encoding and Gradient Boosting Decision Tree (GBDT) model outperformed all the “first-generation predictors” and achieved a 96% prediction accuracy. However, because of the design of word2vec encoding[41], its prediction results cannot provide quantitative information about contributions from different driving forces, and therefore it lacks clear physical interpretability. Recently, a number of additional tools have been developed to quantify phase-separation propensity. One of these, PSPer, focuses on prediction of prion-like RNA-binding proteins that phase separate using a Hidden Markov Model (HMM).[42] PSPer showed good predictability (0.87 Spearman correlation score between its output and the critical concentration of FUS-like proteins), however has limited ability predicting phase-separating proteins that are not RNA-binding. Another, ParSe, combines two physical features, hydrodynamic size of monomeric proteins and beta-turn propensity estimated from polymer models, to predict phase separation propensity, however uses only composition and not residue context when making predictions.[43] A third, PSAP, uses compositional bias of phase-separating proteins and 10 amino-acid biochemical features as features for a random forest classifier with a 0.89 AUROC (area under the receiver operating characteristics curve), yet also lacks residue context in the prediction.[44]

A major issue in developing a phase separation predictor is the selection of a negative training set. Most recently developed predictors use sequences of the folded proteins in the RCSB Protein Data Bank (PDB)[45] as the negative set[29, 40], however this leads to a bias towards a final classification algorithm that distinguishes between intrinsically disordered proteins/regions and folded proteins, since most proteins found to phase separate are IDPs or have IDRs. This classification does not identify driving forces of phase separation, however, since many IDPs/IDRs are not phase-separating. To avoid this, in other cases the human proteome was chosen as the negative benchmark, bringing in higher structural complexity.[33, 46, 47] Another computational approach that has been developed to predict propensity to phase separate, FuzDrop, has estimated that up to 40% of the human proteome potentially can undergo phase separation under certain conditions[48]. Therefore, it is clear that training any phase separation prediction algorithm on a negative dataset that includes many false negatives leads to significant challenges.

In the present work, we based our strategy on the idea that a combination of multiple different physical interactions drives phase separation, and developed a machine learning-based predictor (LLPhyScore) that predicts based on a set of phase-separation-related physical interactions or features. While “LLPhyScore” was named by combining the acronym “LLPS” and “physical feature-based scoring”, the tool is not focused only on “liquid-liquid phase separation” but is intended as a general predictor of phase separation by various mechanisms that rely on exchanging multi-valent interactions within IDRs. We adapted the constrained training approach from our previous work on PScore[29] that focused on planar pi-pi interactions and extended it to a total of 16 physical measurements or features. Our predictor development process was divided into two stages. In the first stage, we acquired sequence-based statistics (contact frequency/number of atoms/structure probability) from the PDB database of folded structures for each physical feature/interaction. We divided these observations by distinct residue pairs with varying sequence separation and developed a statistical method to predict expected physical feature values given a protein sequence. In the second stage, we trained the predictor to rank sequences by the weighted combination of expected physical feature values. During the predictor training, we used a genetic algorithm to optimize (i) the number of physical features to utilize in our final algorithm; (ii) the direction of contribution to phase separation (sign) of each feature; and (iii) the weights of each physical feature chosen for the final algorithm. The predictive model is a 3-layer “neural network”-like model that infers statistics of the input sequence based on physical features, residue types and residue counts and positions. The training revealed the better-appreciated importance of pi contacts and disorder, including solvent interactions, but also the less-well-appreciated significance of hydrogen bonds and kinked-beta structure. In order to address the “imperfect negative dataset” issue, we used three different negative training sets: the PDB, a curated human proteome and a mixture of PDB plus curated human proteome, and examined their impact on the final predictor’s performance. The final predictor (LLPhyScore) achieved excellent predictive power (AUROC of 0.978) and demonstrated significant physical interpretability by providing a quantitative breakdown of the contribution from each physical feature to a sequence’s propensity to phase separate.

## Results and Discussion

### Data Preparation

The overall data preparation workflow is shown Figure 1 and includes the following 4 parts:

**Figure 1.**
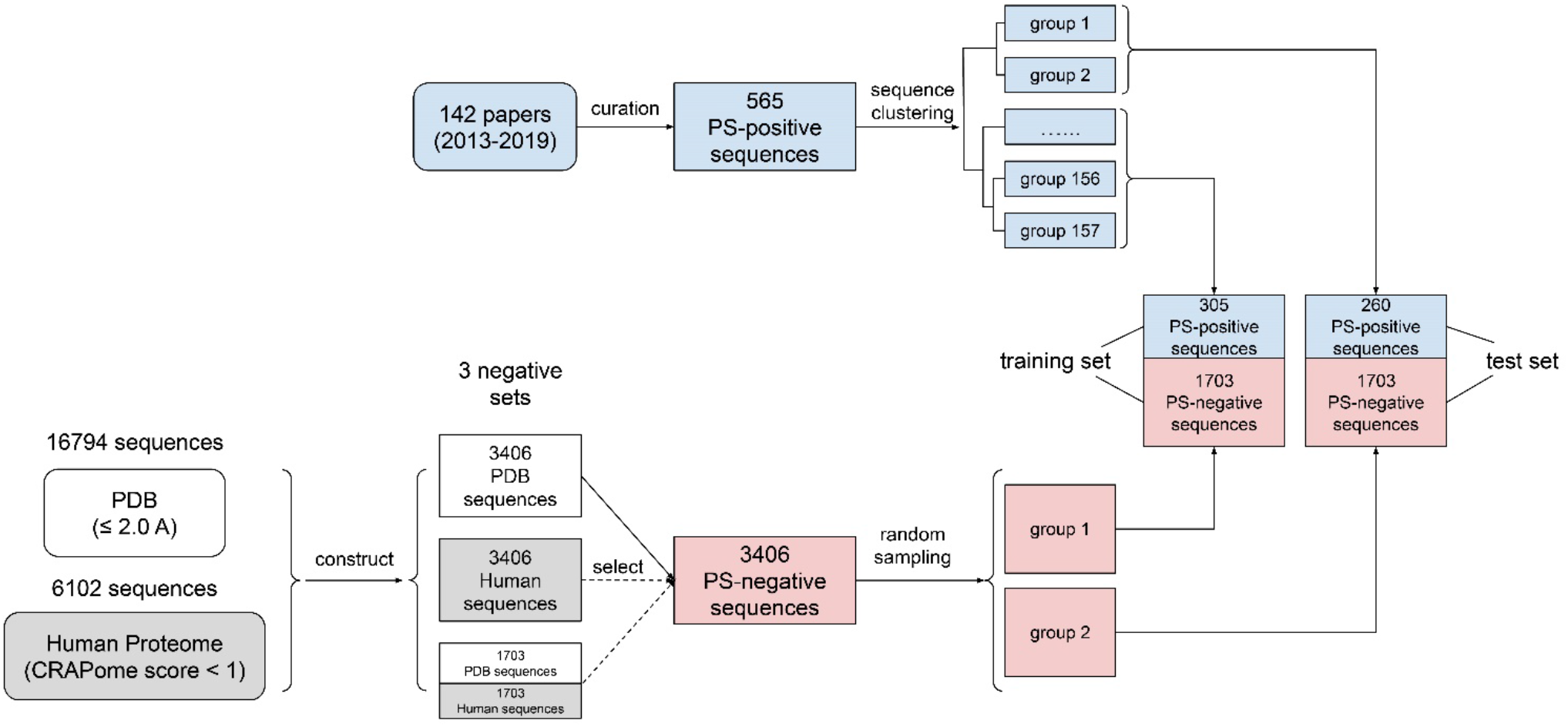
Data curation workflow. A schematic diagram of how data for training were obtained and processed.

(1) Curation of phase separation-positive (PS-positive) sequences: In this paper, we defined PS-positive proteins as proteins that can undergo phase separation on their own *in vitro*. We noticed that in several recently published phase separation sequence databases, including LLPSDB[49], PhaSepDB[50] and PhaSePro[51], two main issues exist: (i) Many phase-separated systems are multi-component, comprised of “scaffold” proteins that are PS-positive and “client” proteins that are phase separation-negative (PS-negative) on their own. However, “client” proteins were often mislabeled as PS-positive. (ii) There were many sequence errors (e.g. missing fluorescence tags; incorrect species; mishandled mutations and cleavages). To tackle these issues, we screened 142 recent papers (**Supplementary Table S10)** from Jul 2013 to Jan 2019, excluded sequences that can only undergo phase separation with DNA/RNA/other proteins from our positive set, and manually extracted 565 sequences (see **Supplementary File S1**) as our PS-positive set (workflow shown in Figure 1). Then, we used LLPSDB and PhaSepPro to cross-check the sequences.

(2) Clustering of PS-positive sequences (for train-test split): A common practice used in other related work[29, 40, 47] is to split training and test data to use previously discovered sequences for training and newly found sequences for test. However, we noticed there are many similar sequences reported at different times (e.g. sequences from a family that was worked on by the same lab in many years, or different mutants of the same wild type), so performing a time-based split will cause high bias as we are training and testing similar samples on the algorithm. This issue would be even more problematic considering the limited sample size for PS-positive proteins reported at the start of our work (<1000 samples). Therefore, before splitting the training and test set, we applied hierarchical clustering on 565 PS-positive sequences, and obtained 157 sequence groups, as shown in Supplementary Figures S1 and S2, where sequences within the same group have pairwise similarity of higher than 50%. The subsequent train-test split was then conducted based on sequence groups instead of individual sequences, so that training and testing set proteins are derived from separate sequence groups.

(3) PS-negative training sets: We created two negative sequence databases: (i) PDB sequence database, from which we collected 16794 sequences (see **Supplementary File S2**) from high-resolution (≤2.0 A) structures in PDB, and (ii) curated human proteome sequence database, from which we collected 20380 human proteome sequences (see **Supplementary File S3**) from Uniprot and removed sequences with either null values or high values (top 20%) in CRAPome[38, 52]. We chose CRAPome as the method of filtering out phase separation-prone sequences because it is an empirical measurement of non-specific interactions in human proteins rather than predictions[52]. This results in a “clean” human set of 6102 sequences (**Supplementary Table S6**). It is worth noting that false negatives exist in both the PDB and human negative sequence data sets. While we have attempted to minimize false negatives, even the PDB set is compromised to an unknown degree. Certainly, there are fewer positives in these sets than in the known PS-positive sequences, but perfect discrimination is likely impossible because the training sets are not gold standard truths, and the percentage of human proteome and PDB derived sequences that undergo phase separation is unknown.

Since the positive sample size is much smaller than the negative sample size, we then randomly selected 3406 sequences each from the (i) PDB sequence database and (ii) curated human sequence database, and constructed two negative sets: (a) PDB set (3406 sequences) and (b) curated human set (3406 sequences). Finally, we mixed (a) and (b) at 1:1 ratio, and built (c) mixture of PDB + curated human set of 3406 sequences (randomly selecting 1703 from PDB and 1703 from human). During the Predictor training step, the PDB set was used as the main set for determining both the “signs” of features and the number of features to keep; then, all three sets were used to optimize the “weights” of features, comparing the three final predictors’ performances with each other.

(4) Construction of training/test/evaluation sets: For the training of predictor models and optimization of model parameters, we constructed training and test sets initially by adopting a 70%-30% train-test split ratio to PS-positive and negative sample sets in steps (1) to (3). For positive samples, random sampling was conducted at the clustered group level until >30% sequences went to the test set. However, due to the existence of large sequence groups (30- 50 sequences), the end result actually was close to an even ratio with 305 sequences in the training set and 260 in the test set, as shown in Figure 1. Therefore, we then used an even ratio between training and test sets for the negative samples, where the random split was conducted at the sequence level, given that the issue with similar sequences does not affect the negative set.

For evaluation of the final models’ performances trained on different negative databases (PDB, human, human+PDB) on a defined dataset, we constructed **Evaluation set 1** composed of the entire PS-positive set (565 sequences) and the entire PDB proteome (16794); for the comparison of our predictor’s performance with other state-of-the-art phase separation predictive algorithms, we constructed **Evaluation set 2** composed of the entire PS-positive set (565 sequences) and the entire human proteome (20380 sequences).

For more details, see Supplementary Table S1.

### Construction of physical feature collection in LLPhyScore

We made two core assumptions in this work to develop a sequence-based predictive algorithm: (i) phase separation is driven by multiple physical forces and structural factors; (ii) for any phase-separated system, these forces and features together build up the system’s free energy to drive the phase transition. Then, we constructed a set of 16 sequence-based, phase separation-related physical features, including weak interactions and structural patterns. More details on each of these can be found in **Methods** and Supplementary Table S2.

#### Protein-water interactions

As pointed out by others in the field [36, 53], protein-water interactions represent an largely overlooked driving force in phase separation because of its synergistic nature with other interactions such as pi-pi, hydrogen-bonding and electrostatic interactions[29, 36, 53]. Here, we considered it as a separate force/feature and explored its role in phase separation. We defined protein-water interaction by contacts, and measured solvation contacts and hydrophobic contacts in two inversely correlated terms, a residue-water interaction count and a residue-carbon interaction count. The frequency measurement followed the same protocol as for pi-pi interactions in PScore.[29]

#### Helices and strands

While most phase-separating proteins contain IDRs that play a significant role in driving phase separation, in some cases[33, 37, 54], these IDRs transiently sample folded structure (either helices or beta-structures with varied dynamics and sizes) that can play a critical role. Here, we used the DSSP program to assign secondary structure[55] and enable helices (H) and strands (E) to be considered as contributing features. Disorder was categorized as a separate feature, because most reported phase-separating systems are IDR-driven, and the statistics are highly skewed towards disorder, which could be detrimental to the algorithm training. Boolean values (true or false) instead of frequency were utilized for helices and strands.

#### “Long-range” and “short-range” disorder

Due to the large difference in structural context between short (<5 residues long) and long (>15 residues long) disordered regions[56, 57], disorder was divided into these two categories. Here we defined the presence or absence of disorder as Boolean values (true or false), and measured disorder based on DSSP assignment of consecutive residues in a sequence.

#### Long-range and short-range electrostatic interactions

Electrostatic interactions have been established as another important driving force for phase separation, especially for highly charged sequences in complex coacervation systems, such as for the tau protein[58]. Here we defined electrostatic interaction using coulombic interaction energy with atomic partial charges taken from the Talaris2014 force field[59], dividing interaction energies by the sequence separation of involved atom pairs into short-range (<5 residues apart) and long range (>= 5 residues apart). We note that complex coacervation will not be predicted as the approach is based on phase separation of a single protein.

#### Long-range and short-range hydrogen bonds

Hydrogen bonding was found in some cases to co-exist with other driving forces, including pi-pi contacts[21] and protein-solvent interactions[60]. In this work, we considered it as a separate force and explored its role in phase separation. We used the PHENIX software suite[61] to identify OH-N hydrogen bonds and measured inter-residue hydrogen-bond interaction counts in short-range (<5 residues apart) and long range (>= 5 residues apart) contexts.

#### Long-range and short-range pi-pi interactions

We utilized our previous approach from the PScore phase separation predictor based on planar pi-pi contacts[29], determining contact frequency for residue pairs in the context of short-range (<5 residues apart) and long range (>= 5 residues apart) interactions.

#### Long-range and short-range cation-pi interactions

Cation-pi was found to have specific residue-type preference among cations arginines and lysines and aromatic phenylalanines, tyrosines and histidines, and substitutions of preferred residues in certain systems cause drastic change in phase separation behavior[62]. In order to crudely estimate potential cation-pi interactions we adapted the electrostatic potential by adding partial negative charges above and below the planes of aromatic ring systems, balanced with positive charge in plane, and then calculated the change relative to our standard electrostatic term. These measurements were again split into short-range (<5 residues apart) and long range (>= 5 residues apart).

#### Kinked beta-strands (K-Beta)

It has been observed that specific sequences from some phase-separating proteins can form fibrils of kinked-beta strands, with beta-strand hydrogen-bonding occurring without extended backbone torsion angles and forming fibrils similar to amyloids.[33] Prediction of this feature has previously been done by energetic assessment of a sequence’s ability to adopt the topology found in these fibril structures[63], and we created an analogous classification strategy by identifying sequences in the PDB that are similar or dissimilar (measured by backbone RMSD) to these kinked-beta strands.[64] Two Boolean metrics, K-Beta similarity and K-Beta non-similarity, were determined from RMSD values after structural superposition calculations.

Based on the above 16 (8 pairs) features, we designed a sequence representation system (See **Methods** and Figure 2) to convert a sequence into inferred residue-level feature values (frequencies/numbers/Booleans).

**Figure 2.**
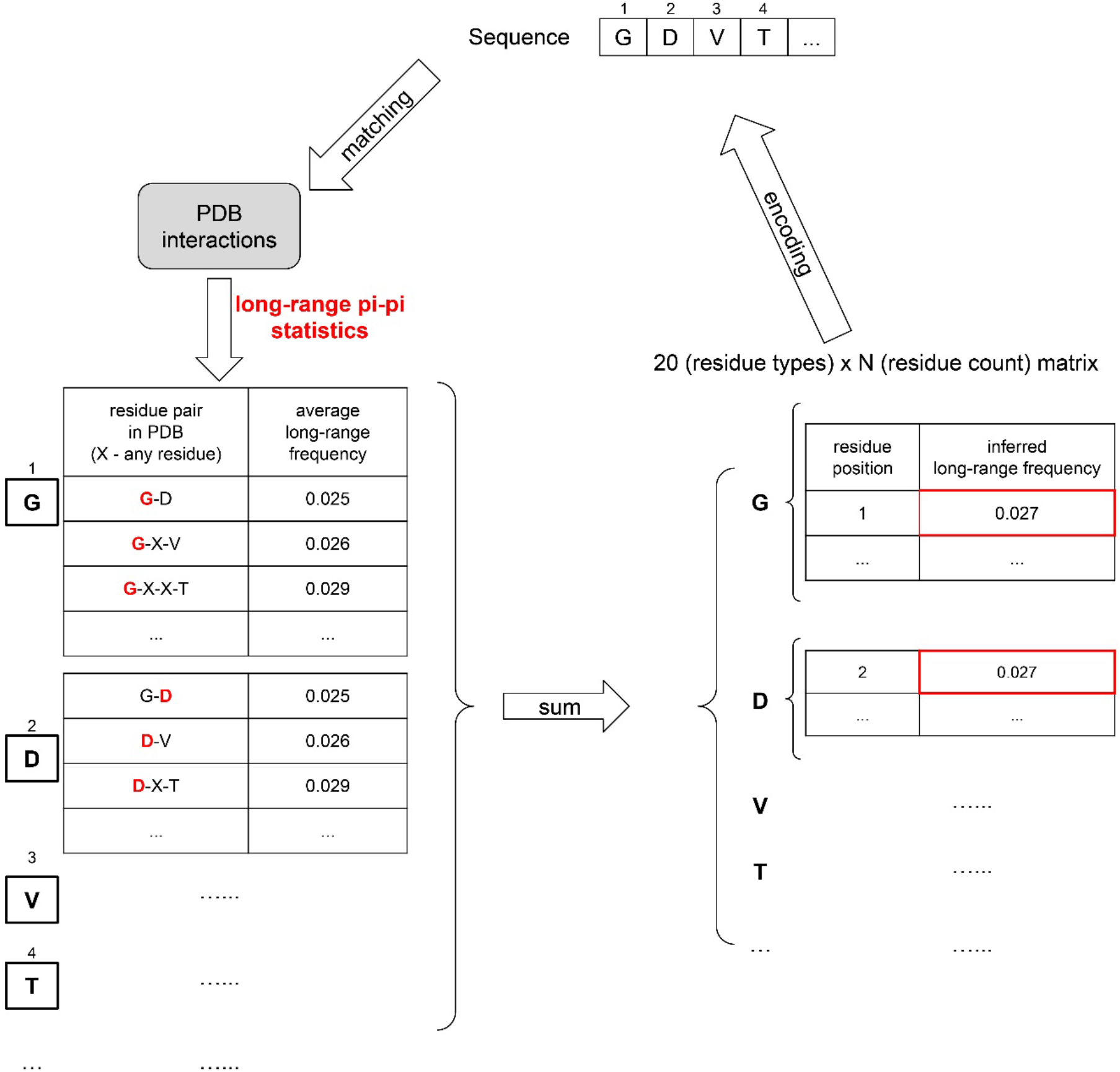
Physical interaction- and structure-based feature extraction. An example is given of the feature representation of sequences for the sequence “GDVT” converted to the pi-pi (long-range) feature matrix.

### Predictor training

The concept of “predictor training” in this work means: (i) for a specific sequence, the algorithm outputs a summed score calculated by a weighted combination of expected physical feature values, and (ii) during the predictor training, we optimize the combination of physical features, as well as the “weight” for each feature. The workflow of predictor training is shown in Figure 3.

**Figure 3.**
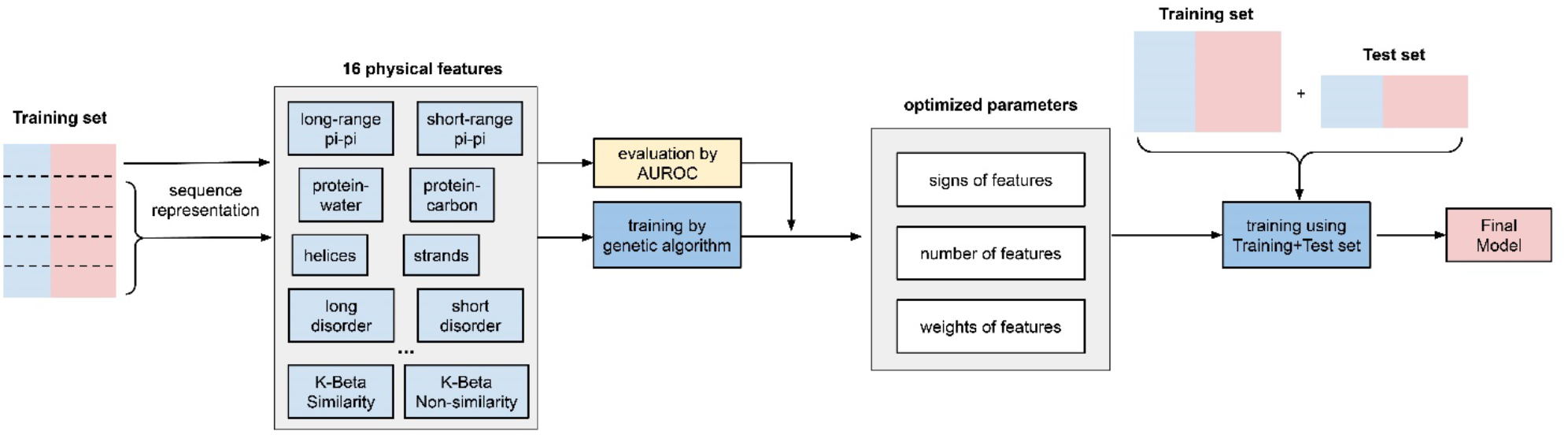
Predictor training workflow. A schematic diagram of the steps in training is shown.

The predictor training has three outcomes, described here:

#### (1) “Signs” of features were determined using individual feature training

From a physical perspective, some features in our list promote phase separation, while other features inhibit it. Therefore, before combining the 16 features, we first trained each feature individually and let the algorithm decide the “direction” (positive or negative) of its contribution to phase separation. As shown in Figure 4 and Supplementary Table S3, the features that were found to contribute negatively are protein-carbon interactions, helical secondary structure, long-range electrostatic interactions, both short- and long-range cation-pi interactions and kinked-beta (K-Beta) non-similarity. These results for electrostatic interactions are surprising but it is not obvious how our approach deals with locally repulsive interactions (clustered charges) that may interact with oppositely charged clusters and how well our crude estimate of cation-pi interactions works. Certainly, complex coacervation is not predicted as this tool is limited to homotypic phase separation, i.e., involving a single protein sequence.

**Figure 4.**
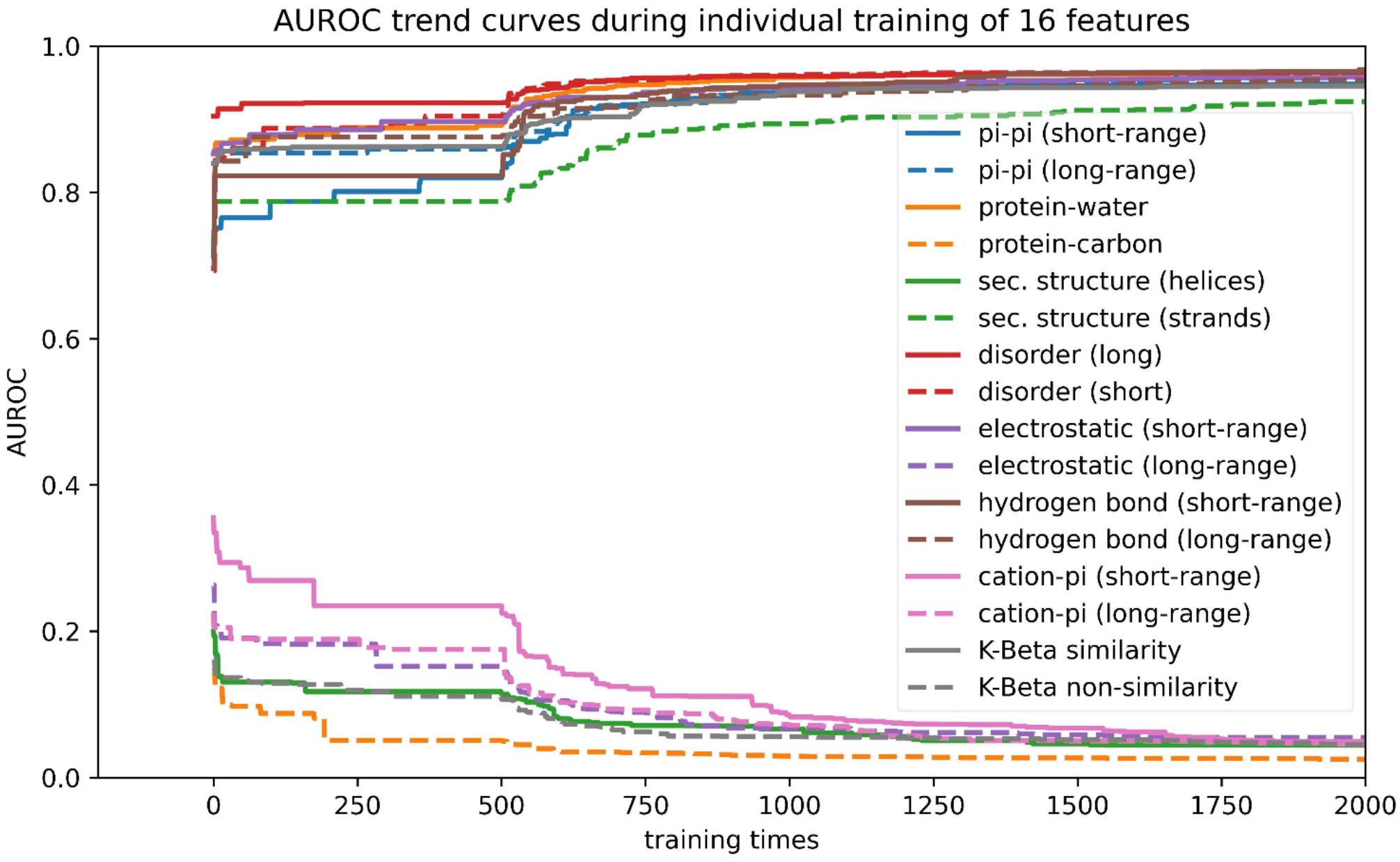
Direction of correlation of features with phase separation. Training curves of 16 features to reveal the direction of correlation of each feature with phase separation. Features that rise towards AUROC=1.0 have “positive” features; features that decline towards AUROC=0.0 have “negative” signs.

#### (2). The number of features to include was determined using competitive feature training

After determining the “signs” of the features and applying them, we combined 16 features and allowed them to “compete” with each other through “competitive” training, then ranked their importance based on the final contribution (positive or negative) of each feature to distinguishing phase-separating from non-phase-separating proteins, as shown in Figure 5. While all 16 features achieved an average z-score greater than 1.5, the average z-scores for protein-water, protein-carbon, long-range hydrogen bond and long-range pi-pi were larger than 3.0 and those for disorder (within both short and long segments) and kinked-beta similarity were larger than 2.5. The competitive training approach enables quantitative comparison of the significance of these physical interactions in phase separation over the input positive set. We then came up with 3 different combinations of features according to the ranking, combining the top 8 or top 12 features based on ranking or combining all 16 features, in order to identify the minimal number of features that provides both good performance and physical interpretability. We conducted competitive training on each of the 8, 12, and 16 feature algorithms and assessed their performance. As shown in Supplementary Table S4, the combinations of 12 and 16 features did not demonstrate better performance than the combination of 8 features. To avoid overtraining, as well as improve the potential for physical interpretability, we chose the 8-feature combination in the final predictor training.

**Figure 5.**
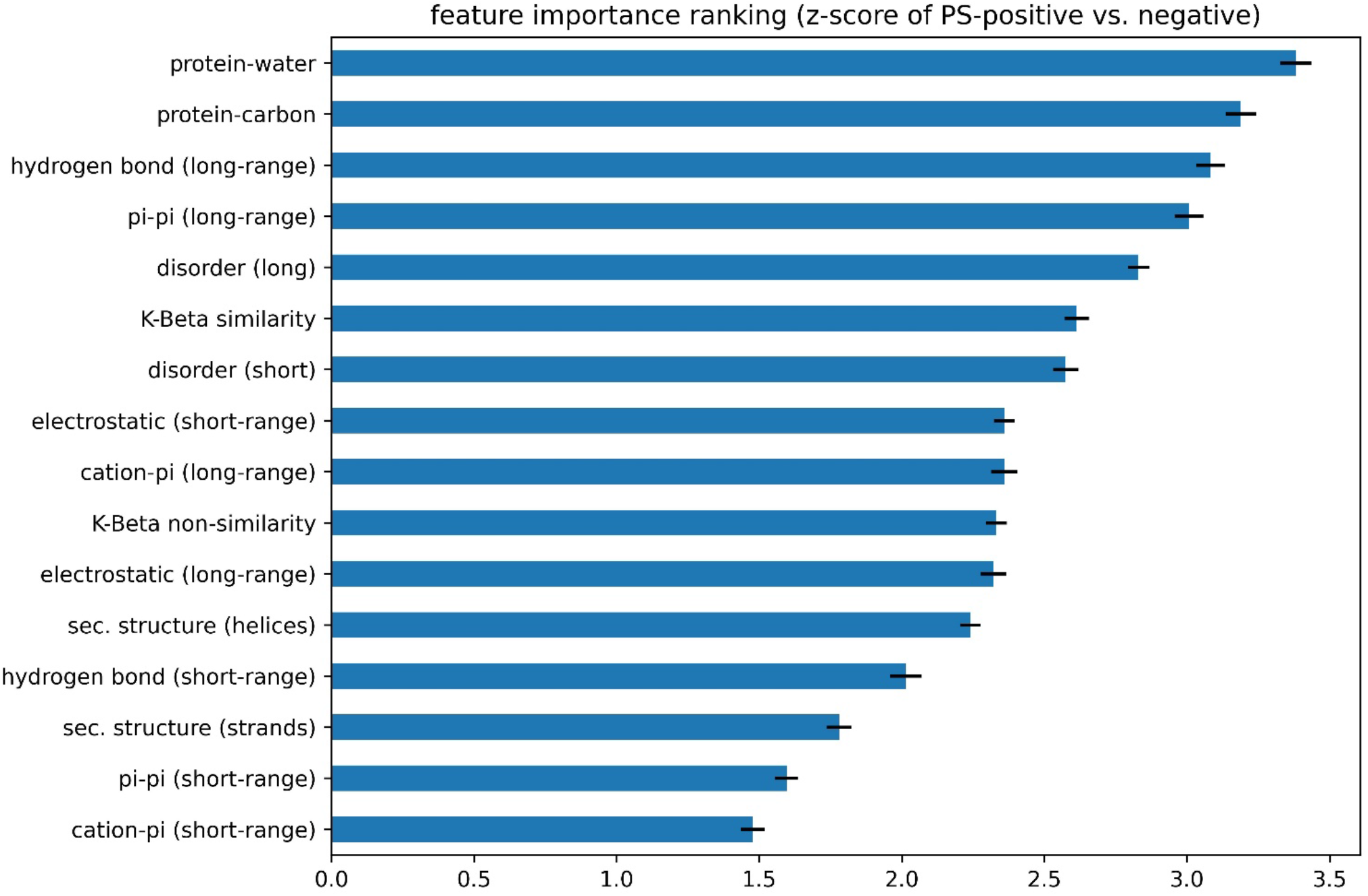
Ranking of the importance of features to phase separation. The z-score of PS-positive sequences’ individual feature values against the mean PS-negative sequences’ values.

#### (3) The “weights” of features in the final predictor were determined using competitive feature training on the entire dataset

We built the final predictor (“LLPhyScore”) and optimized the “weights” for the chosen 8 features with their respective signs by competitive training on Training set 1 (Supplementary Table S1) and tested the model performance on Test set 1 (Supplementary Table S1). We chose AUROC as the model performance metric, which is 0.969 for training and 0.942 for test (Supplementary Table S4) with the PDB as the negative set. This indicates that minimal overtraining occurred during the “weights” optimization. Then, we trained the “weights” again on (Training set 1 + Test set 1) to get the final predictor called “LLPhyScore-PDB model” based on its use of the PDB as a negative set. LLPhyScore-PDB model achieved an AUROC value of 0.978 (Supplementary Figure S3) on Evaluation set 1 (Supplementary Table S1) and good separation between positives and negatives (Supplementary Figures S4 and S5).

### Model performance comparison against different negative training sets

As noted previously, there is no perfect negative sample set for phase separation predictor development; therefore, after we trained the LLPhyScore-PDB model (on Training set 1 + Test set 1), we also trained the LLPhyScore-Human model (on Training set 2 + Test set 2) and the LLPhyScore-Human+PDB model (on Training set 3 + Test set 3), and evaluated the three final models using both Evaluation set 1 and Evaluation set 2. The results shown in Figure 6 and Supplementary Figures S3, S4 and S5 indicate that the PDB model showed the best performance on Evaluation set 1, against the PDB (AUROC of 0.978), but worst performance on Evaluation set 2, against the human proteome (AUROC of 0.824); the Human model showed the best performance on Evaluation set 2 (AUROC of 0.941) and worst performance on Evaluation set 1 (AUROC of 0.908). This indicates that the negative training set of different models has a significant impact on the final model performance. The model using only folded proteins from the PDB as negative training sequences tends to have less power to generalize on Evaluation set 2 (human proteome), which contains many disordered regions. On the other hand, the model only using Human proteins as the negative training sequences still has a strong ability to discriminate most PS-positive sequences from PDB sequences in Evaluation set 1. This is also reflected by the fact that the Human+PDB model showed a more balanced result for Evaluation set 1 (AUROC of 0.947) and Evaluation set 2 (AUROC of 0.933).

**Figure 6.**
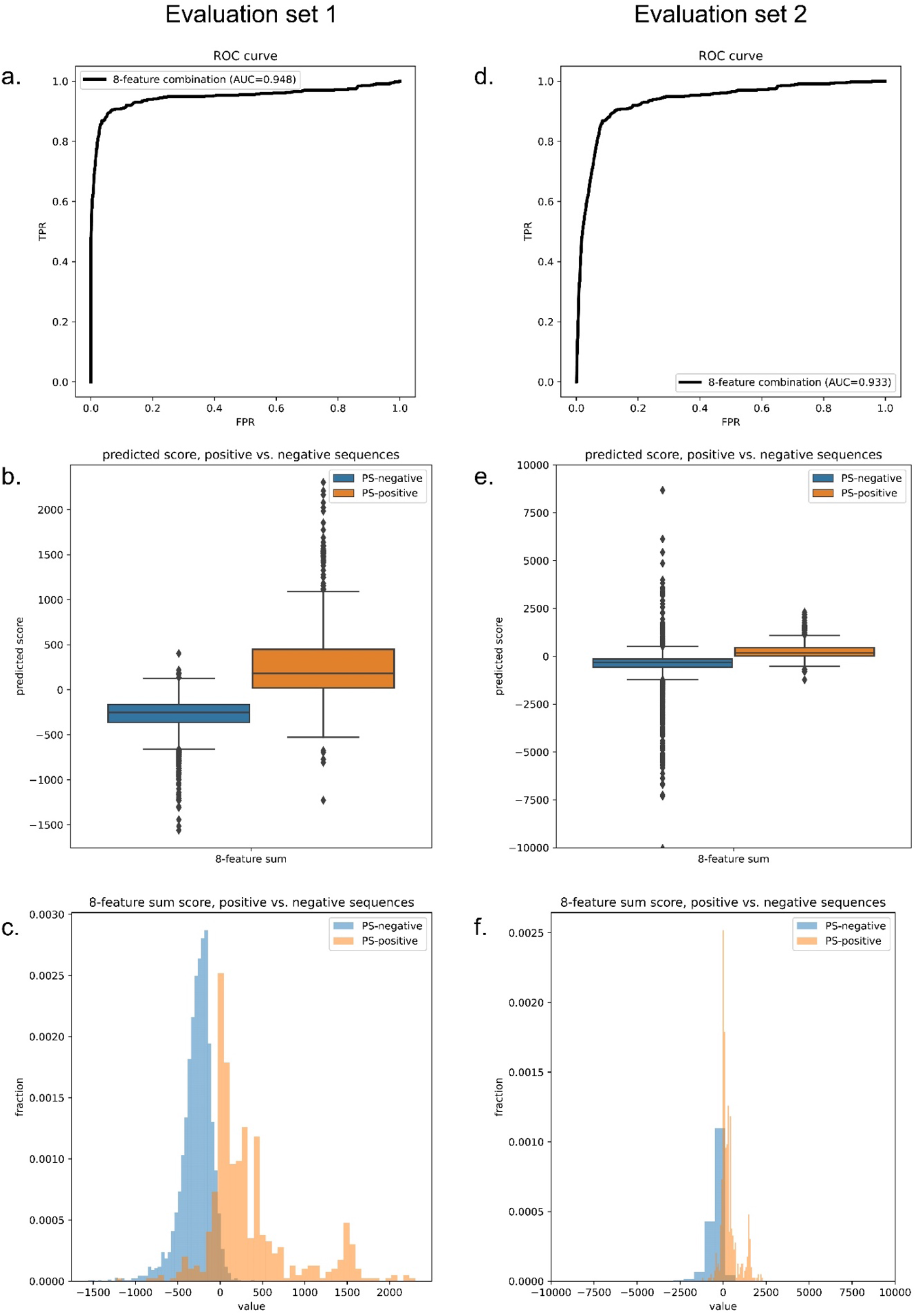
Final predictor model performance. Performance plots of the final Human+PDB model on Evaluation set 1 (left) and Evaluation set 2 (right). (a,d) ROC curves. (b,e) Predicted score boxplots of positive vs. negative sequences. (c,f) Distribution histograms of positive vs. negative sequences.

### Predictor validation

To validate the final predictor’s performance, we constructed three sets of baselines. (1) Instead of providing PDB-based physical feature values into the genetic algorithm, we provided random values from a normal distribution N(0, 1) in the weight training step. (2) Instead of providing sequence-based physical feature values, we provided random values from the distribution of residue-specific physical feature values. (3) Instead of optimizing 20 weights for 20 residue types for each physical feature, we optimized 1 weight for all 20 residue types for each physical feature (removing residue specificity) during training. As shown in Figure 7 for the Human+PDB model and Supplementary Figure S6 for all three models, baselines 1 and 2 showed very high training AUROC but low test AUROC, whereas the final models have both high training and test AUROC. This suggests that the final predictors’ good performances did not come from overtraining by the genetic algorithm, which is the case for baselines 1 and 2. The comparison between baseline 3 and final models also suggests that it is important to have residue specificity in our model for good prediction performance.

**Figure 7.**
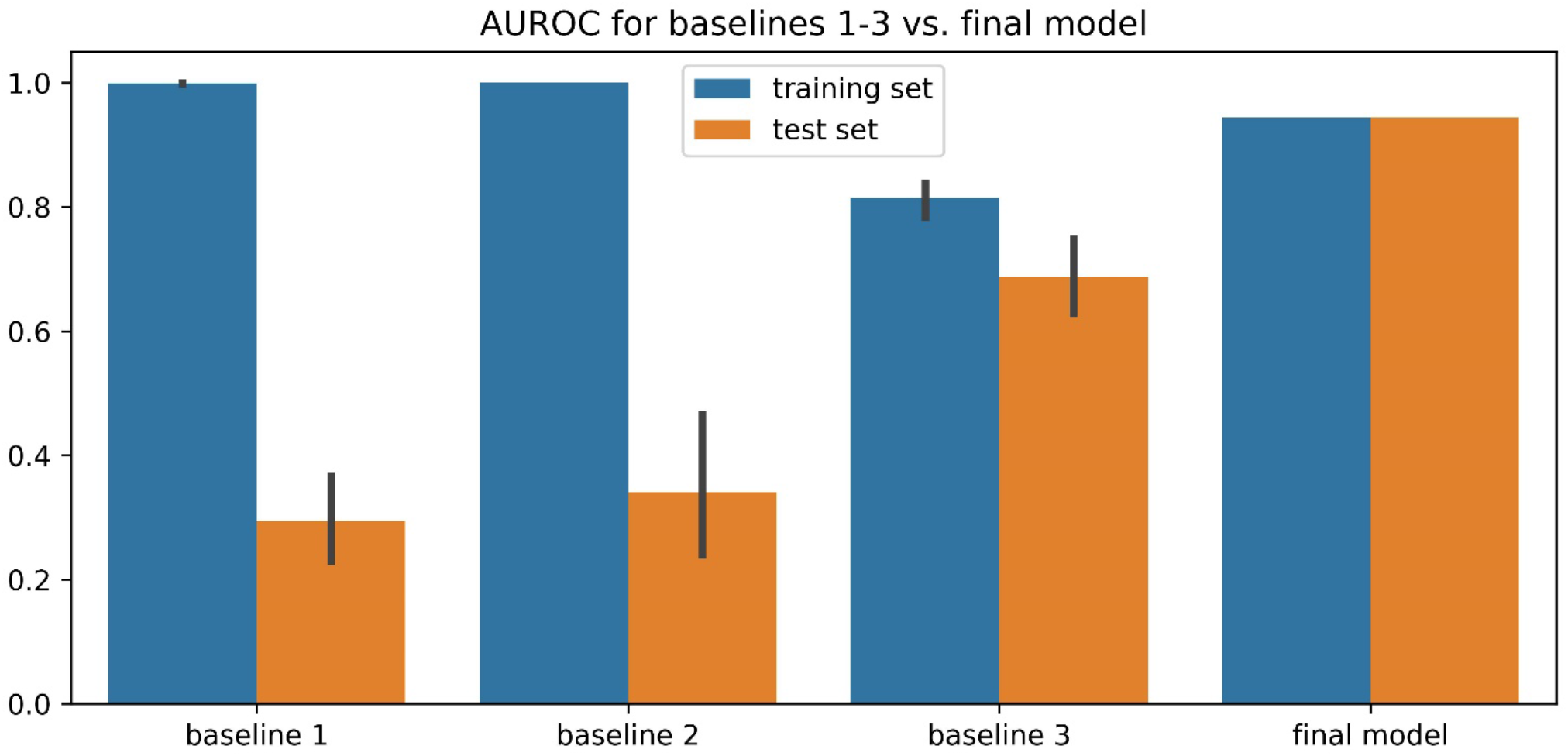
Comparison of three training baselines and the final Human+PDB predictor model for validation. Baseline 1 was created by providing random values from a normal distribution N(0, 1) in the weight training step instead of providing PDB-based physical feature values into the genetic algorithm. Baseline 2 was created by providing random values from the distribution of residue-specific physical feature values instead of providing sequence-based physical feature values. Baseline 3 was created by optimizing 1 weight for 20 residue types for each physical feature (removing residue specificity) during training instead of optimizing 20 weights for 20 residue types for each physical feature.

### Comparison of prediction using 8 features or single features

To test whether a combination of 8 features can outperform prediction by a single feature, we extracted from the three final models each of the feature components as 1-feature predictors, and evaluated these 1-feature predictors on Evaluation set 1. As shown in Figure 8a and Supplementary Figure S7, receiver operating curves (ROCs) of 1-feature predictors were outperformed by 8-feature predictors. We also plotted Venn diagrams showing their recalled sequences at a confidence threshold that returns 2% of the PDB as positive result (chosen based on the methods described in previous work [32, 38, 40]) as shown in Figure 8b and Supplementary Figure S8. We observed that each of the 1-feature predictors missed a number of sequences (48-350 sequences) that were captured by the 8-feature models. This result supports our underlying assumption that phase separation is driven by a combination of different physical features, and driving forces for different sequences can vary.

**Figure 8.**
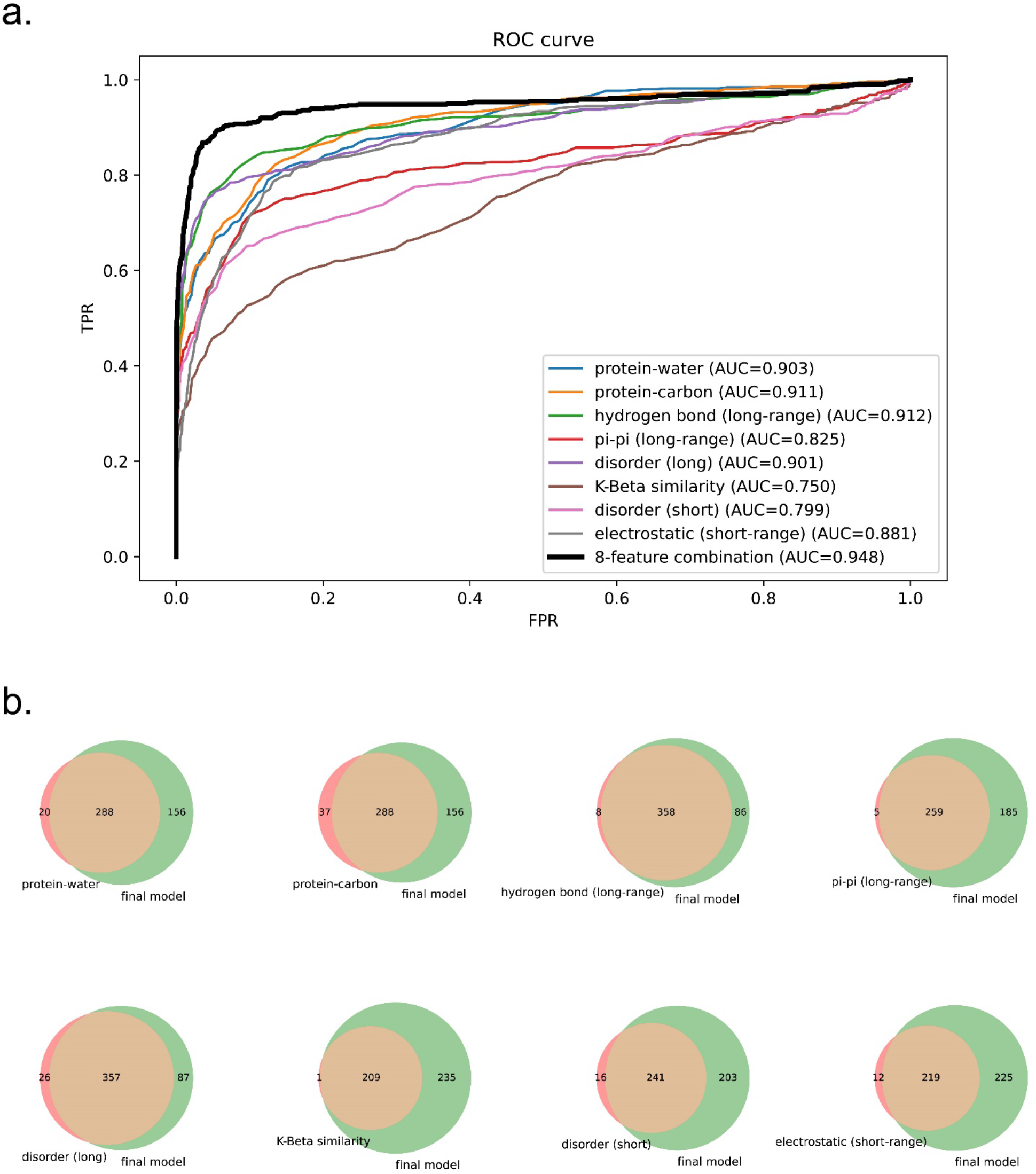
Comparison of performance of predictors trained on 8 features vs. 1 feature for the Human+PDB model. (a) ROC curves of 1-feature predictors vs. 8-feature predictor. (b) Venn diagrams showing the coverage overlaps of PS-positive sequences by 1-feature predictors vs. the 8-feature predictor at a confidence threshold that returns 2% of the PDB.

### Comparison between LLPhyScore and other phase separation predictors

We compared the performance of our predictor (LLPhyScore, three final models) with PSPredictor, as well as two first-generation predictors, PScore and catGRANULE in Figure 9. The comparison was conducted on both Evaluation set 1 (against PDB) and Evaluation set 2 (against human proteome).

**Figure 9.**
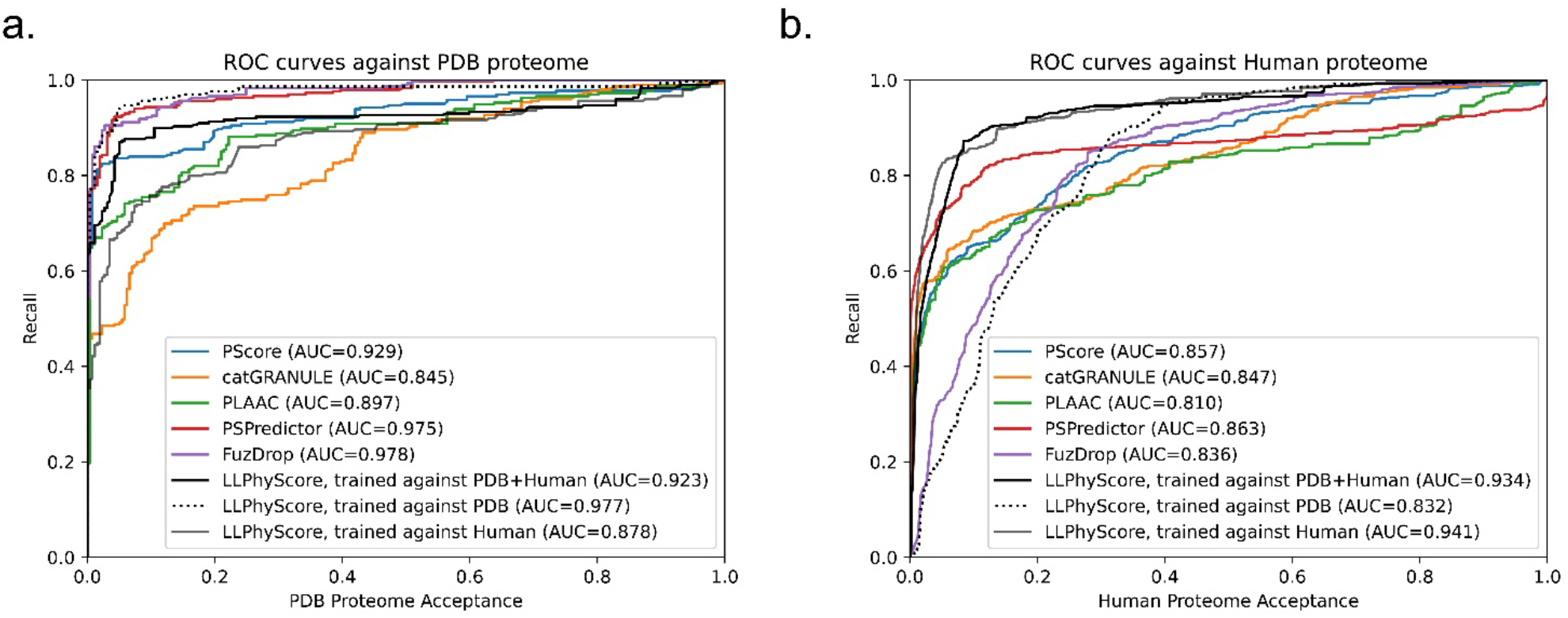
Comparison of LLPhyScore (3 models) with other phase separation predictors. Relationship between percent recall and total percentage of (a) PDB proteins and (b) human proteins accepted at given thresholds for PScore, catGRANULE, PLAAC, PSPredictor, FuzDrop and LLPhyScore.

We can see that the LLPhyScore-PDB model showed the best performance on Evaluation set 1 and even slightly outperformed PSPredictor, which was trained against 5258 sequences from the PDB. The LLPhyScore-PDB model also showed a better AUROC than PScore, which is based solely on planar pi-pi interactions. The LLPhyScore-Human+PDB model showed slightly decreased performance on Evaluation set 1 compared to the LLPhyScore-PDB model, however was still better than all first-generation predictors. The LLPhyScore-Human model showed a comparable performance as PLAAC.

However, on Evaluation set 2, the LLPhyScore-PDB model didn’t show better recall statistics than other first-generation predictors until a 30% acceptance threshold, as shown in Figure 9b. This is in line with the estimate of up to 40% of the human proteome driving phase separation[48], and could be considered support for an estimate of at least 30% of the proteome being involved in phase separation. On the other hand, the LLPhyScore-Human model and LLPhyScore-Human+PDB model both showed good performance on Evaluation set 2, indicating that, by mixing Human and PDB sequences, the training algorithm can optimize PS-positive sequences from both negative sets. We also see (Figure 9a,b) that the LLPhyScore-PDB model showed comparable recall trends with FuzDrop. As a phase separation predictor also based on biophysical principles combined with statistical training, FuzDrop uses a protein’s binding entropy as the target function. The fact that the LLPhyScore-PDB model and FuzDrop showed similar statistics supports the utility of approaches directly addressing the biophysical features and energetic driving forces underlying the formation of condensates.

### Feature-based breakdown of scores for different sequences

To further explore the general expectation that phase separation of different sequences can be driven by different physical features, we clustered PS-positive sequences based on their single feature-scores after normalization. As shown in Figure 10 and Supplementary Figures S9 and S10, FUS, Nup98, an elastin-like peptide (ELP), and MEG-3 were categorized into different clusters, which demonstrates the ability of LLPhyScore to treat different types of sequences. For the LLPhyScore-Human+PDB model, the breakdown of scores (Figure 10) shows that Nup98 has high scores for protein-carbon but low scores of disorder, pi-pi, and K-beta, whereas for FUS, the scores are high for most of the features.

**Figure 10.**
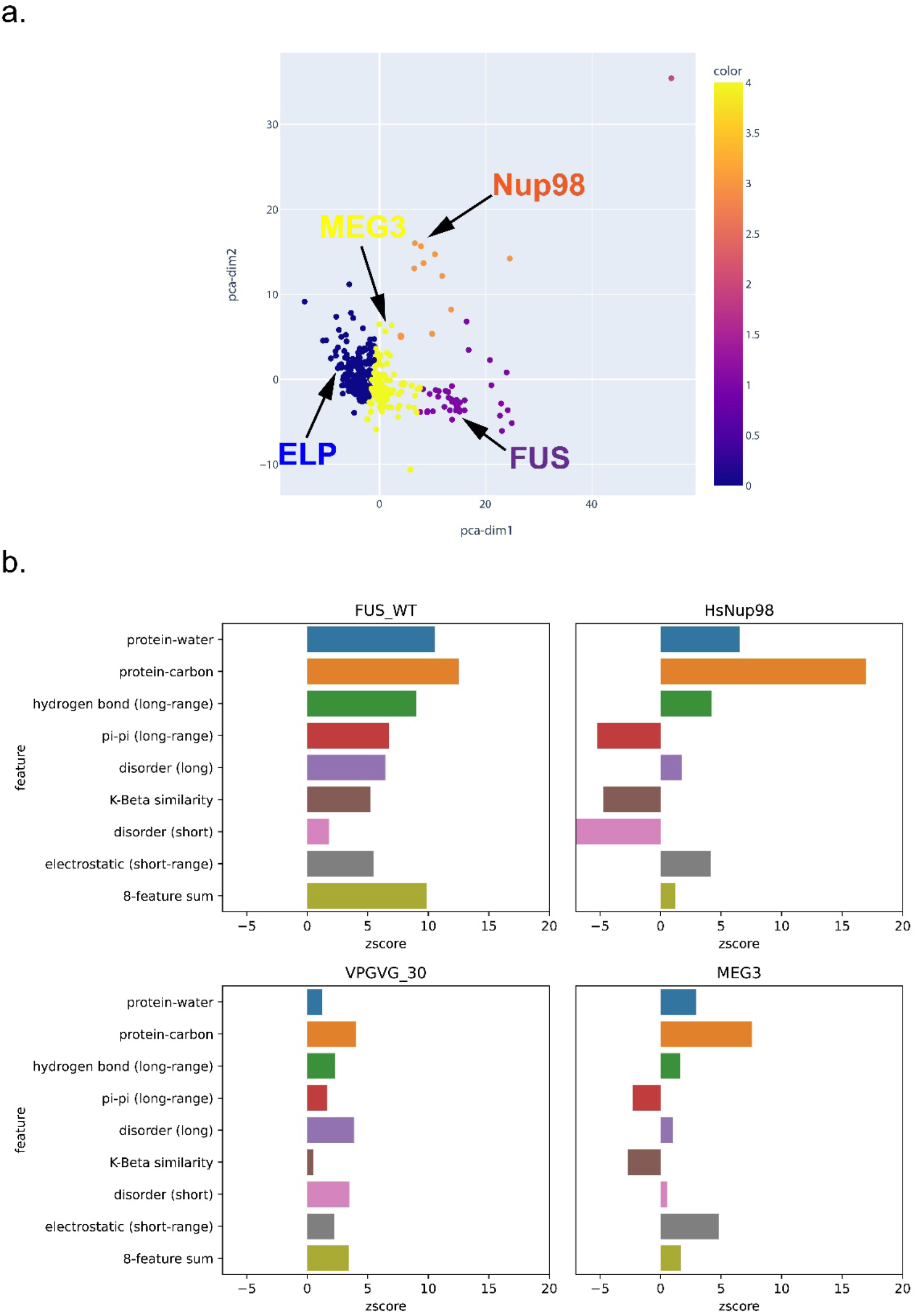
Feature score-based clustering for PS-positive proteins for the Human+PDB model. (a) Plot of 2 abstracted dimensions for clustering based on feature z-scores, showing the separation of different types of phase-separating sequences. (b) The score breakdown of 4 example sequences from 4 distinct clusters in (a): FUS (human), Nup98 (human), elastin-like peptide (ELP, VPGVG_30, 30 repeats of VPGVG) and MEG-3 (*C. elegans*).

### Gene Ontology term enrichment

We analyzed the enrichment of GO terms for human proteins in the top 2% (high confidential threshold) of scores from the LLPhyScore models for the whole human proteome predicted by 8 single features as well as the combination of 8 features in the final predictor. As shown in Figure 11, Supplementary Figure S11 and **Supplementary Table S5**, most GO terms identified by first-generation predictors [38] and by PSPredictor [40] were also enriched in sequences identified by LLPhyScore, such as extracellular matrix and nuclear body. For certain annotations associated with phase separation such as cytoplasmic stress granule, postsynaptic density and transcription factor complex, we observed significant differences depending on which features were utilized, which suggests that, for different biomolecular condensates with different functional roles for phase separation, the key features linked to phase separation are also different, and are rooted in their sequence-specific biophysical landscape.

**Figure 11.**
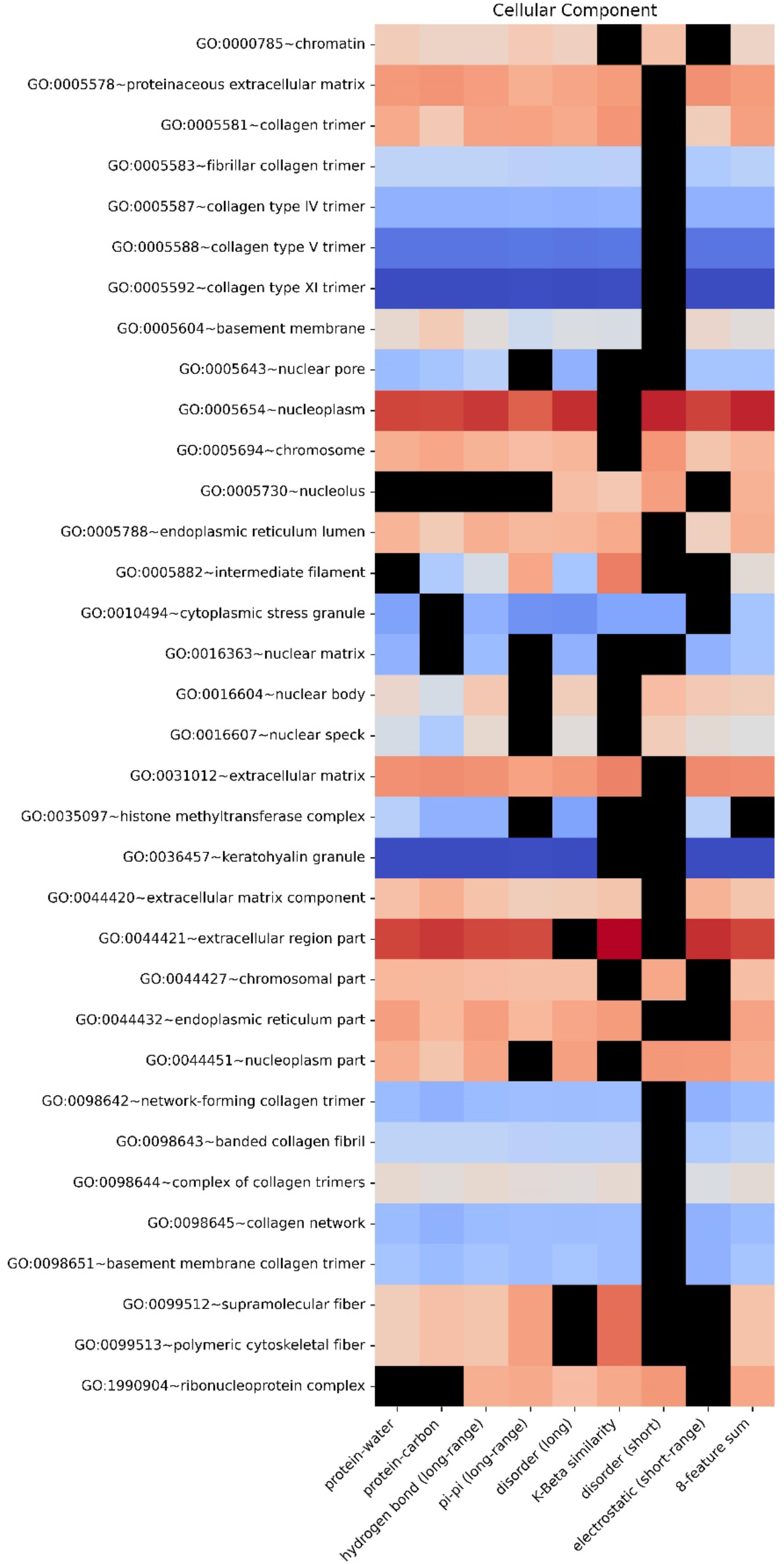
Enrichment heatmap by GO functional annotations for different features for the human+PDB model. Heatmap showing the enrichment of proteins with a given functional annotation that fall under a 2% confidence threshold for each single feature score and 8-feature sum score. The color gradient shows the natural logarithm of the enrichment percentage.

### Physical insights into phase separation based on LLPhyScores of the PDB set

Assessment of the physical basis of LLPhyScore predictions is complicated by detailed choices made during the training process, where the weight given to a feature by the final model not only reflects that feature but reflects the full sequence context of the residue. Therefore, to assess how scores relate to the physical features for which we trained, we applied the predictor to the sequences of known structures in order to assess phase separation scores by comparing directly to “true” measurements of sequences in observed structural contexts. For this, we scored each amino acid independently, comparing the physical features associated with being in the top 50% of scores against the overall distribution for that residue type. Figure 12 shows high-score enrichment statistics for a variety of physical features, including secondary structure (a), short-range pi-pi (b), kinked-beta similarity and dissimilarity (c), disorder (d), short- and long-range electrostatics (d,e), and local water/carbon contacts (f,g).

**Figure 12.**
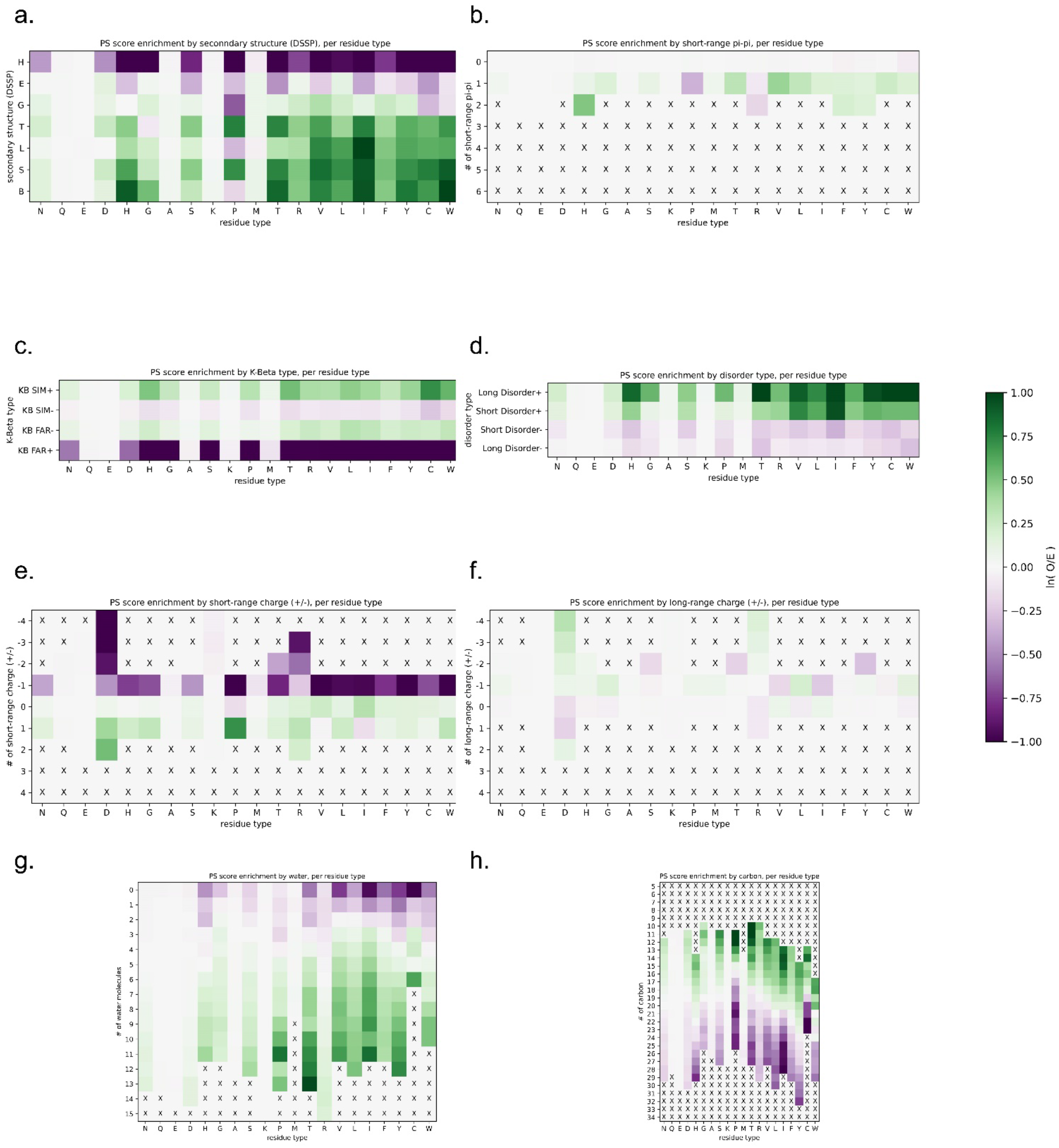
LLPhyScore score enrichment by 8 selected physical features for the PDB proteome, per residue type, for the human+PDB model. Heatmaps show the score enrichment in PDB protein sequences by each feature’s discrete values, normalized to each residue type. The color gradient shows the natural logarithm of the observed over expected ratio. Enrichment for (a) secondary structure (H, alpha helix; E, beta sheet; G, 3_10_ helix; T, hydrogen bonded turn; L, loop; S, bend; B, single pair beta-sheet), (b) short-range pi-pi, (c) K-beta, (d) disorder, (e) short-range electrostatic, (f) long-range electrostatic, (g) protein-water and (h) protein-carbon. The color-bar for all heatmaps is shown at the right.

For protein sequences found in our PDB set the predictor generally gives low scores to structures that can satisfy their interactions locally. Helical residues which fully satisfy their backbone hydrogen bonds typically have low scores (Figure 12a), as do residues with stabilizing charge interactions found between nearby local residues (Figure 12e). Notably, charge interactions between non-local residues (Figure 12f) have above average scores, consistent with the known effects of blocks of like-charge in driving phase separation.[26, 65, 66]

For secondary structure, there appear to be three categories of effects based on backbone hydrogen-bonding satisfaction and torsion angle regularity. Fully self-satisfied structures, specifically helices, have the lowest scores. Ordered but not necessarily locally-satisfied, which includes beta-strands as well as 3_10_ helices (often associated with short helices[67, 68] and capping motifs[69]), have intermediate scores. Irregular secondary structure, including elements with defined hydrogen-bonding patterns (turns, bulges, and bent/kinked strands), as well as solvent-bound loops, have the highest scores. In general, the ability to form hydrogen-bonds to solvent is consistently associated with higher scores, as is a lack of repetitive ordered structure. In this analysis, bent and twisted strands typically score better than fully disordered residues, especially for proline, suggesting that the availability of backbone hydrogen bonding plays a role, and not just the lack of structure.

The differences between disorder prediction and phase separation prediction is further defined in Figure 12d. In general, disordered residues are more likely to score high, with long stretches of disorder scoring higher than short disordered loops. However, the majority of this bias comes from hydrophobic or aromatic residues, specifically V, L, I, F, H, Y, and W. This is consistent with disorder on its own being insufficient for phase separation, with disorder that forces hydrophobic and aromatic residues into contact with solvent supporting phase separation.

This solvent relationship can also be observed directly by measurement of solvent interactions and overall burial, as shown in Figure 12g,e. In general, residues with a high number of observed water contacts have higher scores, and residues with a high degree of burial (assessed by the number of carbon contacts) have lower scores. However, this trend is more pronounced for hydrophobic residues and is not observed for polar or negative residues (N, Q, E and D). This may be expected given that the hydrophobic effect is driven by solvent, with the energy associated with a reduction of solvation driving hydrophobic residues together (i.e., solvent relationships are what makes hydrophobics sticky). In this context, we observe that hydrophobic residues forced to be in contact with solvent by their local sequence context are predicted to contribute to phase separation.

The notion that sequences which force solvation are prone to phase separation matches observations for secondary structure. We note that while extended beta-sheets can often exclude solvent, by forming flat planar interactions with other sheets, kinked beta-strands cannot. Figure 12c shows that sequences with high structural similarity to kinked-beta have higher scores, especially for hydrophobics and aromatics.

Together, our analyses of the LLPhyScores for PDB structures supports the view that disorder itself does not drive phase separation, but locally unsatisfied sequences that are constrained in their ability to exclude solvent, including those that can adopt irregular or kinked-beta structure to contribute backbone hydrogen bonds, do drive phase separation. These results may contribute to the current discussion of the role of sequences with propensity to form kinked-beta structure in protein phase separation.[33, 63]

### High scoring structures in the PDB trend towards disorder

The protein structure databank is often used as a negative set when training phase separation classifiers but this is not a ground truth, and the true fraction of proteins with structures present in the PDB that are also capable of driving phase separation is unknown. To demonstrate this issue we scored the set of PDB reference sequences used in this study and observe that the highest scoring proteins are not random; the score selects for proteins with significant disorder relative to the average structure found within the PDB (Figure 13). This is true using multiple definitions of disorder: (i) of the highest scoring 1% (N=167), 99 have more than 50% of the reference sequence missing from density (Figure 13a), and (ii) for the residues which are found within the density (Figure 13b), 128 of these proteins have more than 50% of the residues in secondary structure classes other than helix and strand, with 62 of these having more than 50% of their residues in contiguous loop/turn/random coil segments of 4 or more residues in length.

**Figure 13.**
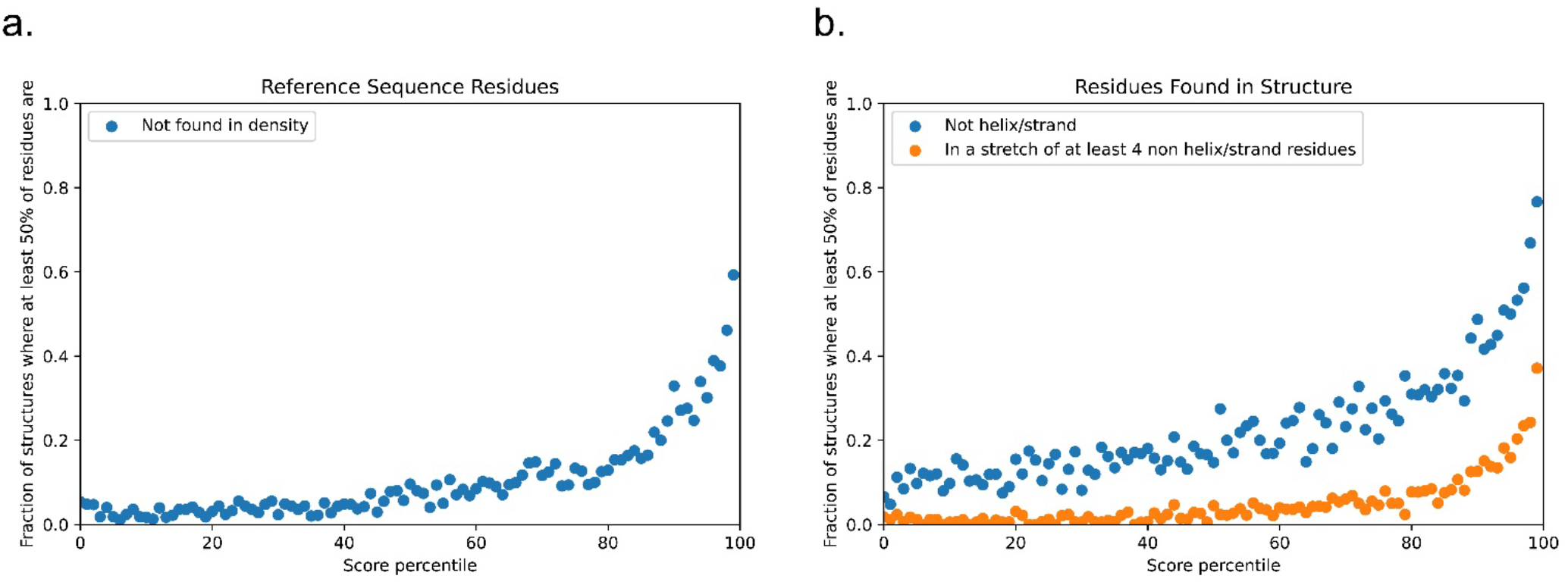
Disordered character of PDB sequences according to the LLPhyScore of chain reference sequences. Panel (a) shows the fraction of proteins in each percentile bin of LLPPhyScore for which more than 50% of the reference sequence is missing from density (protein sequence that does not show up in the structure). Panel (b) shows the disordered/irregular structural character of residues that are within density in the structure, with blue showing the fraction of proteins in each percentile bin for which more than 50% of the observed residues have a DSSP assignment other than helix or strand, and orange shows the fraction for which more than 50% of such residues are found in stretches of at least 4 residues in length with no helical or sheet structure.

The highest scoring sequences for LLPhyScore in the PDB depart significantly from an expectation of well-ordered folded domains, and their function is unlikely to be defined simply by their ability to form the state observed within these structures. Future work in training classifiers would benefit from a biophysically defined empirical negative set.

## Conclusion

In this work, we demonstrated the utility of combining different physicochemical interactions as driving forces in the prediction of protein phase separation. We addressed the issue of “imperfect negative training set” by training three predictor models on three different negative sets and compared their performance. We optimized the combination of physical features in the final predictor models and achieved a superior performance over first-generation predictors. Importantly, our predictors enable physical interpretability not possible with another comprehensive predictor, PSPredictor. Our results are consistent with the understanding that phase separation is driven by a combination of physical factors, including pi-pi contacts, disorder, hydrogen bonding, including in the context of kinked-beta structure, electrostatics, and water interactions. By clustering sequences based on their physical-feature scores, we can differentiate phase-separating sequences by their contributing driving forces, suggesting one contributor to the basis for specificity in the formation of the large number of unique biomolecular condensates found in biology. LLPhyScore should be a useful tool for the protein phase separation field to provide hypotheses regarding key interactions driving phase separation, as well as for screening proteins that may play important biological roles in the context of biomolecular condensates.

## Methods

### Curation of PS-positive sequences

We did a search on PubMed for all papers published from Jul 2013 to Jan 2019 that contains keywords related to phase separation (“phase separation”, “liquid condensates”, “membraneless organelles”, etc.), and manually screened out 142 papers out of 689 articles that describe *in vitro* phase-separating systems. Then, we extracted all sequences from the papers (main content/supplementary information/uniprot entry) that had clear evidence of phase separation on their own (either detailed phase diagram, or mentioned as “phase separation positive” in the text) in the content. A total of 565 sequences were extracted and were checked twice.

### Clustering of PS-positive sequences

The clustering of positive sequences was done by hierarchical clustering (shown in Supplementary Figures S1 and S2). First, a 20×20 dipeptide count number grid was calculated for each sequence, with each number being the number of a residue pair (e.g. AG) in the sequence (**Equation 1**). Then, a Jaccard similarity value was calculated for any two sequences by dividing the overlap by union of two 20×20 grids (**Equation 2**). If two sequences have different lengths, then a sliding window of the smaller length will be applied to the longer sequence, and the highest similarity value calculated for all sliding windows will be kept. Finally, we used the hierarchical clustering package in Python scikit-learn[70] to conduct the clustering for 565 sequences. A cutoff similarity threshold of 0.5 was chosen.

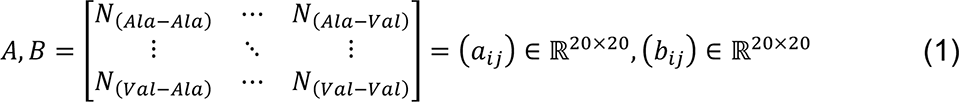

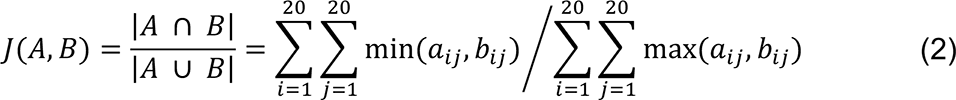

### Preparation of PS-negative sequences

Two PS-negative sequence databases were prepared. First, 16794 sequences were collected from high-resolution (≤2.0 A) structures in PDB as the first negative sequence database – “PDB base”; Second, 20380 human proteome sequences were collected from Uniprot[71], then we used their CRAPome scores calculated in Vernon et al.[38] as a filter for PS-positive sequences. Sequences with either null values or high values (top 20%) in CRAPome were removed, resulting in a “clean” human set of 6102 sequences – “Human base” (**Supplementary Table S6**).

From these two PS-negative sequence databases, three negative sets were prepared: (a) PDB set, including 3406 sequences randomly selected from PDB base; (b) Human set, including 3406 sequences randomly selected from Human base; (c) (Human+PDB) set, including 1703 sequences randomly selected from PDB set and 1703 sequences randomly selected from Human set.

### Construction of Training/Test/Evaluation Datasets

The construction of training set and test set starts with PS-positive sequences. Random sampling was conducted on 565 PS-positive sequences at clustered group level, with 305 sequences went to training set and 260 sequences went to test set. Then, a 50%-50% split ratio was applied on three PS-negative sets at sequence level, with 1703 sequences from each set went to training set, and another 1703 sequences went to test set. A total of three training-test sets pairs were constructed accordingly.

Two evaluation sets were constructed. (1) Entire PS-positive set (565 sequences) + entire PDB base (16794); (2) Entire PS-positive set (565 sequences) + entire Human base (20380 sequences).

For more details, see Supplementary Table S1.

### Physical feature-based sequence representation

Eight different general phase separation-driving factors were taken into account to represent a protein sequence, in total makes 16 physical features, as summarized in Supplementary Table S2. For each of these features, its sequence-based statistics (contact frequency/number of atoms/structure probability) in the PDB were acquired by mining the structures of folded proteins in PDB. The observations were split by distinct residue pairs with varying sequence separations, leading to a database of “feature values”, with each “feature value” being an empirical, per-amino acid energy potential corresponding to the frequencies of specific contact types in the PDB. Then, for a given input sequence, inferred “feature values” for each residue of this sequence were obtained by matching its residue pair and sequence context to the “feature value database”. For example, the short-range pi-pi contact frequency for valine in tripeptide valine-glycine-tryptophan can be inferred by taking the average short-range pi-pi contact frequency for residue pair valine-glycine with 0 separation and valine-tryptophan with 1-residue separation (see also Figure 2).

Specific definitions for each of these were as follows:

#### Pi-Pi Contacts

Pi-Pi contacts were defined using the method in Vernon et al.[29], and then divided into short range and long range by sequence separation. Less than five residues apart is defined as short range, and greater than or equal to 5 residues apart is defined as long range.

#### Hydrogen Bonding Terms

Structures were probed for hydrogen OH-N hydrogen bonds using PHENIX[61], with the following commands used to extract hydrogen bond information. phenix.reduce -Quiet -FLIP [pdb file] > ./PHENIX_ALL/PHENIXL.pdb phenix.probe “NITROGEN,OXYGEN,HYDROGEN” -Quiet -ONEDOTeach -NOCLASHOUT - SUMMARY -NOVDWOUT ./PHENIX_ALL/PHENIXL.pdb | grep greentint > ./N17.PHENIX/HLIST.’+pdb

Bonds were than classified as short range and long range by sequence separation (short range < 5, long range ≥ 5).

#### Water/Carbon Contact Counts

Water and carbon counts were calculated only for the subset of proteins in our training set that have a total number of water molecules greater than the number of protein residues. This captures almost all models with resolution <= 1.8 but removes lower resolution models. Counts are measured for residues in their crystallographic context (measurement includes atoms from symmetry partners).

#### Secondary Structure

DSSP letter code was used for secondary structure assignments, with H/G used for helix, E for strand, and all others binned to loop.

#### Disorder

For identifying disordered residues a DSSP assignment of “not G/H/E” over a span of at least 3 residues was used to classify residues as loops. These loop residues were then assigned as short disorder if they fall within 3 residues of G/H/E and as long disorder if they do not.

#### Charge

PHENIX (via the phenix.reduce command) was used to complete PDB structures by adding hydrogen atoms and charge interactions were calculated using the following pseudocode, with partial charges taken from the Talaris energy function[59].

q1 = partial_charge for atom X of amino acid 1

q2 = partial_charge for atom Y of amino acid 2

absF = 330.0 * abs(q1*q2)/(distance**2)

if q1*q2 < 0.0: absF *= −1.0

if SequenceSeparation >= 10: add absF to electrostatic (long range)

if SequenceSeparation < 10: add absF to electrostatic (short range)

Final per-residue values were then binned as follows:

bin = np.clip(int( round( residue_value / 16.0 ) ), −9, 9)

#### Cation-Pi

We recalculated the electrostatic scores after adding arbitrary partial charges to the surfaces of aromatic rings, with a partial charge value of −0.05 added 0.85 Å above and below the plane of the ring for each atom, counterbalanced by a partial charge of 0.1 at the atom. The cation pi-score is then taken from the difference between this modified score and the unmodified electrostatic score.

#### Kinked Beta

Superpositions to kinked beta fibrils were made for chain A in each of 5 structures, PDB IDs 6bwz, 6bxv, 6bxx, 6bzm, and 6bzp. The full chain of each was superimposed to every overlapping window (same number of residues as the chain with none missing) in our PDB training set and kinked beta similarity was measured for each individual PDB residue by taking the minimum CA-RMSD over all measurements the residue was involved in. Residues were then classified as K-Beta similar if the minimum CA-RMSD was under 1.0 Å and as K-Beta dissimilar if it was over 2.0 Å.

These 16 physical features are converted to an inferred feature statistics value for every sequence with representation at the residue level and sequence level. At the residue level, each amino acid is represented by 16×3 numbers describing the impact of each of the 16 biophysical forces on each residue: (1) the amino acid position number, (2) the score from comparing to the upper feature value threshold (W_U_) and (3) the score from comparing to lower feature value threshold (W_L_).

Inferred feature statistics for a protein sequence is based on 16×20×N matrices, based on three components in the sequence representation, which function as 3 layers of our machine learning model architecture. (i) A sequence is characterized by 16 physical features acting on each residue. (ii) The impact of each physical feature is dependent on residue type, represented by 20 residue-type groups. (iii) N is the number of residues of a specific type within the sequence, with *z* being the position (or index, see below).

Thus, inferred feature statistical values are determined by translating protein sequences into 16×20×N matrices (See **Equation 3** and Figure 2).

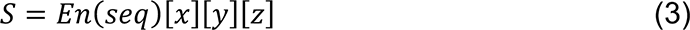

Where

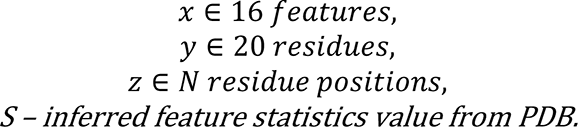

### Predictor training

Predictor training has the following steps: (1) For each physical feature and each residue type, we set a upper and lower threshold (“weight”) for its inferred feature value, thereby constructing a 16×20×2 (each feature has two weight values - upper and lower threshold) array (**Equation 4**). (2) We initialize the “sum feature score” for each physical score to 0. (3) For each residue in a sequence, if its feature score is higher than the upper threshold, we consider this residue as “abnormally active” in this physical feature, and reward the corresponding “sum feature score” by adding 1 to it; if its feature score is lower than the lower threshold, we consider it as “abnormally inactive” in this physical feature, and penalize the corresponding “sum feature score” by subtracting 1 from it; if its feature score is between the upper and lower threshold, we consider it to be “within normal range”, and do nothing (**Equations 5 and 6**). (4) By optimizing the AUROC score function (**Equation 7**) for each feature, we found the best feature combination, and the best weight that maximize the gap of sum feature score(s) between PS-positive sequences and PS-negative sequences (**Equation 8**). (5) By summing “sum feature scores” and training the weights of features using genetic algorithm, we calculate a “total sum probability” for any sequence, which is the final estimate of its phase separation ability (Equation 9).

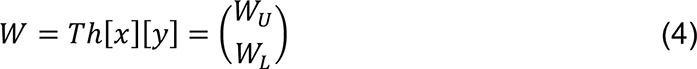

Where

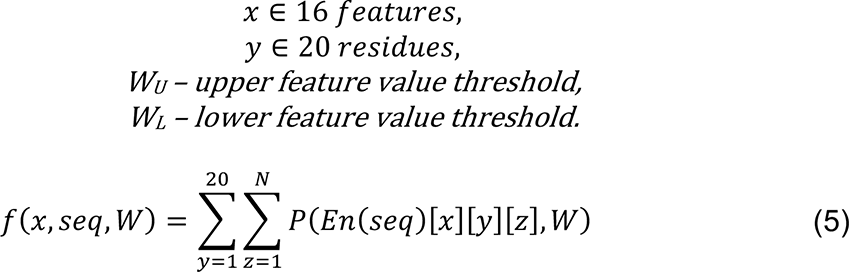

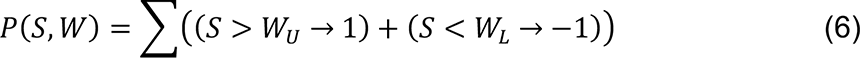

Where

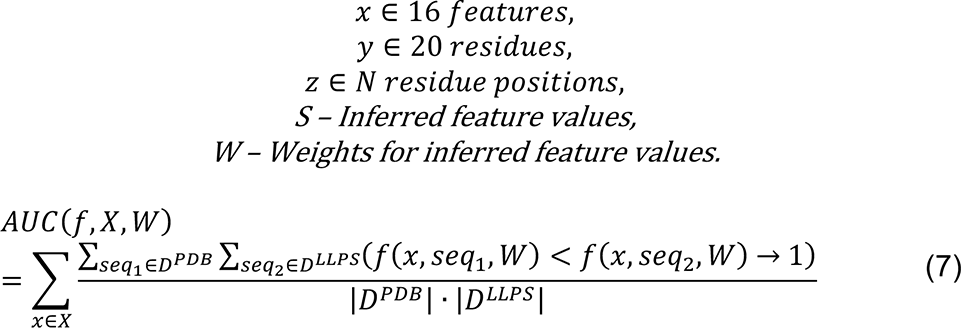

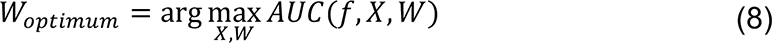

Where

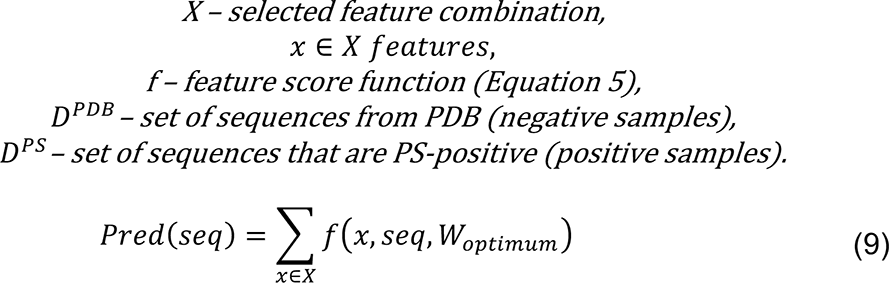

Where

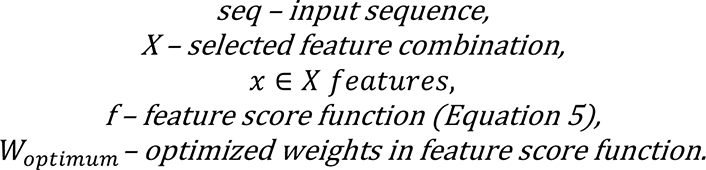

The optimization process for parameters in a predictive algorithm is called “training”. In this work, training of the phase separation predictor has two parts: (i) training of the upper and lower weights of “binary feature score” for 16 features x 20 residue types; (ii) training of the combination of features to include in the final predictor. Numerically, the number of parameters trained is 16×20×2 weights = 640 weights (16 biophysical forces; 20 residue types; two weights, W_U_ (upper threshold) and W_L_ (lower threshold). Another “hyperparameter” being trained here is the selection of biophysical forces to include, with only 8 out of the 16 biophysical forces ultimately used to avoid overfitting (requiring only 320 weights). The data used for training initially is from the sequences of the PS-positive proteins in the training set split off from the test set (565 – 260 sequences) and the PS-negative sequences (1703 from either the PDB, Human or Human+PDB). The data used for training the “final models” used all 565 sequences of the PS-positive proteins and 3406 sequences from either the PDB, human, or PDB+human PS-negative sets.

This training was conducted on the positive and negative training datasets using a genetic algorithm. Specifically, we randomly generated an initial set of 640 weights, and then, for each iteration, we randomly picked a subset of these 640 weights to change and accepted the changes that improved the behavior (loss function based on “genetic operators”). We performed many iterations until a fixed number of generations were reached. The loss function is the AUROC curve (area under the receiver operating characteristic curve) as described above (**Equation 7**); the performance of the predictor was then evaluated using the test set as well as by comparison against baseline models.

Importantly, we used a genetic algorithm to optimize weights (parameters) with the overall architecture being a 3-layer “neural network”-like predictive model with a non-convex loss function. For more details on implementation of training and prediction, please see https://github.com/julie-forman-kay-lab/LLPhyScore.

### Proteome analysis

Human proteins with scores in the top 2% of human proteome using 8 predicted single-feature scores as well as the final predictor (8-feature sum score) were uploaded separately to DAVID 6.7 (https://david-d.ncifcrf.gov/)[72]. The enrichments of biological process, cellular component, and molecular function GO terms were analyzed for the proteins with respective p-values (EASE score) obtained. The resulting GO term enrichments were compared against the results in Vernon et al.[29], Vernon et al.[38], and Chu et al.[40].

## Supporting information

Supplementary_Figures_jpg

## Supplementary Materials/Data and Code Availability

This article contains supplementary material. Supplementary Figures S1-S11 and Supplementary Tables S1-S4 are included below, following the main text references. Additional Supplementary Tables and Files are provided as separate files:

One attached Excel file contains, on separate tabs, **Tables S5-S10.**

**Table S5**. GO enrichment analysis.

**Table S6**. Uniprot IDs of 6102 sequences from human proteome using CRAPome as filtering method.

**Table S7**. Uniprot IDs of 3406 sequences from PDB base.

**Table S8**. Uniprot IDs of 3406 sequences randomly selected from human base in Table S6.

**Table S9**. Uniprot IDs of 6812 sequences from PDB+human base.

**Table S10**. Detailed information of 565 PS-positive sequences with PMID of each sequence’s paper.

**File S1**. 565 PS-positive sequences (fasta file).

**File S2**. 16794 PDB sequences (fasta file).

**File S3**. 20380 Human sequences (fasta file).

The software for running LLPhyScore and more details on the training are provided in the following GitHub: https://github.com/julie-forman-kay-lab/LLPhyScore.

## Acknowledgements

We sincerely thank Zhuqing Zhang and Jianfeng Pei for providing PSPredictor scores and Monika Fuxreiter and Michele Vendruscolo for providing FuzDrop scores, in both cases for the human proteome and PDB sequences. The authors acknowledge Alan Moses and Andrew Chong for critical reading of the manuscript.

## Author Contributions

Conceptualization, R.M.V., H.C., J.D.F.-K.; Methodology, R.M.V., H.C.; Software, H.C.; Validation, R.M.V., H.C.; Formal Analysis, R.M.V., H.C.; Investigation, H.C., R.M.V.; Resources, J.D.F.-K.; Data Curation, H.C.; Writing – Original Draft Preparation, H.C., R.M.V.; Writing – Review & Editing, J.D.F.-K., R.M.V., H.C.; Visualization, H.C., R.M.V.; Supervision, R.M.V., J.D.F.-K.; Project Administration, J.D.F.-K.; Funding Acquisition, J.D.F.-K.

## Supplementary Material

**Figure S1.**
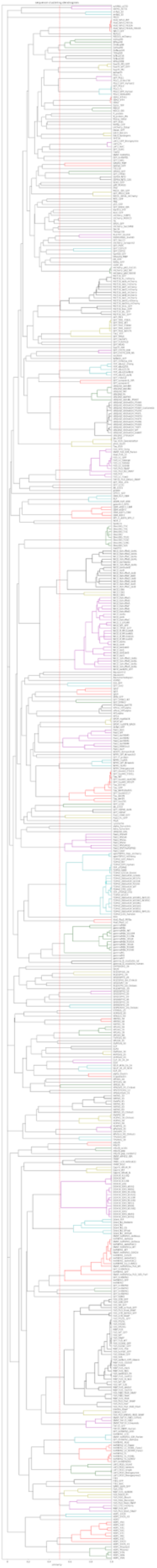
Hierarchical clustering dendrogram of PS-positive sequences.

**Figure S2.**
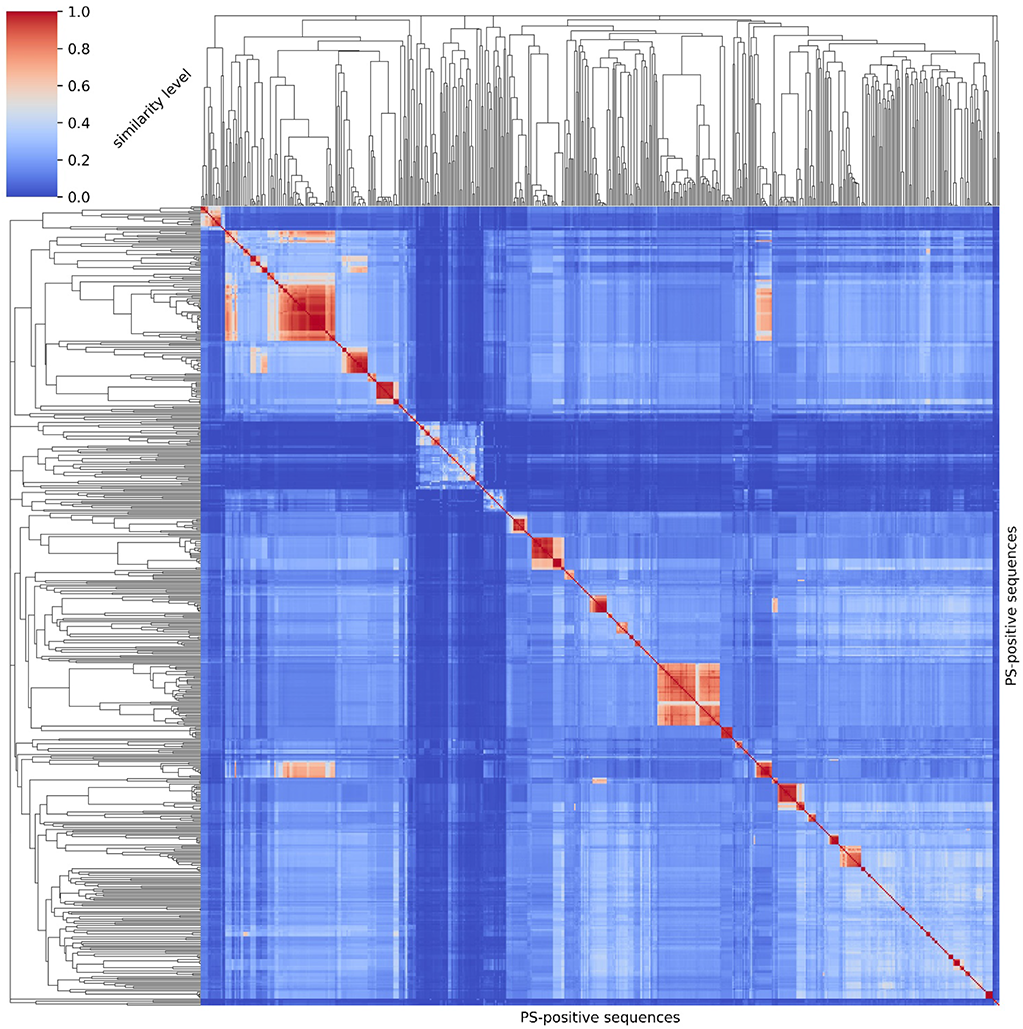
Sequence clustering map of PS-positive sequences. Left and top: clustering dendrograms showing the pairwise similarity level (0 - 1.0) for all sequences. Middle: heatmap of similarity levels showing that inter-cluster sequences have pairwise similarity higher than 0.5 in general.

**Figure S3.**
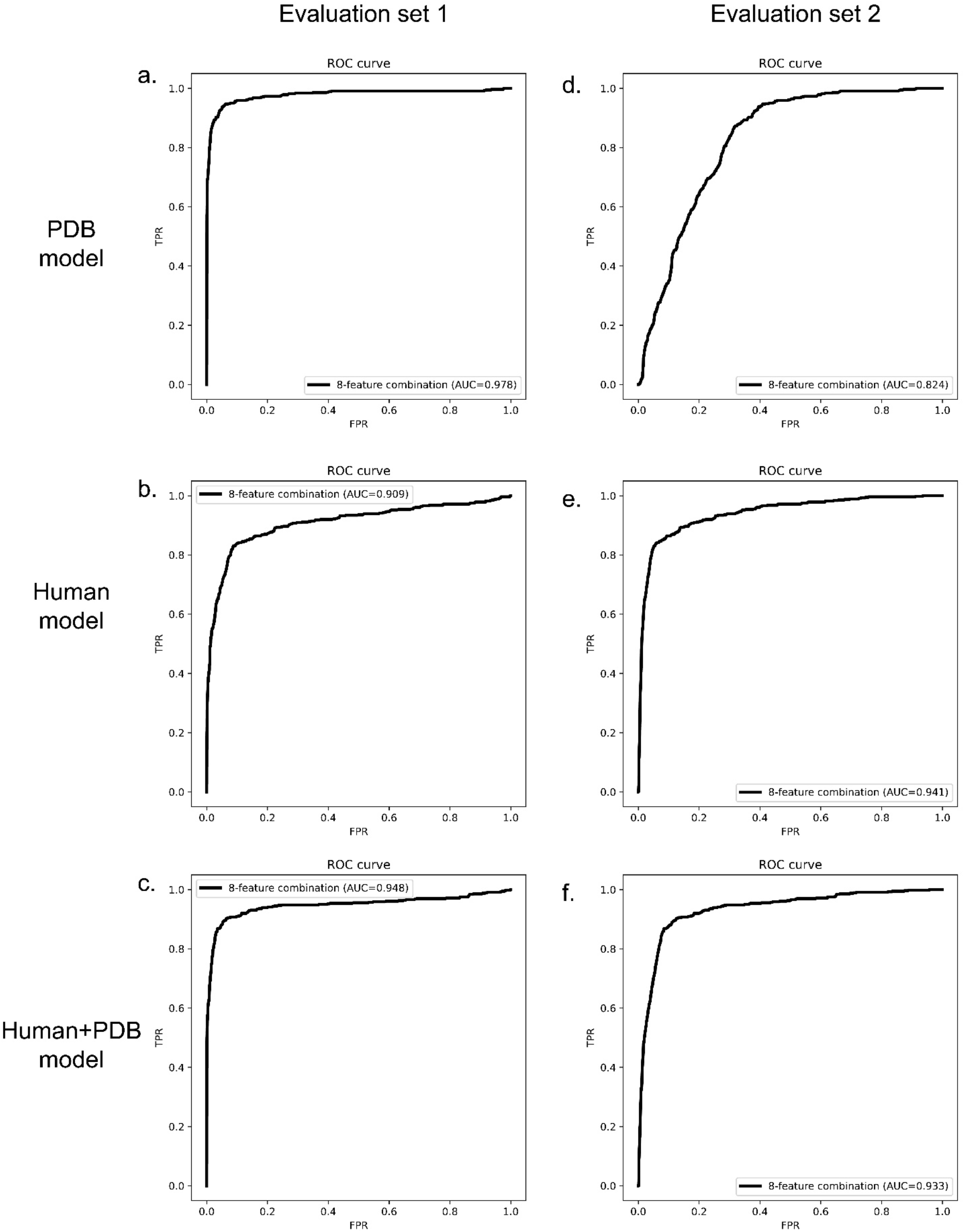
Final model performance via ROC curves, for 3 models. ROC curves on Evaluation set 1 (left) and Evaluation set 2 (right) for 3 different models: (a,d) PDB model, (b,e) Human model and (c,f) Human+PDB model.

**Figure S4.**
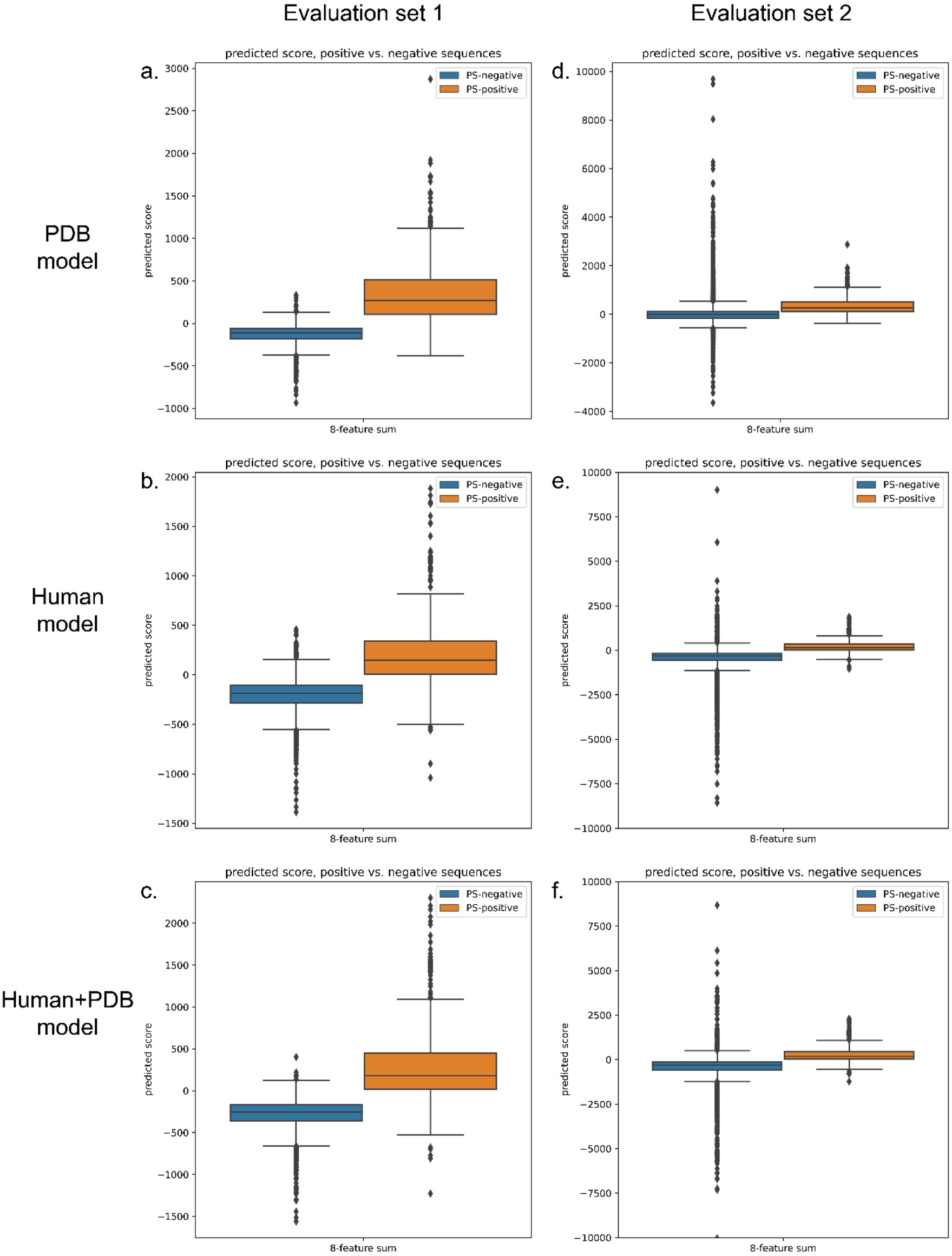
Final model performance via boxplots, for 3 models. Predicted score boxplots of positive vs. negative sequences on Evaluation set 1 (left) and Evaluation set 2 (right) for 3 different models: (a,d) PDB model, (b,e) Human model, and (c, f) Human+PDB model.

**Figure S5.**
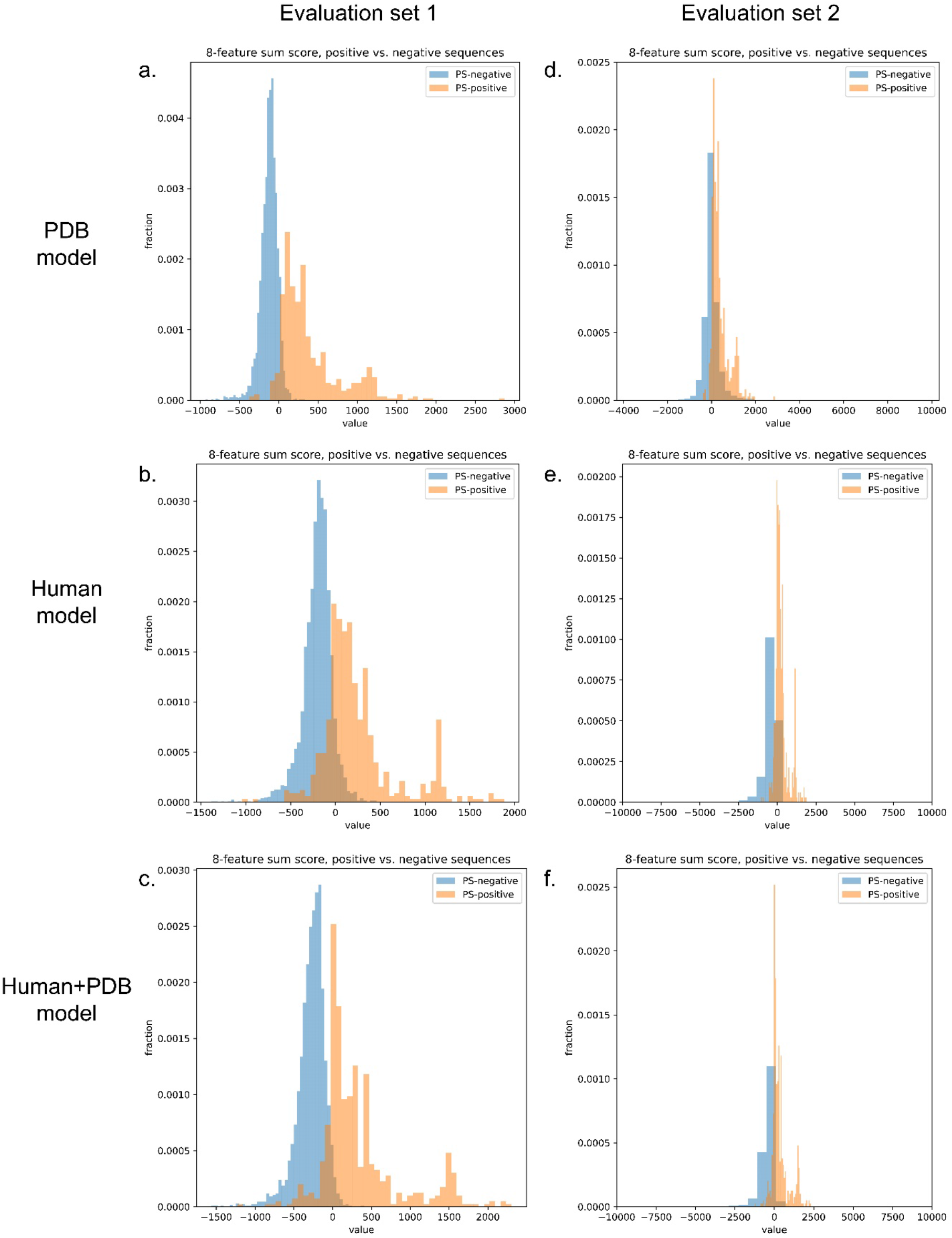
Final model performance via histograms, for 3 models. Distribution histograms of positive vs. negative sequences on Evaluation set 1 (left) and Evaluation set 2 (right) for 3 different models: (a,d) PDB model, (b,e) Human model, and (c,f) Human+PDB model.

**Figure S6.**
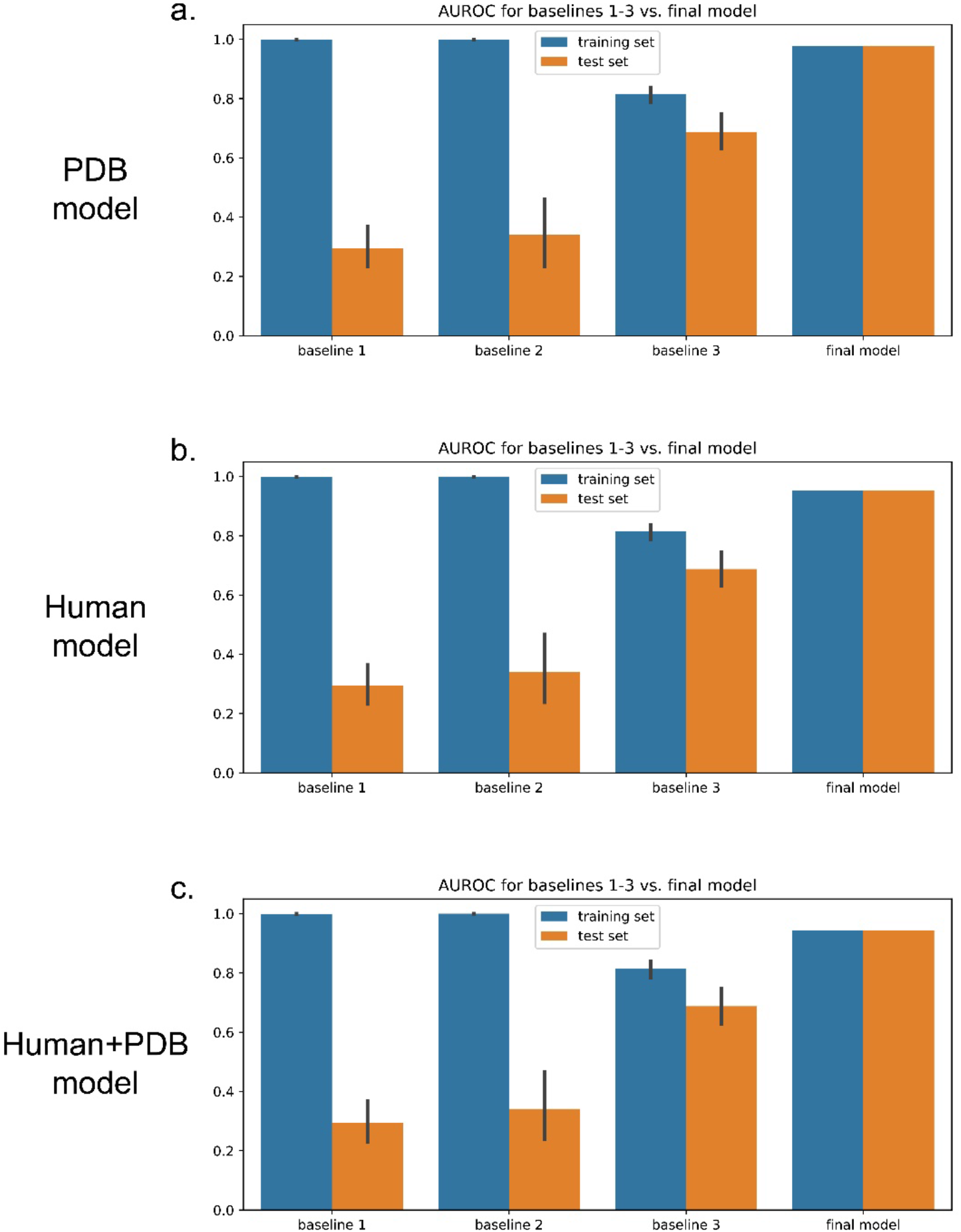
Comparison of three training baselines and the final predictor models for validation, for the 3 models. Performance comparisons are shown for (a) PDB model, (b) Human model, and (c) Human+PDB model. Baseline 1 was created by providing random values from a normal distribution N(0, 1) in the weight training step instead of providing PDB- based physical feature values into the genetic algorithm. Baseline 2 was created by providing random values from the distribution of residue-specific physical feature values instead of providing sequence-based physical feature values. Baseline 3 was created by optimizing 1 weight for 20 residue types for each physical feature (removing residue specificity) during training instead of optimizing 20 weights for 20 residue types for each physical feature.

**Figure S7.**
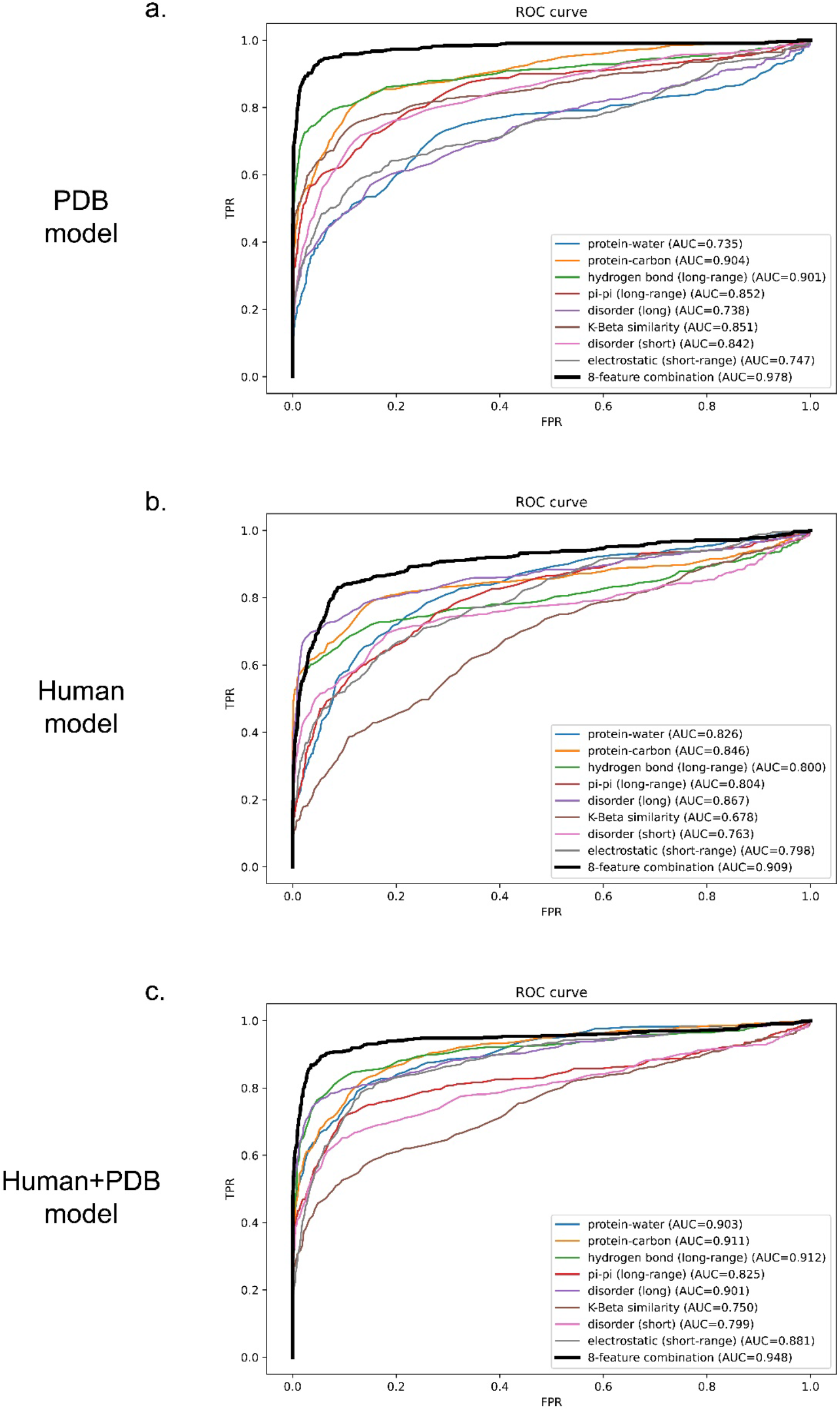
Comparison of performance via ROC curves of predictors trained on 8 features vs. 1 feature, for 3 models: (a) PDB model, (b) Human Model, and (c) Human+PDB model.

**Figure S8.**
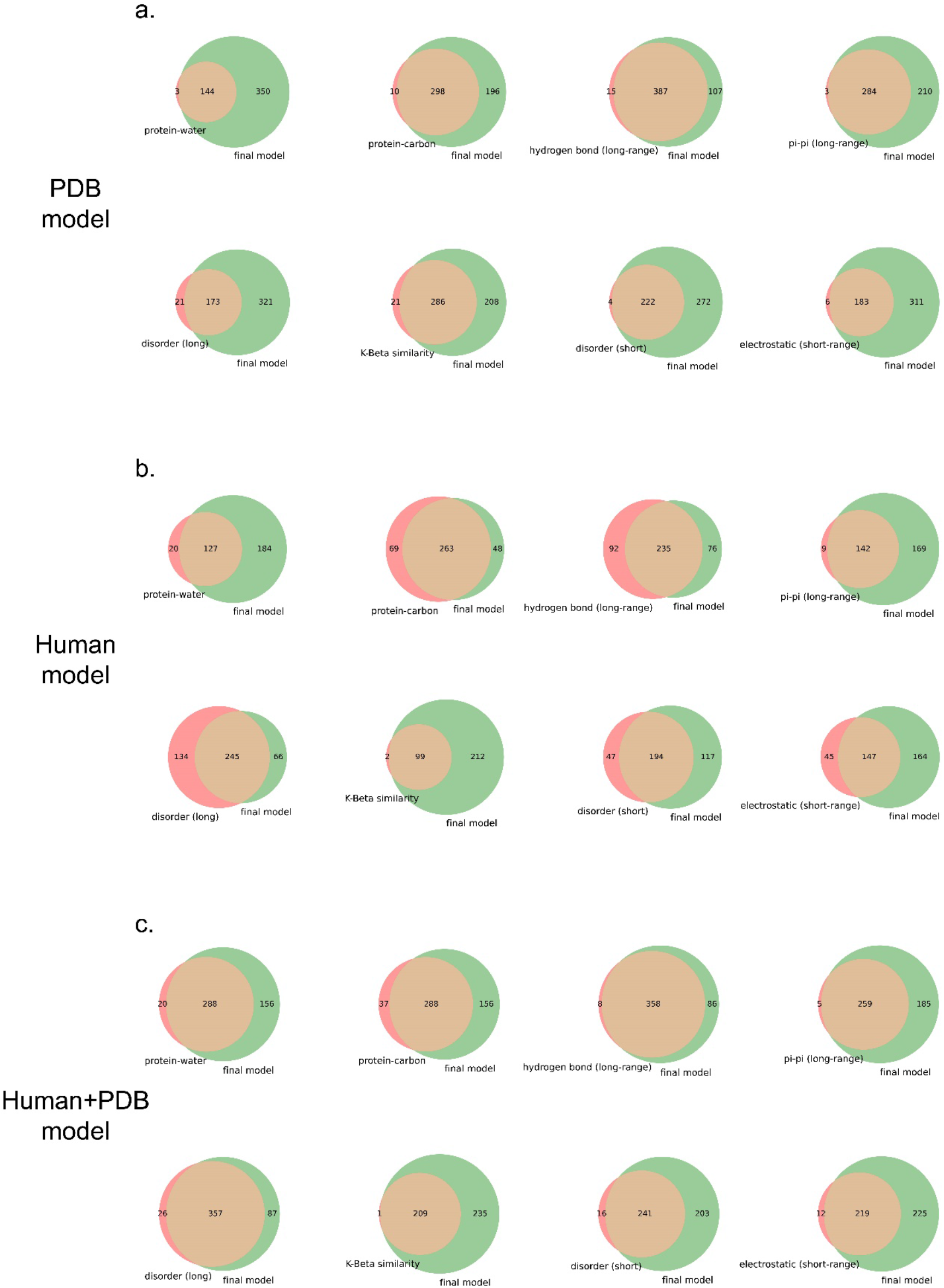
Comparison of performance via Venn diagrams of predictors trained on 8 features vs. 1 feature for the 3 models. Venn diagrams showing the coverage overlaps of PS-positive sequences by 1-feature predictors vs. the 8-feature predictor at a confidence threshold that returns 2% of the PDB, for 3 models: (a) PDB model, (b) Human model, and (c) Human+PDB model.

**Figure S9.**
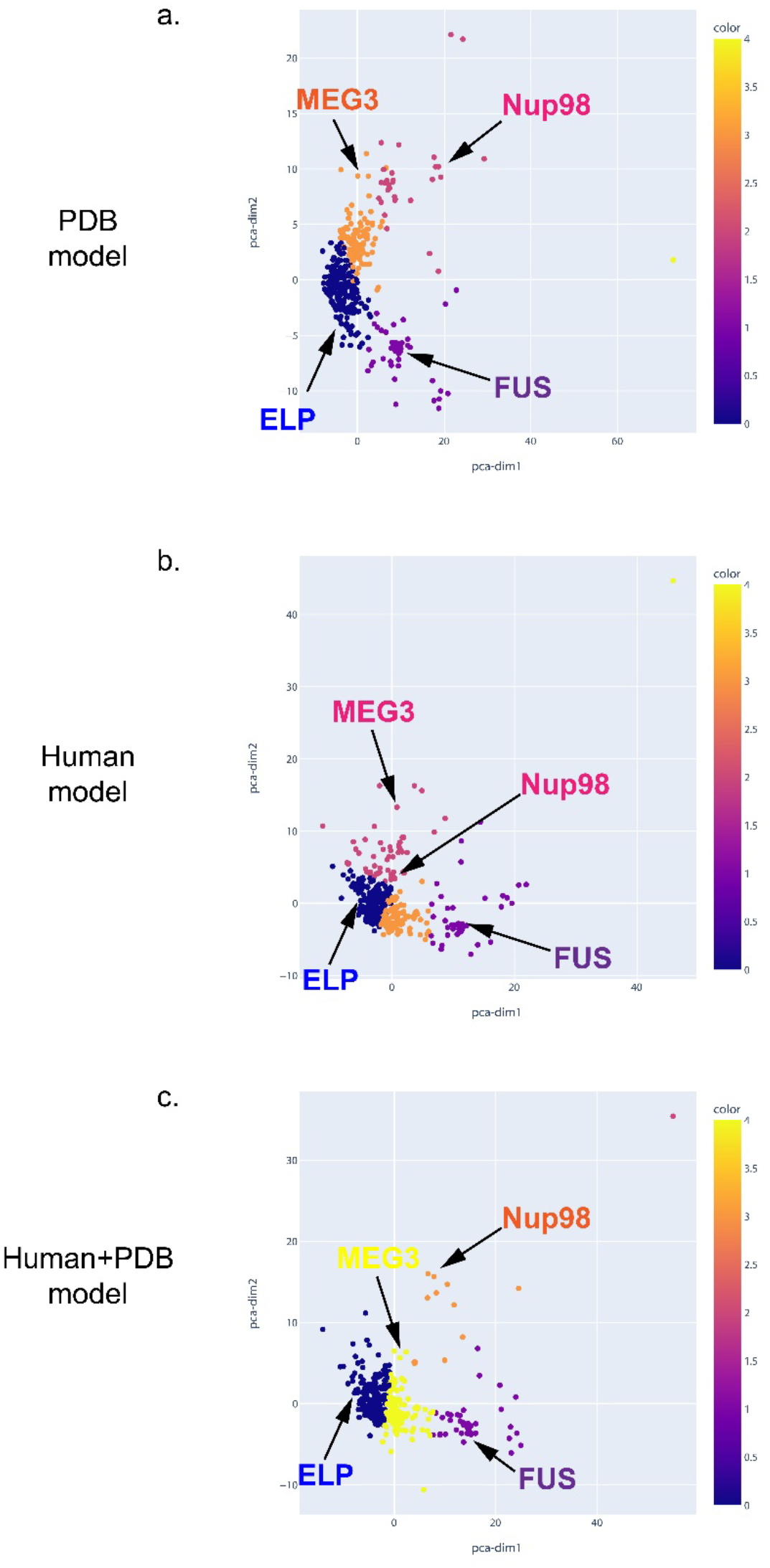
Feature score-based clustering for PS-positive proteins for the 3 models. Plots of 2 abstracted dimensions for clustering based on feature z-scores, showing the separation of different types of phase-separating sequences, for 3 models: (a) PDB model, (b) Human model, and (c) Human+PDB model.

**Figure S10.**
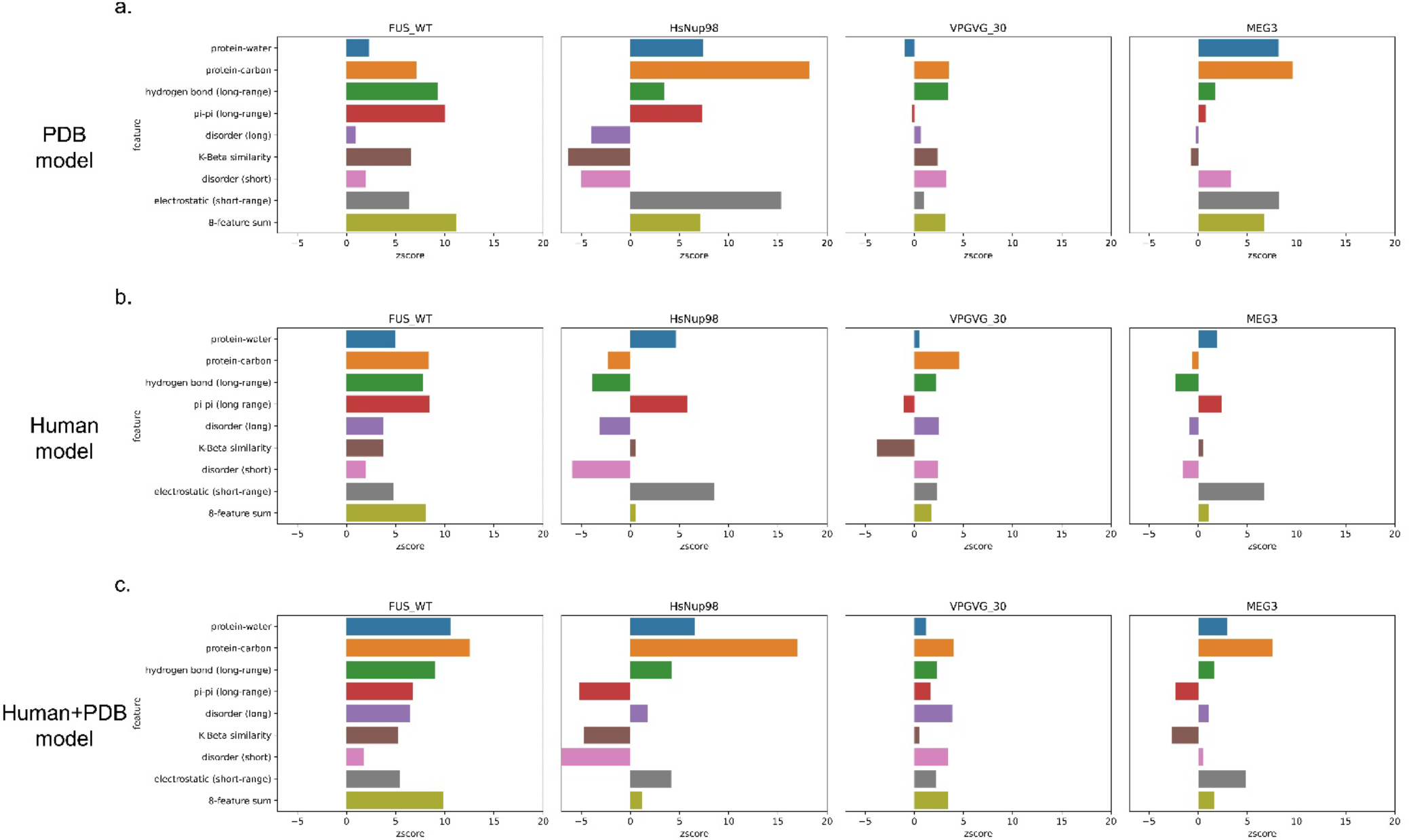
Feature score breakdown for example sequences from distinct clusters of PS-positive proteins for the 3 models. The score breakdown of 4 example sequences from 4 clusters in Fig S8 is shown for FUS (human), Nup98 (human), elastin-like peptide (ELP, VPGVG_30, 30 repeats of VPGVG) and MEG-3 (*C. elegans*) for 3 models: (a) PDB model, (b) Human model, and (c) Human+PDB model.

**Figure S11.**
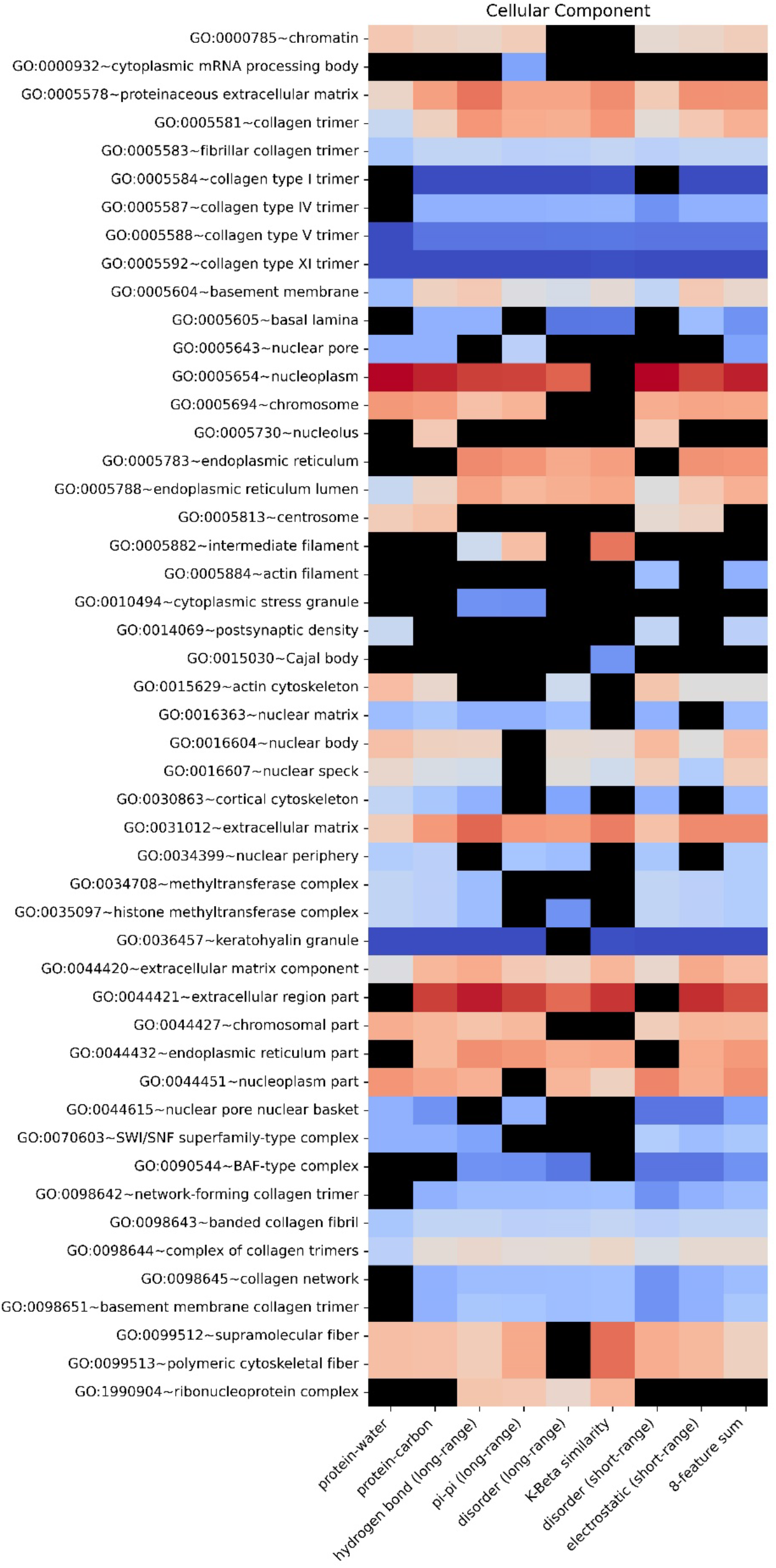
Enrichment heatmap by GO functional annotations for different features for the PDB model. Heatmap showing the enrichment of proteins with a given functional annotation that fall under a 2% confidence threshold for each single feature score and 8-feature sum score. The color gradient shows the natural logarithm of the enrichment percentage.

**Table S1.**
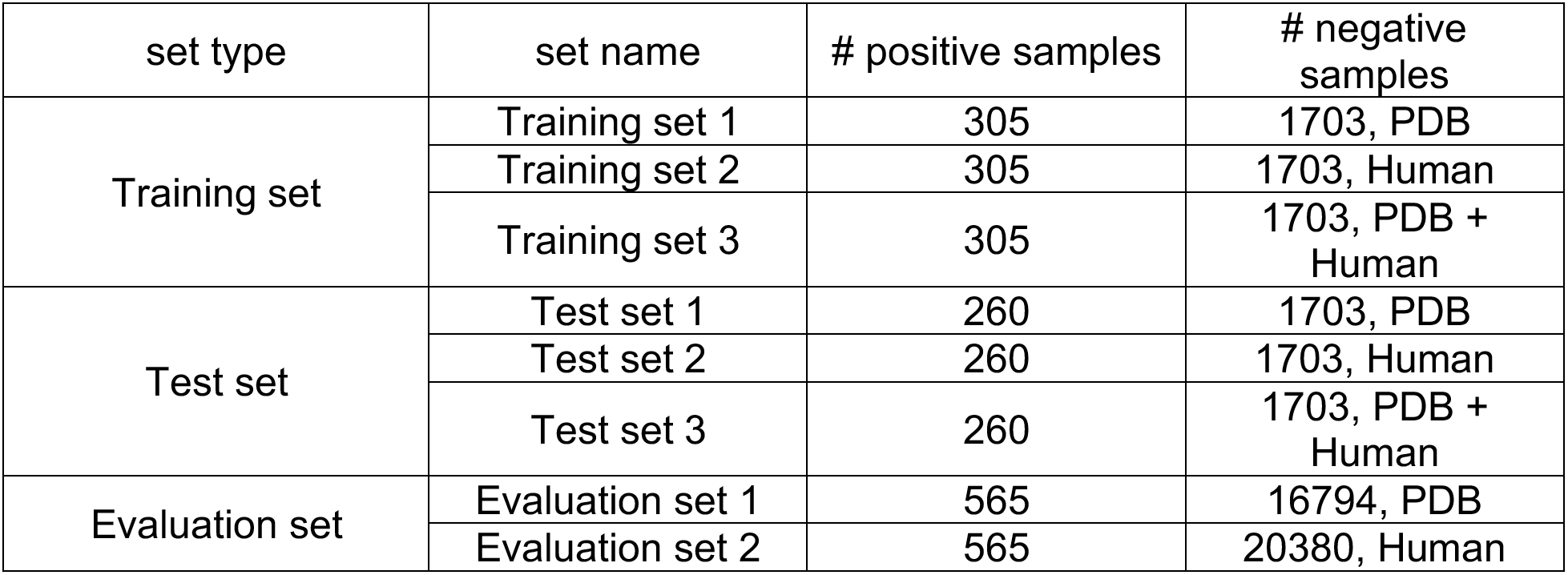
Constructed Training/Test/Evaluation sets and number of sequences in each set.

**Table S2.**
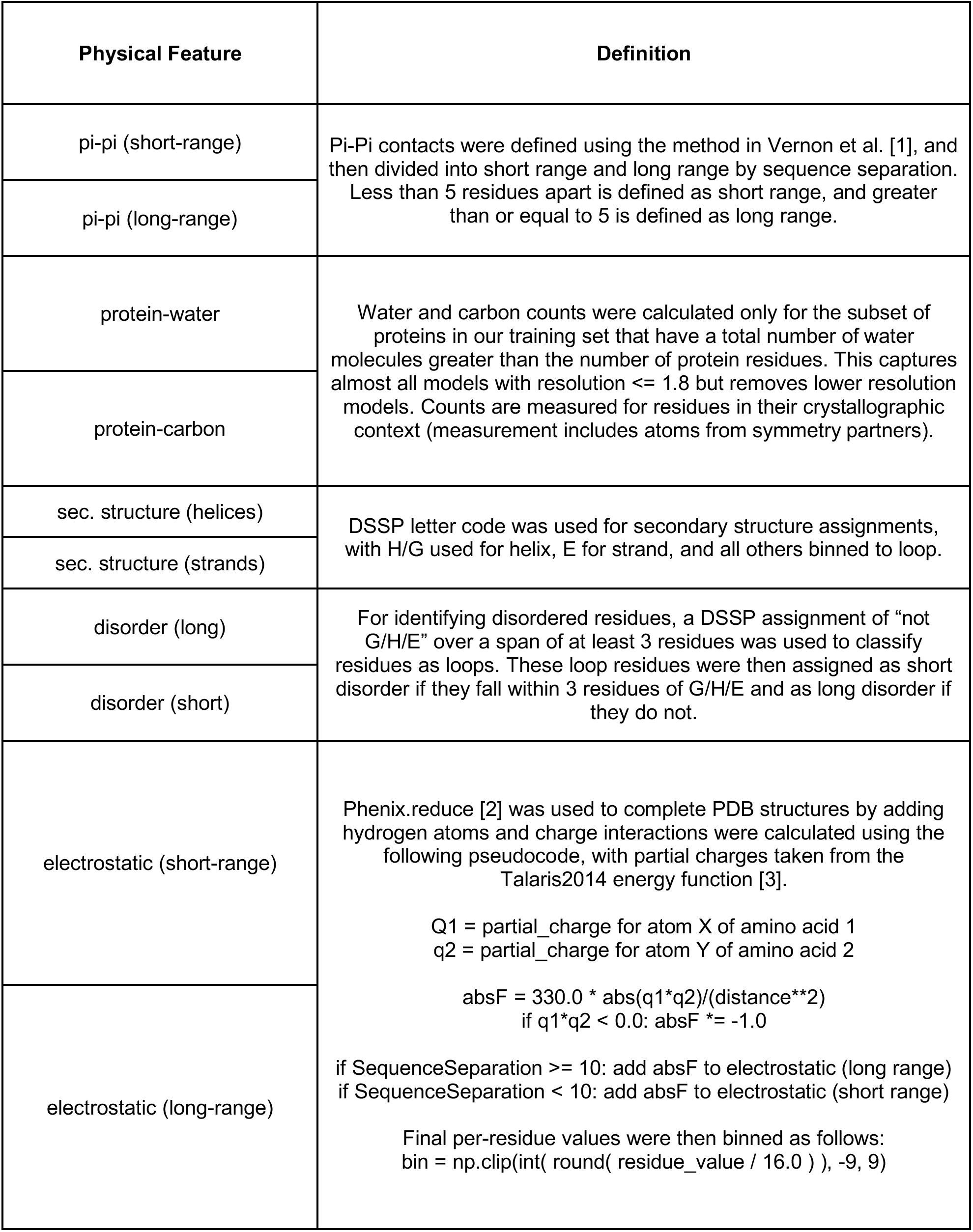
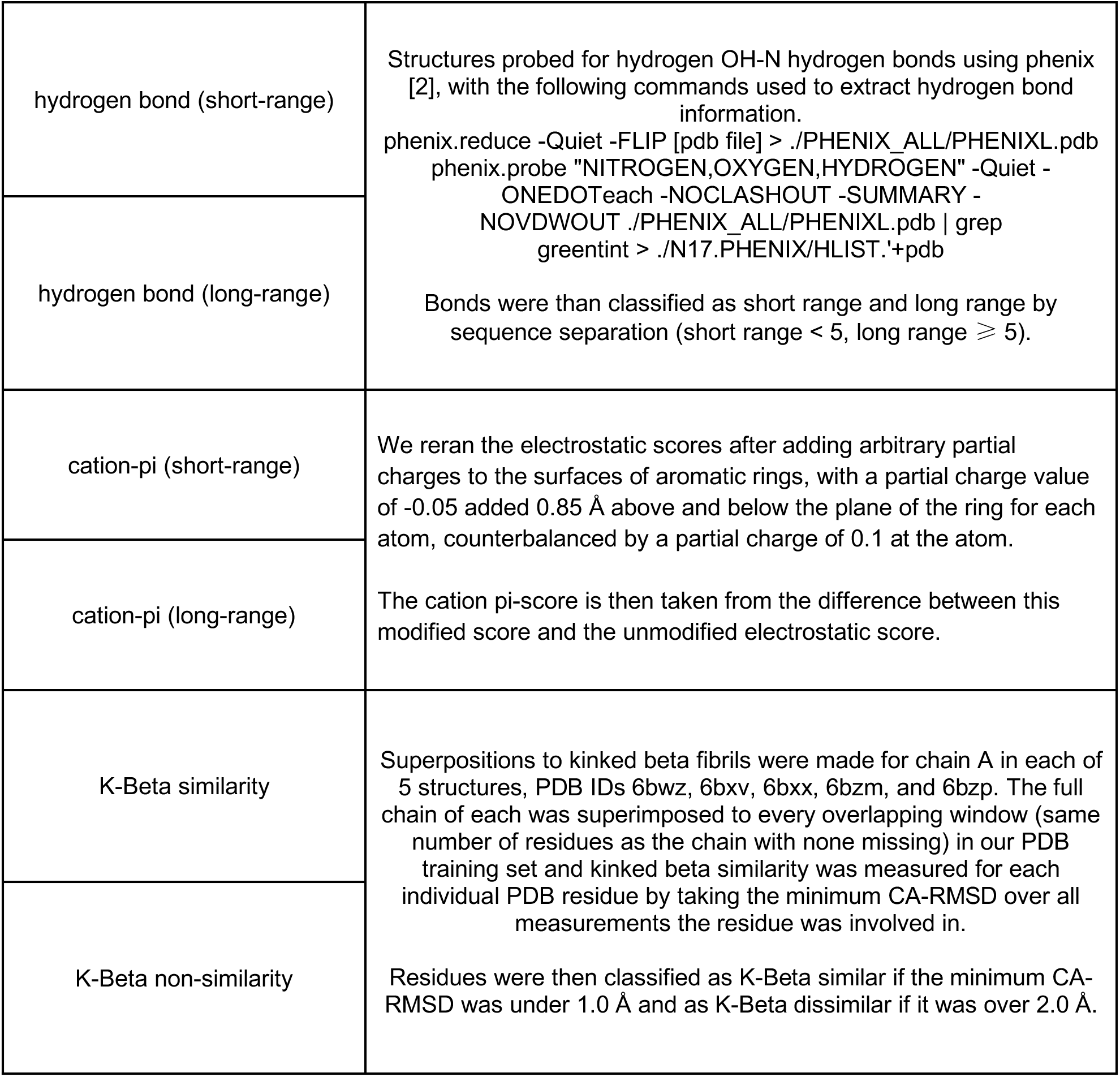
16 physical features with corresponding definitions.

**Table S3.**
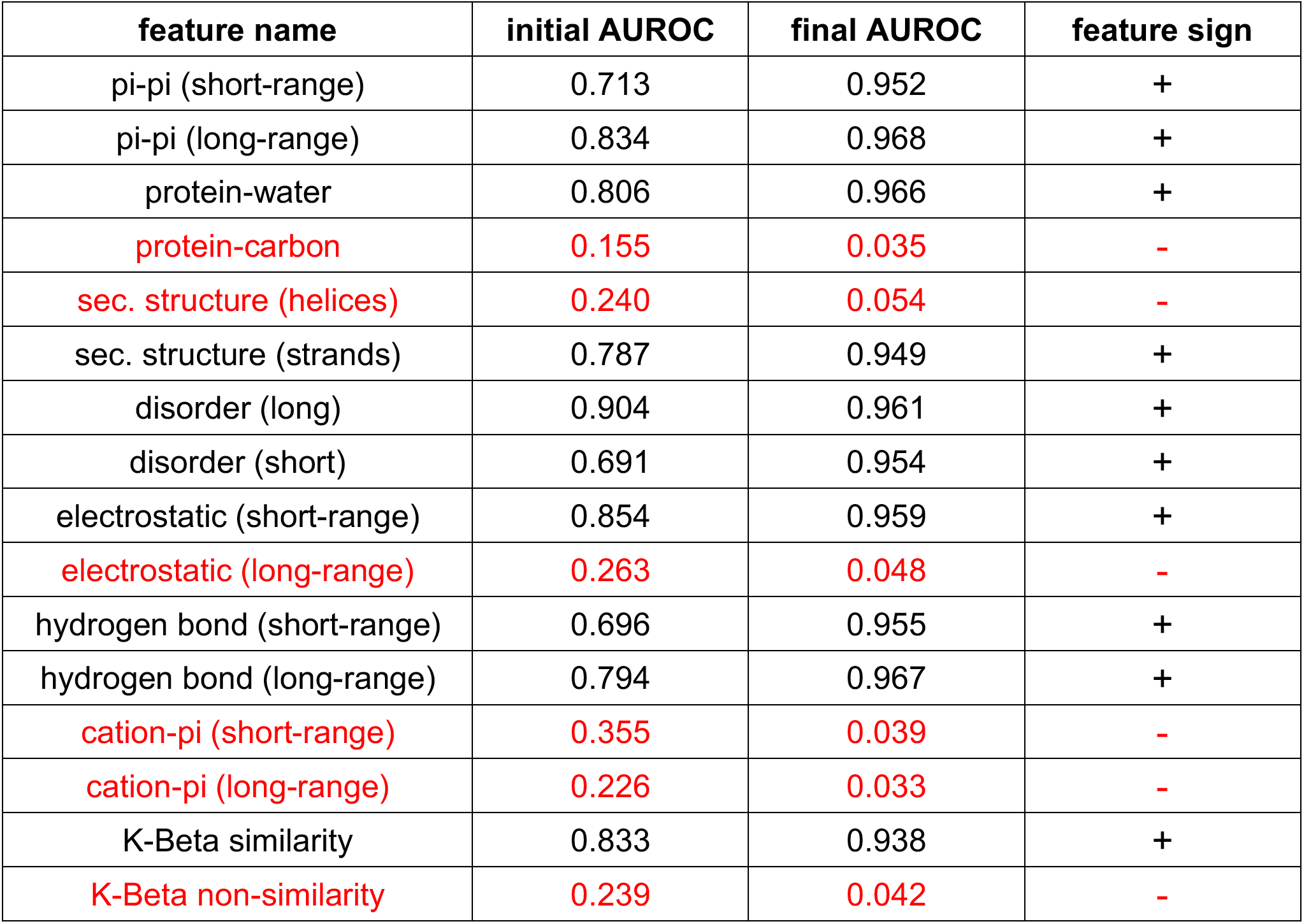
The “signs” of 16 features determined by AUROC direction during individual feature training. Feature with a positive correlation to phase separation prediction are in black and those with a negative correlation to phase separation prediction are in red.

**Table S4.**
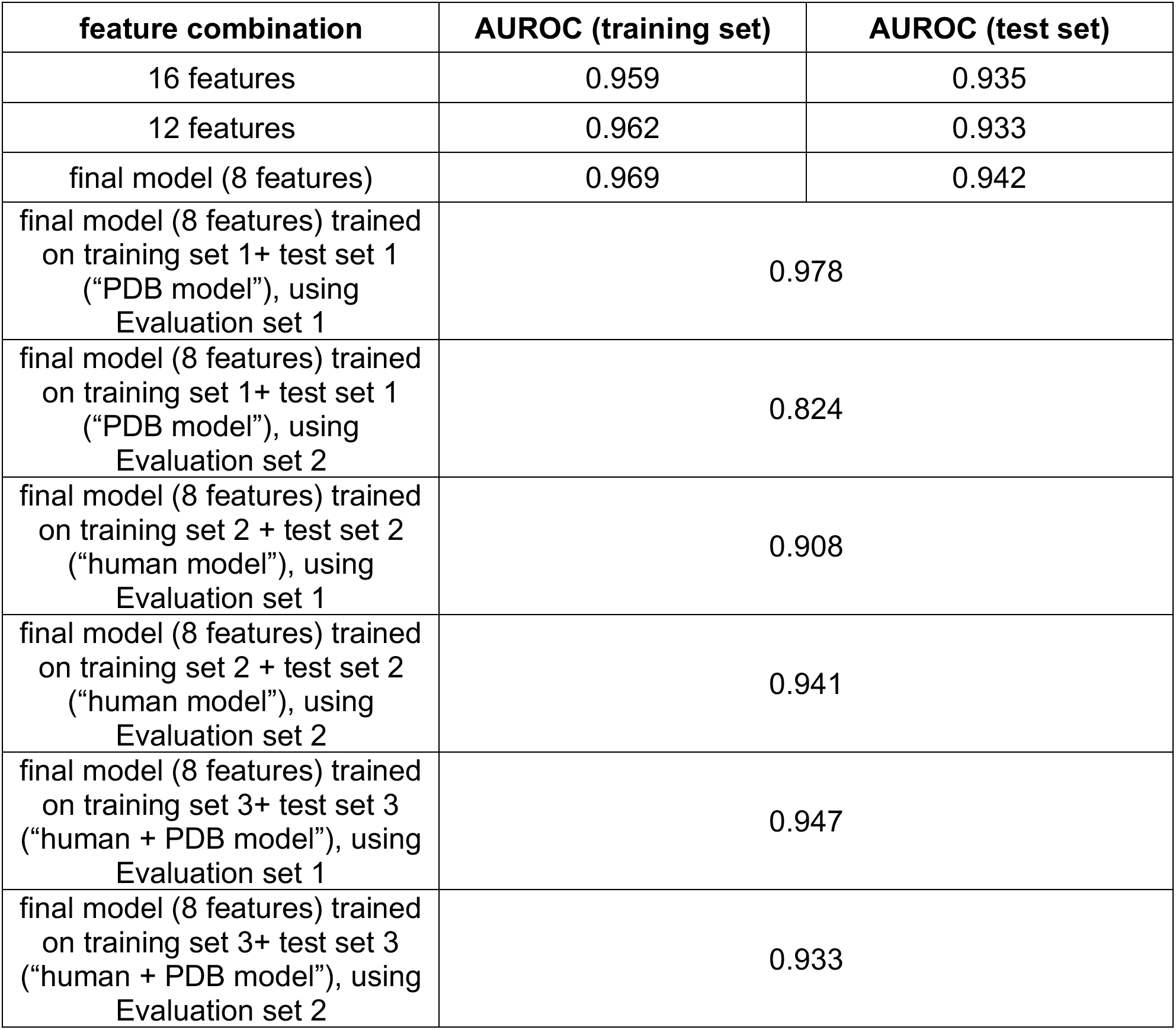
The AUROC of different trained feature combinations.

## Separate Supplementary Files

One attached Excel file contains, on separate tabs, **Tables S5-S10**.

**Table S5.** GO enrichment analysis.

**Table S6.** Uniprot IDs of 6102 sequences from human proteome using CRAPome as filtering method.

**Table S7.** Uniprot IDs of 3406 sequences from PDB base.

**Table S8.** Uniprot IDs of 3406 sequences randomly selected from human base in Table S6.

**Table S9.** Uniprot IDs of 6812 sequences from PDB+human base.

**Table S10.** Detailed information of 565 LLPS-positive sequences with PMID of each sequence’s paper.

**File S1**. 565 PS-positive sequences (fasta file).

**File S2**. 16794 PDB sequences (fasta file).

**File S3**. 20380 Human sequences (fasta file).

