## Supplementary figures and images for "An interpretable machine learning algorithm to predict disordered protein phase separation based on biophysical interactions"

### Fig_S1_dendrogram_vertical.jpg

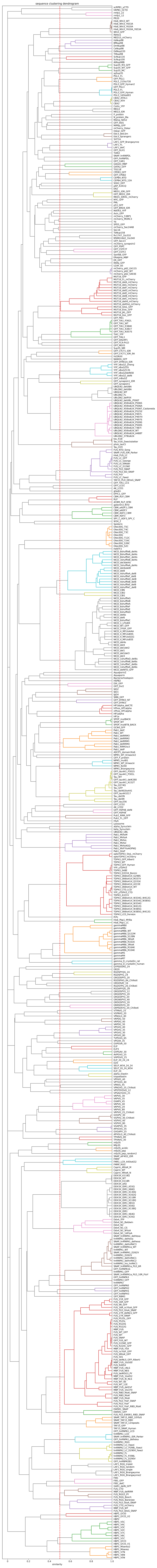

### Fig_S2_Clustermap.jpg

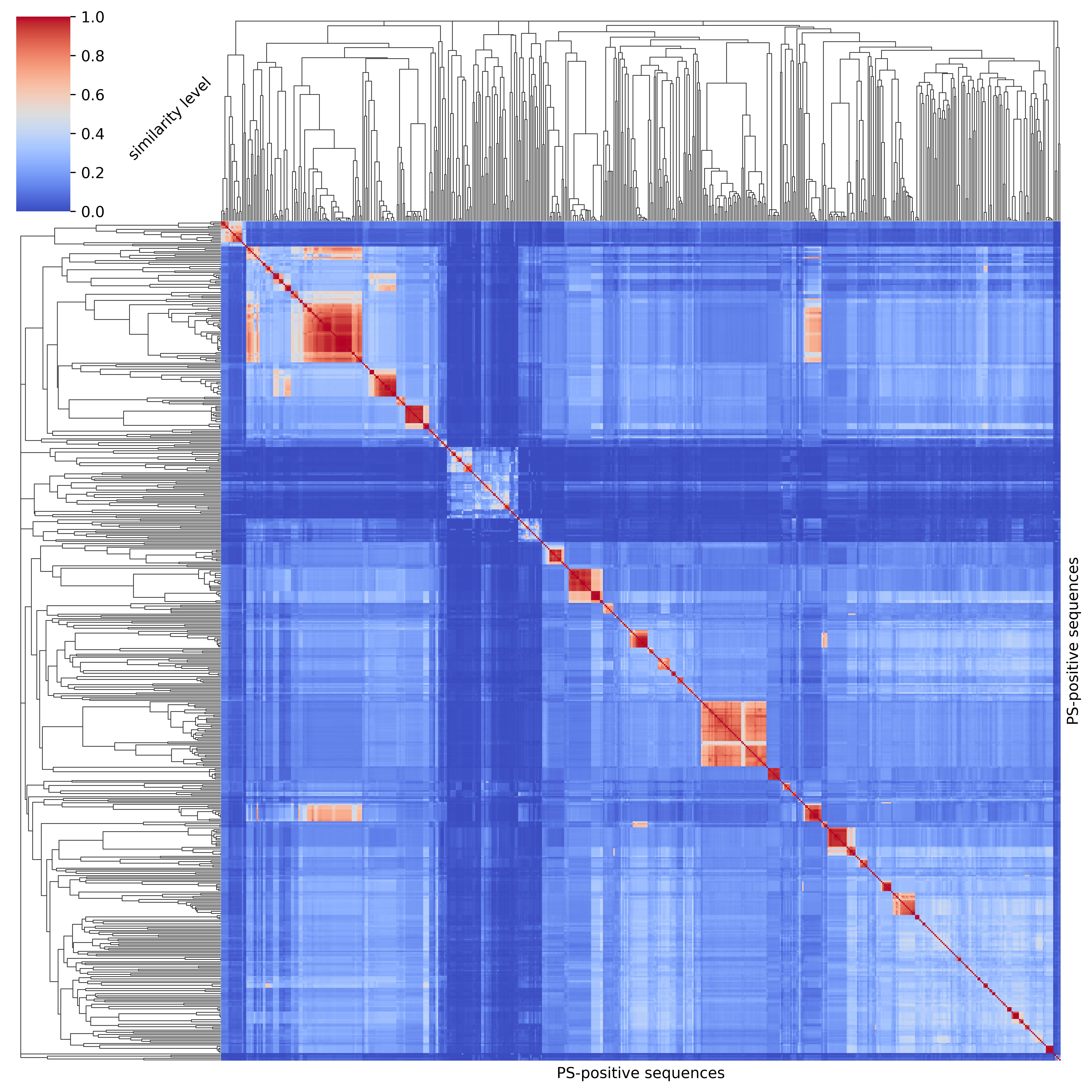

### Fig_S3_Final_model_performance_3models-AUC.jpg

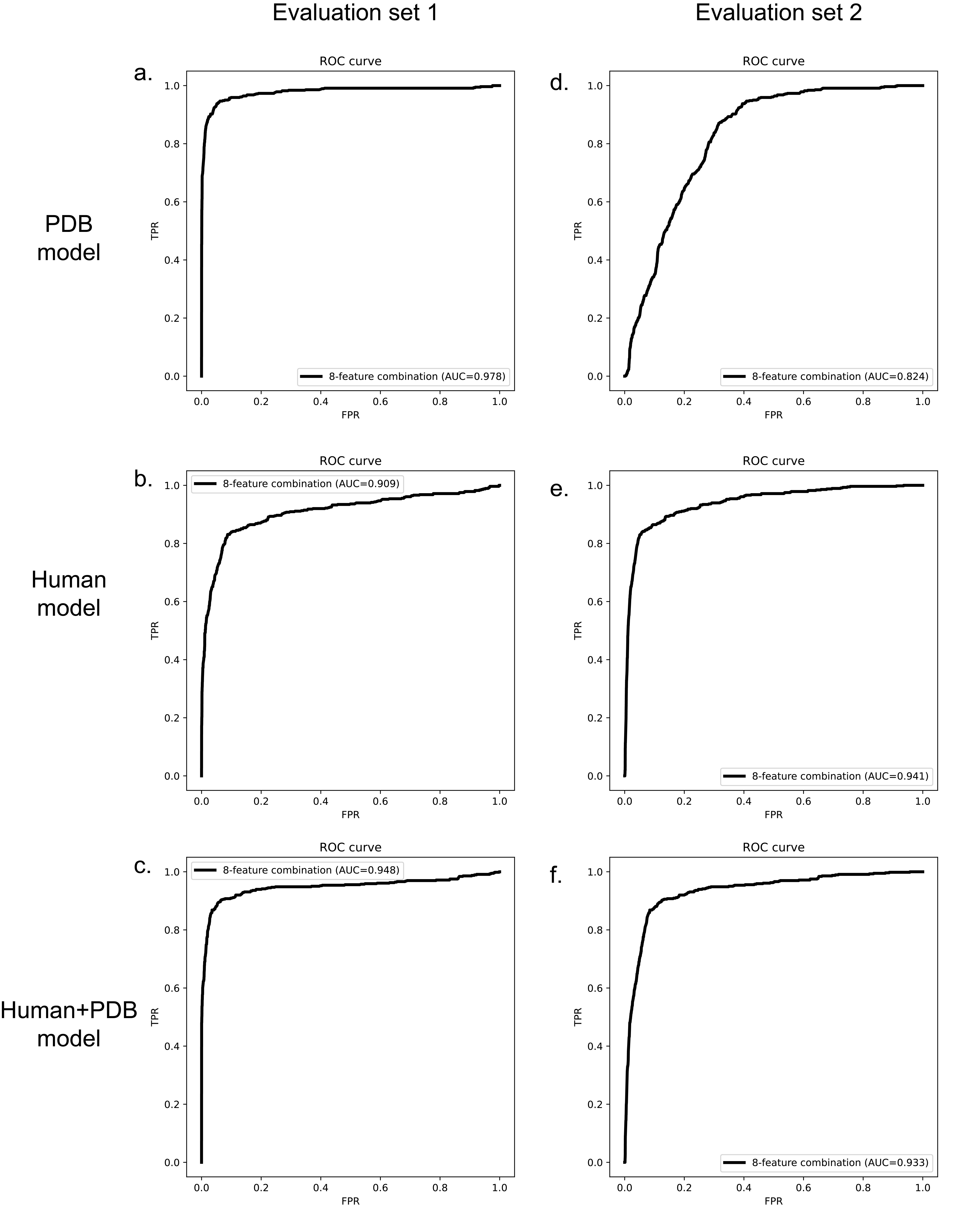

### Fig_S4_Final_model_performance_3models-boxplot.jpg

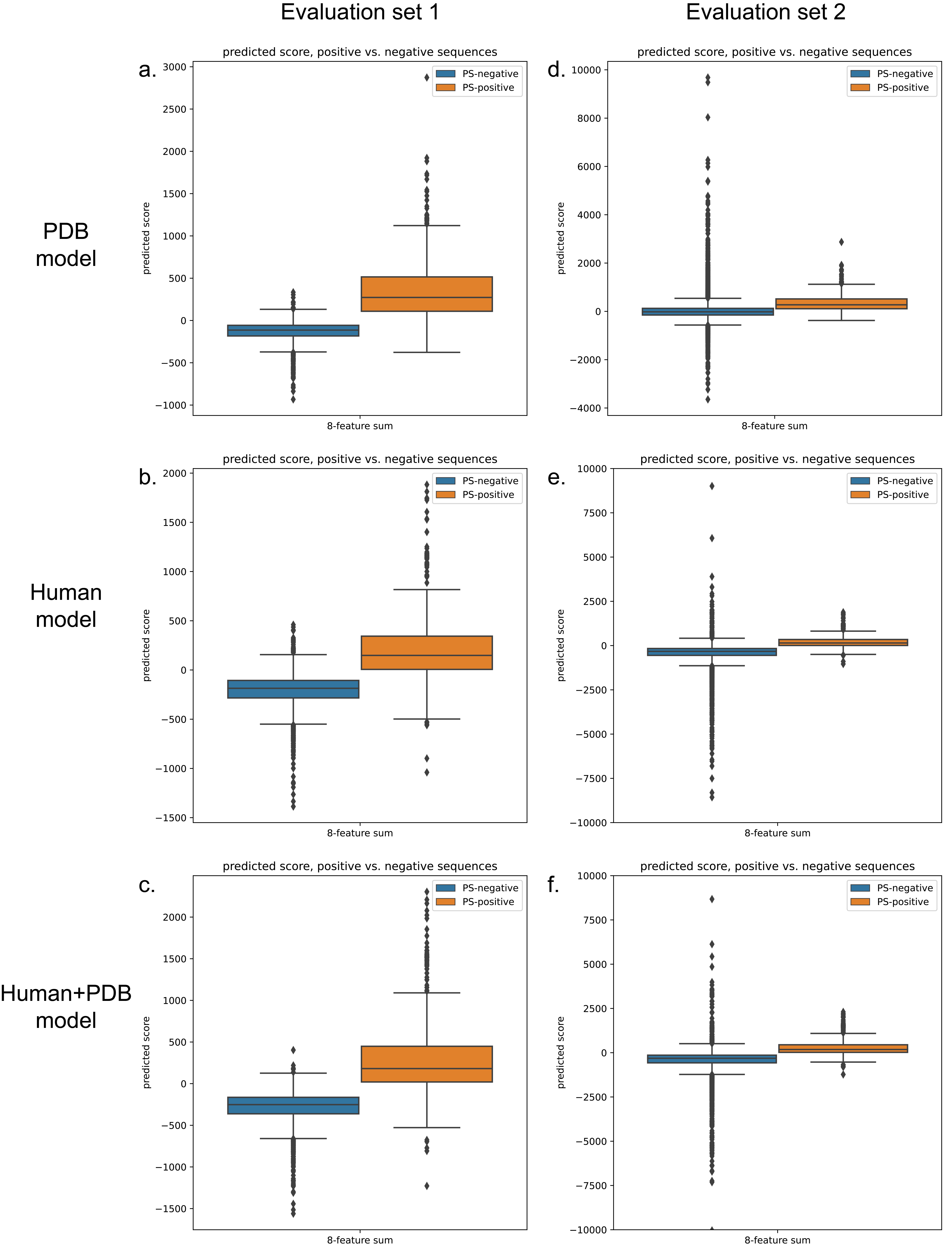

### Fig_S5_Final_model_performance_3models-histogram.jpg

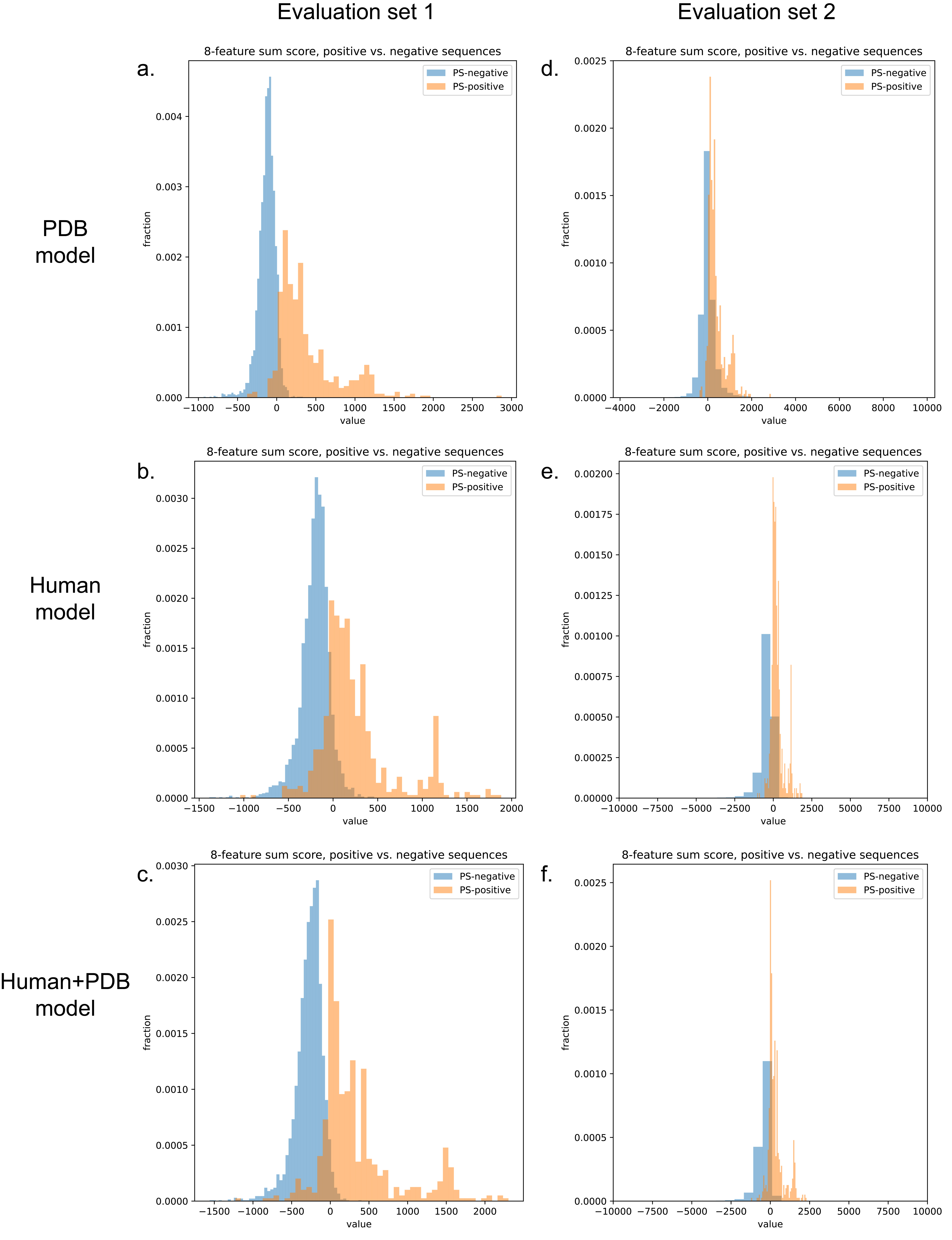

### Fig_S6_Training_baseline_vs_final_model_3models.jpg

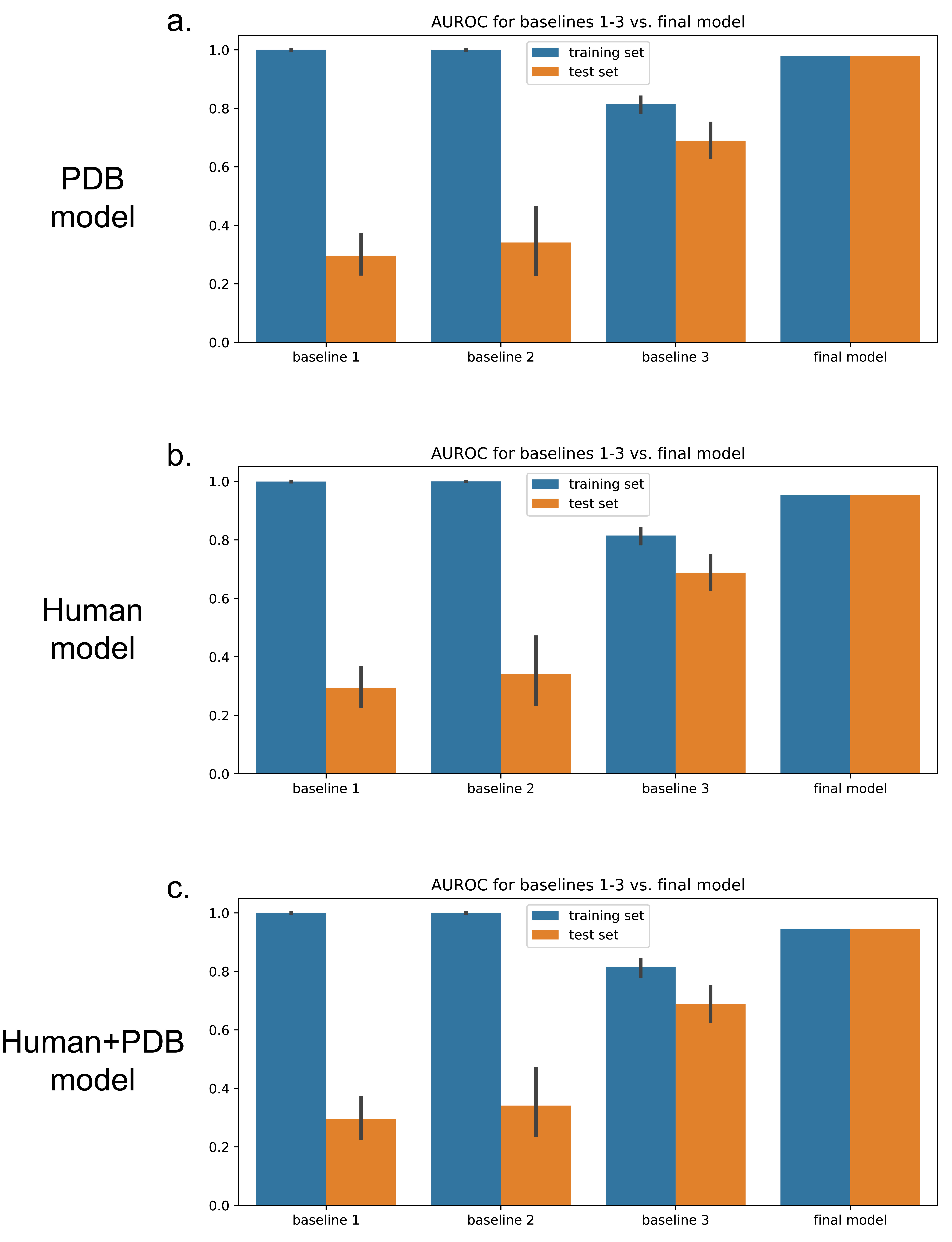

### Fig_S7_8features_vs_1feature_3models-AUC.jpg

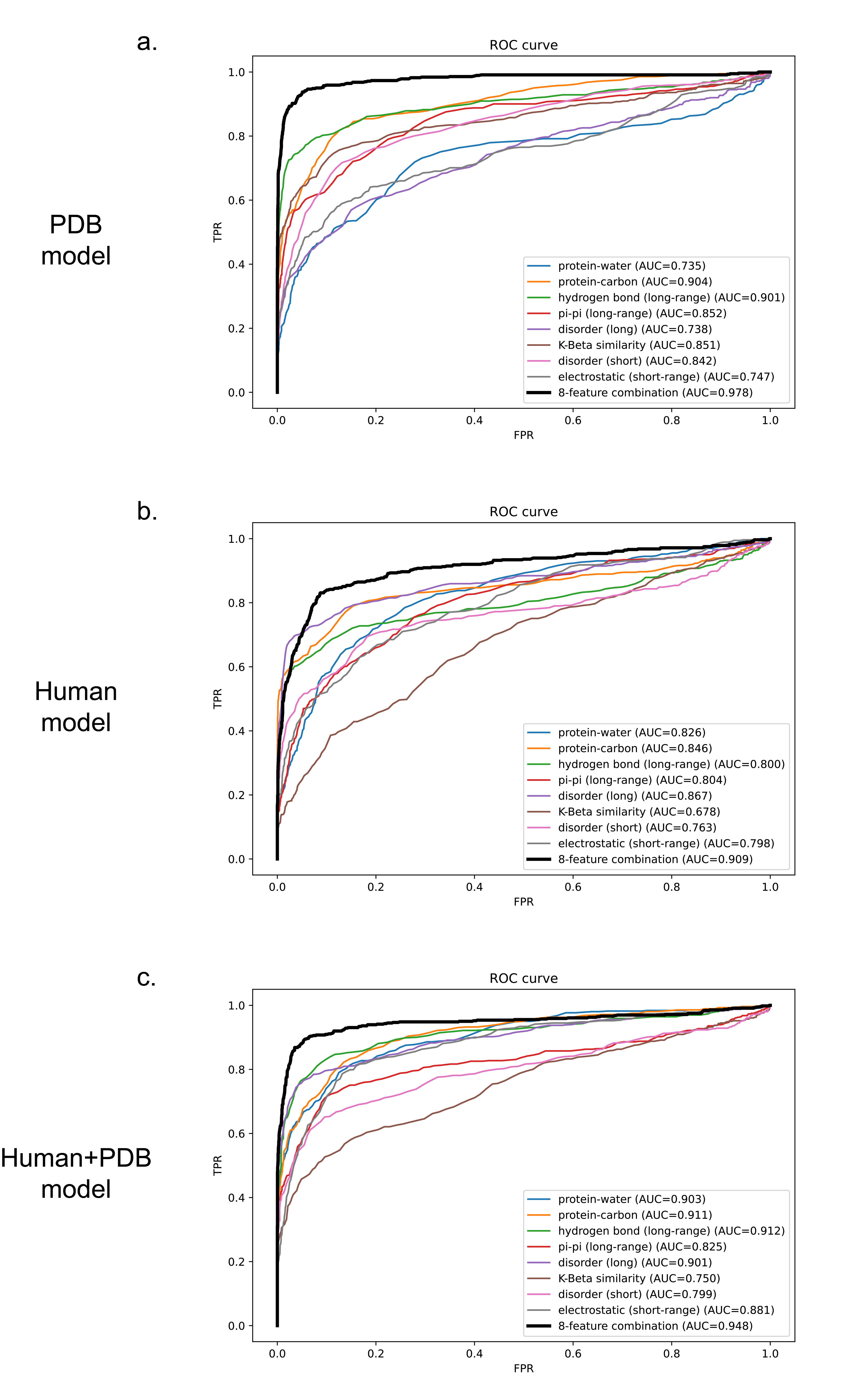

### Fig_S8_8features_vs_1feature_3models-venn.jpg

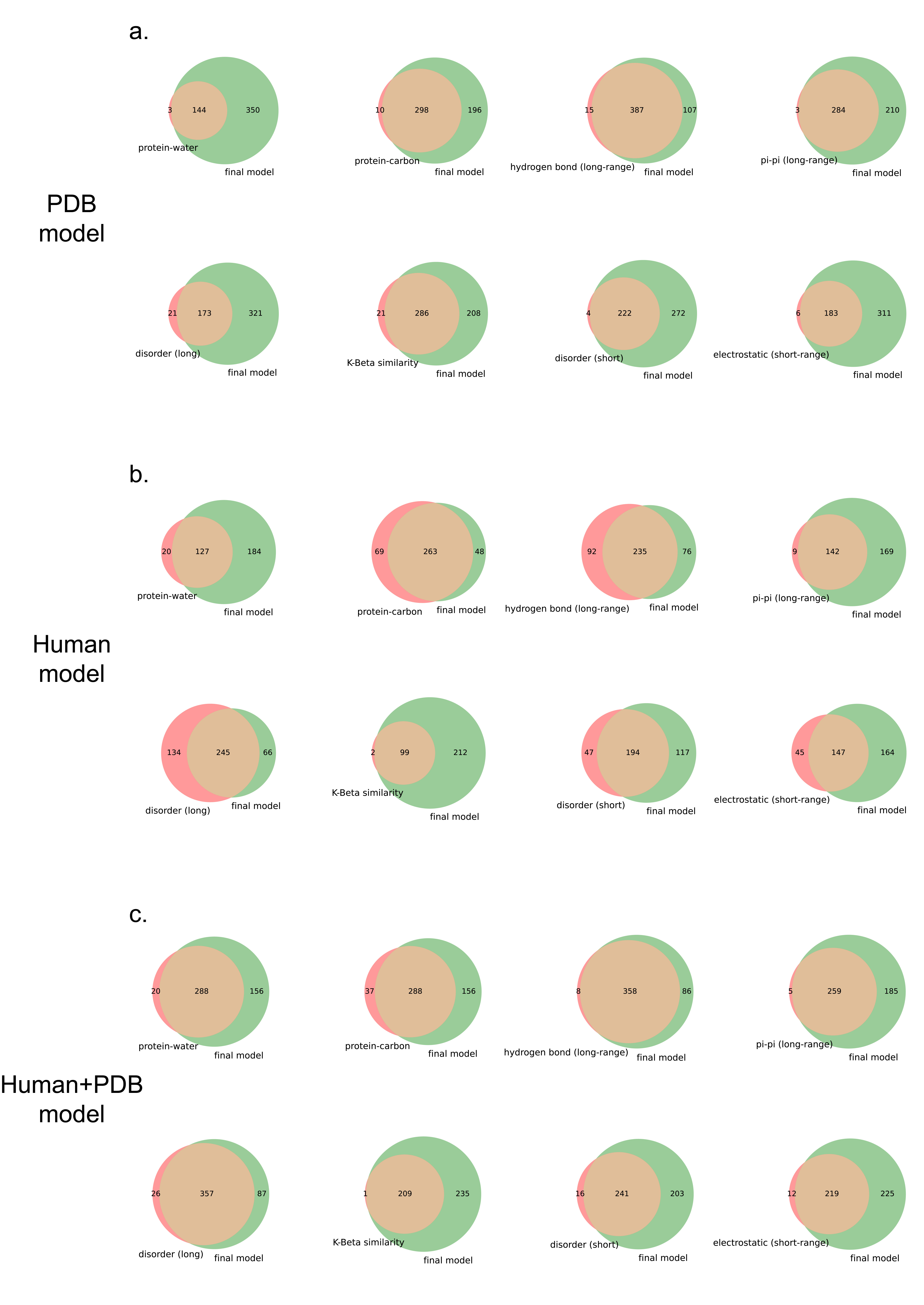

### Fig_S9_Feature_fraction_based_clustering_3models-clusters.jpg

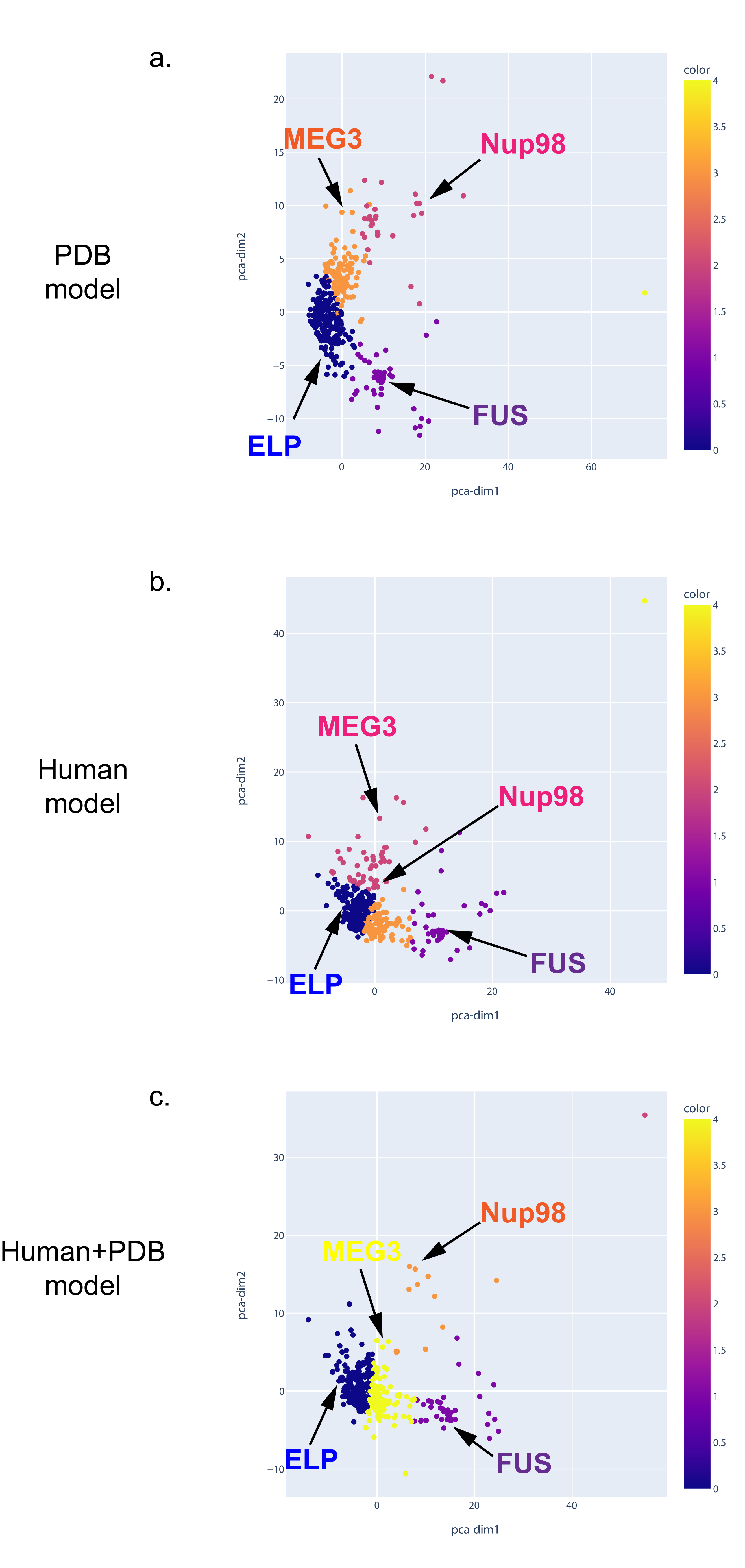

### Fig_S10_Feature_fraction_based_clustering_3models-examples.jpg

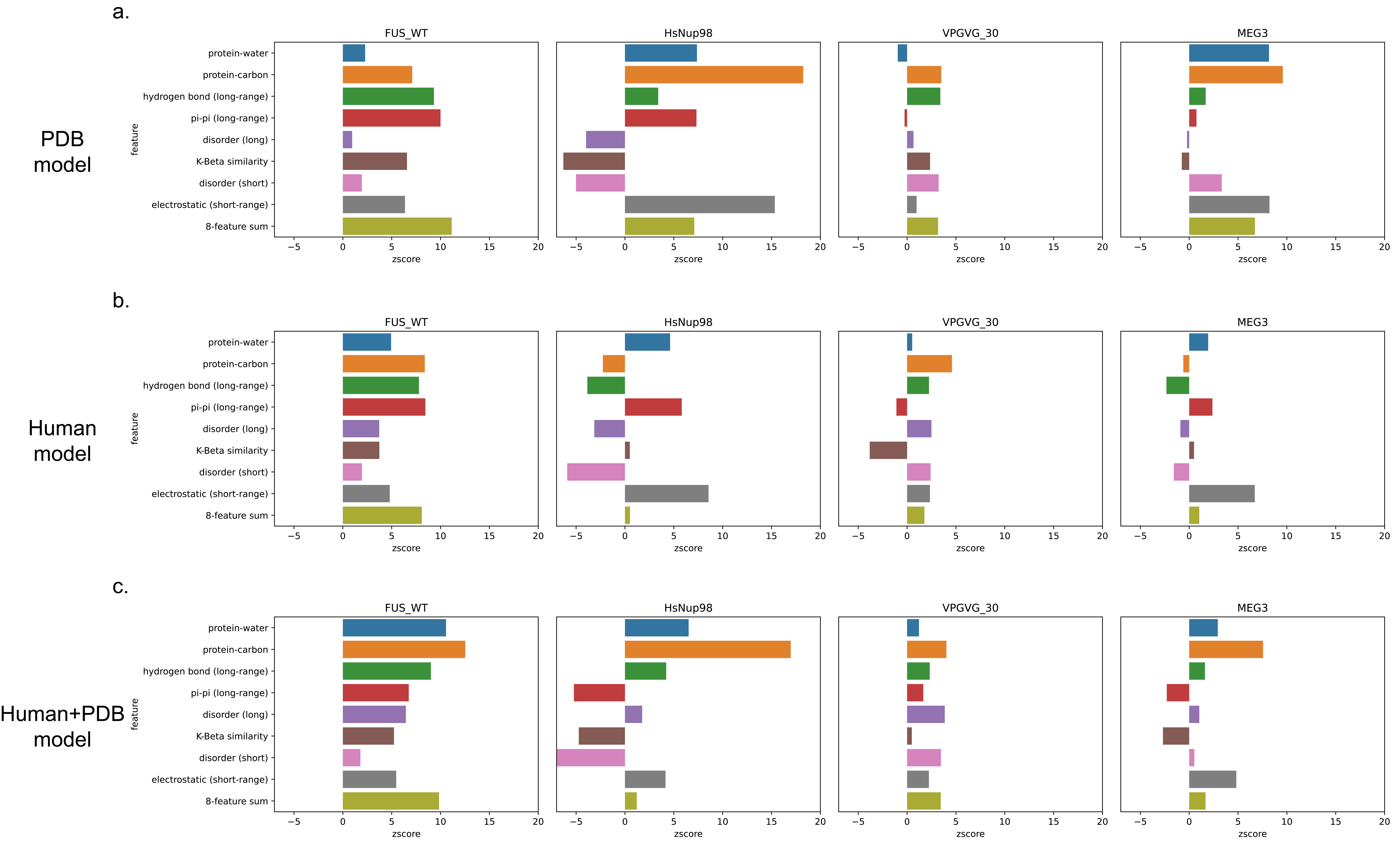

### Fig_S11_GO_Analysis-PDB.jpg

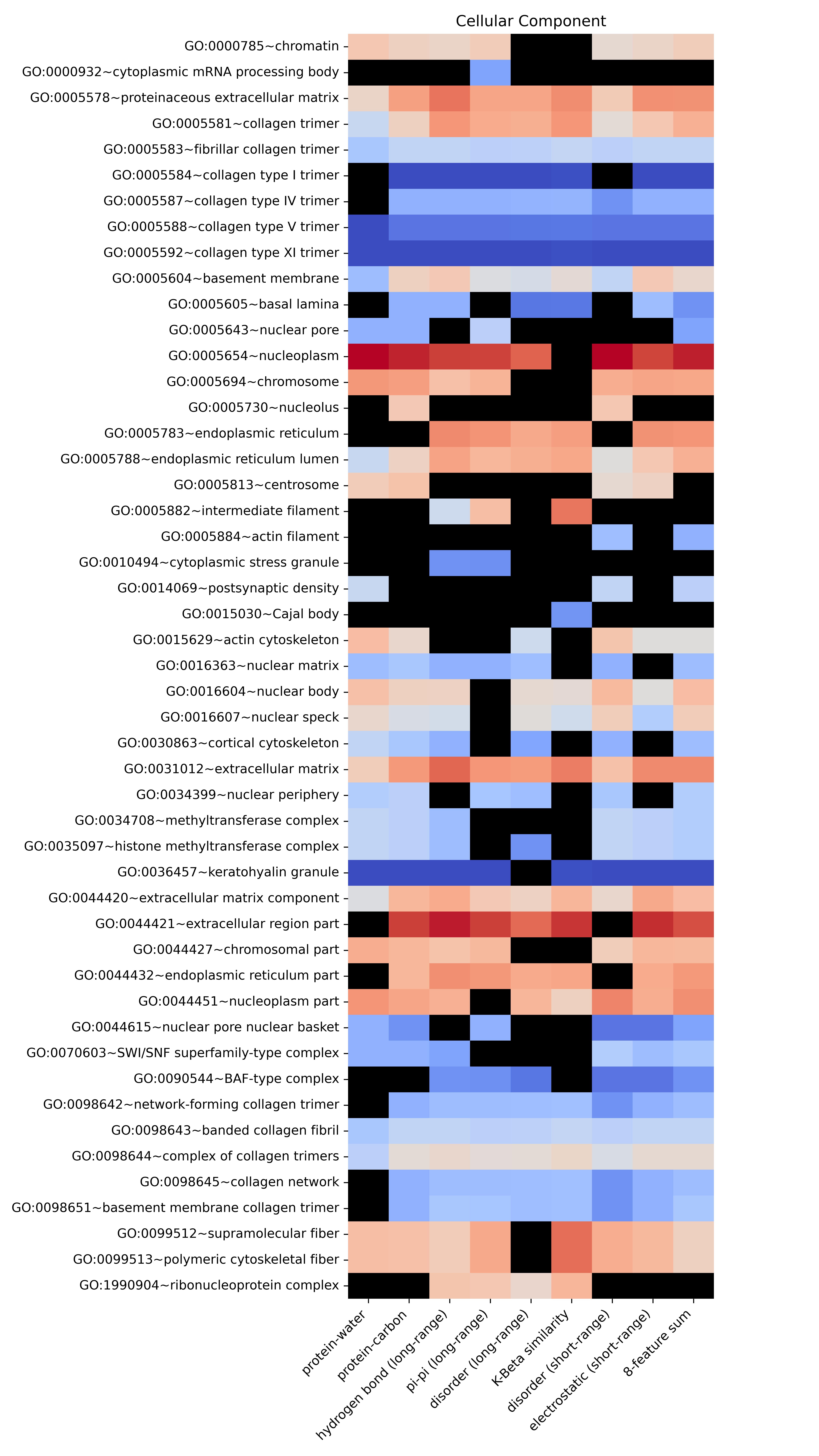
