## Supplementary_Material for "An interpretable machine learning algorithm to predict disordered protein phase separation based on biophysical interactions"

[1]Molecular Medicine Program, Hospital for Sick Children, Toronto, Ontario M5G 0A4, Canada
[2]Department of Biochemistry, University of Toronto, Toronto, Ontario, M5S 1A8, Canada
[+]Current Address: Shopify Inc., 150 Elgin Street 8th Floor, Ottawa, ON K2P 1L4, Canada
[++]Current Address: Amgen Inc. Research and Development, Burnaby, BC V5A 1V7, Canada

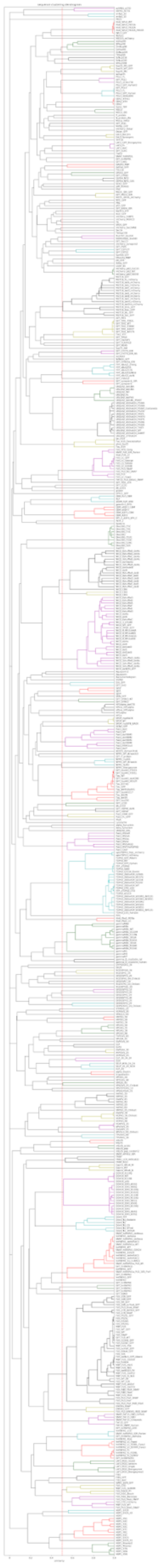

**Figure S1**. Hierarchical clustering dendrogram of PS-positive sequences.

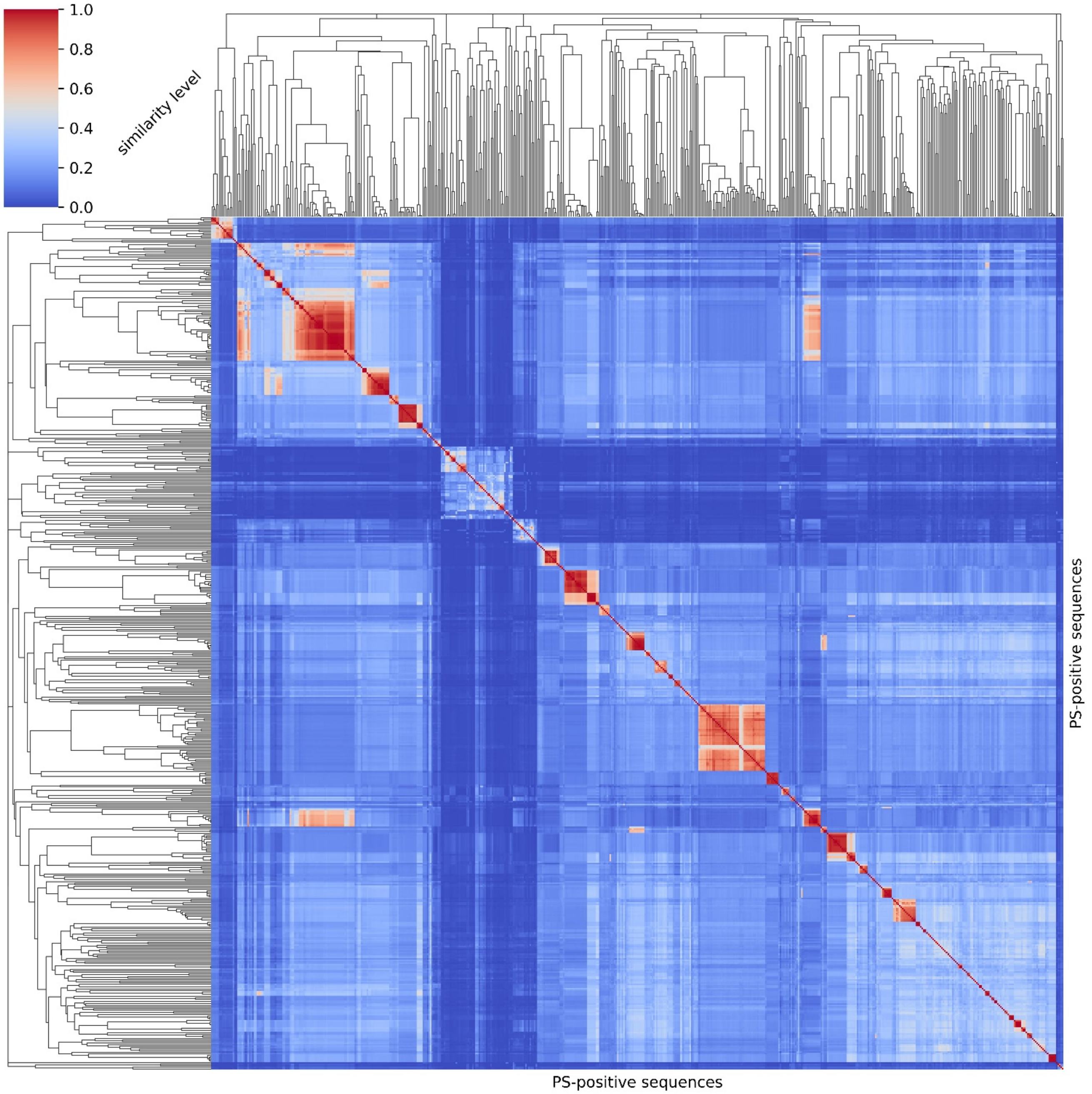


**Figure S2**. Sequence clustering map of PS-positive sequences. Left and top: clustering dendrograms showing the pairwise similarity level (0 - 1.0) for all sequences. Middle: heatmap of similarity levels showing that inter-cluster sequences have pairwise similarity higher than 0.5 in general.

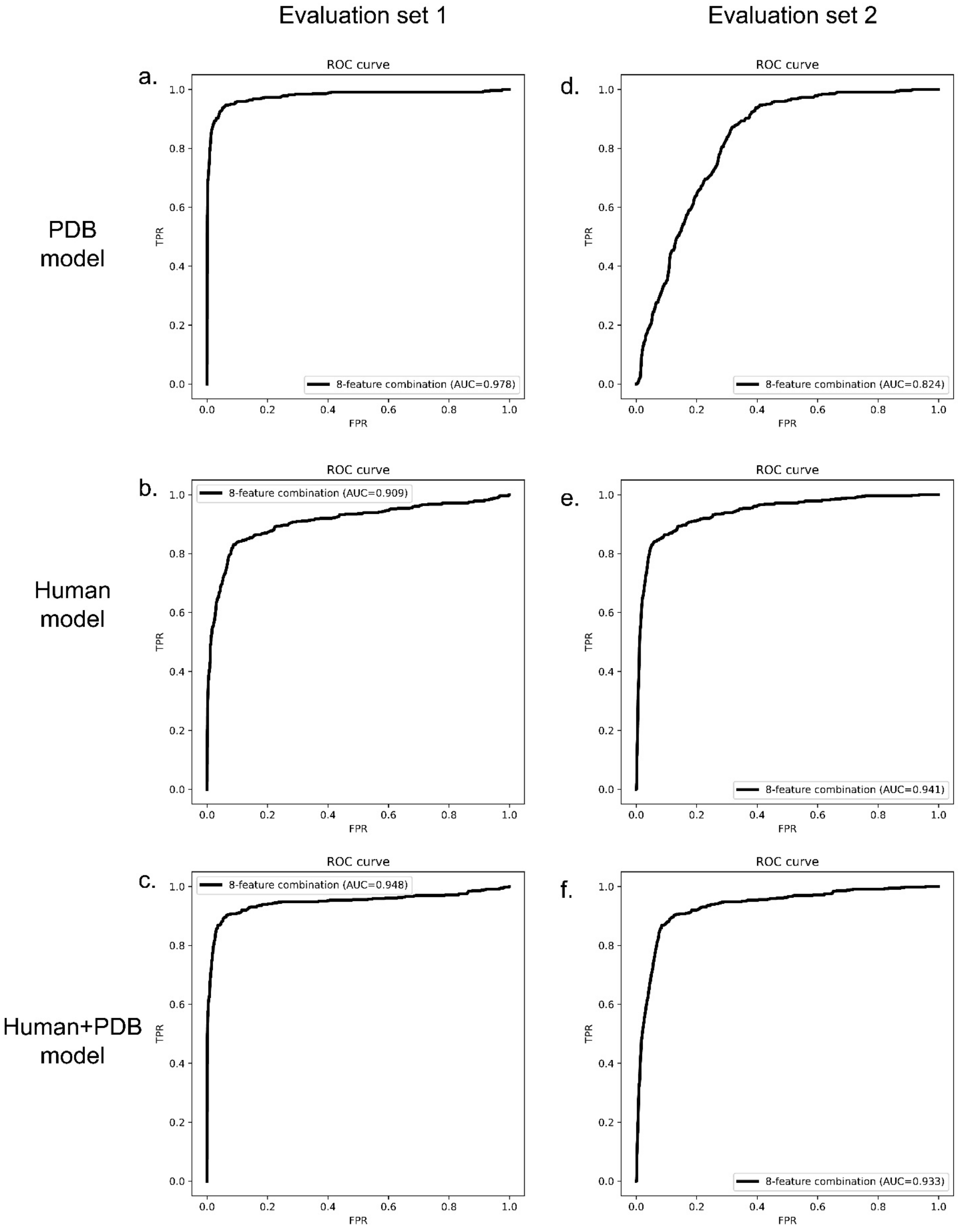
Evaluation set 1
Evaluation set 2
PDB model
Human model
Human+PDB model
a.
b.
c.
d.
e.
f.
ROC curve
TPR
FPR
8-feature combination (AUC=0.978)
8-feature combination (AUC=0.909)
8-feature combination (AUC=0.948)
8-feature combination (AUC=0.824)
8-feature combination (AUC=0.941)
8-feature combination (AUC=0.933)

**Figure S3**. Final model performance via ROC curves, for 3 models. ROC curves on Evaluation set 1 (left) and Evaluation set 2 (right) for 3 different models: (a,d) PDB model, (b,e) Human model and (c,f) Human+PDB model.

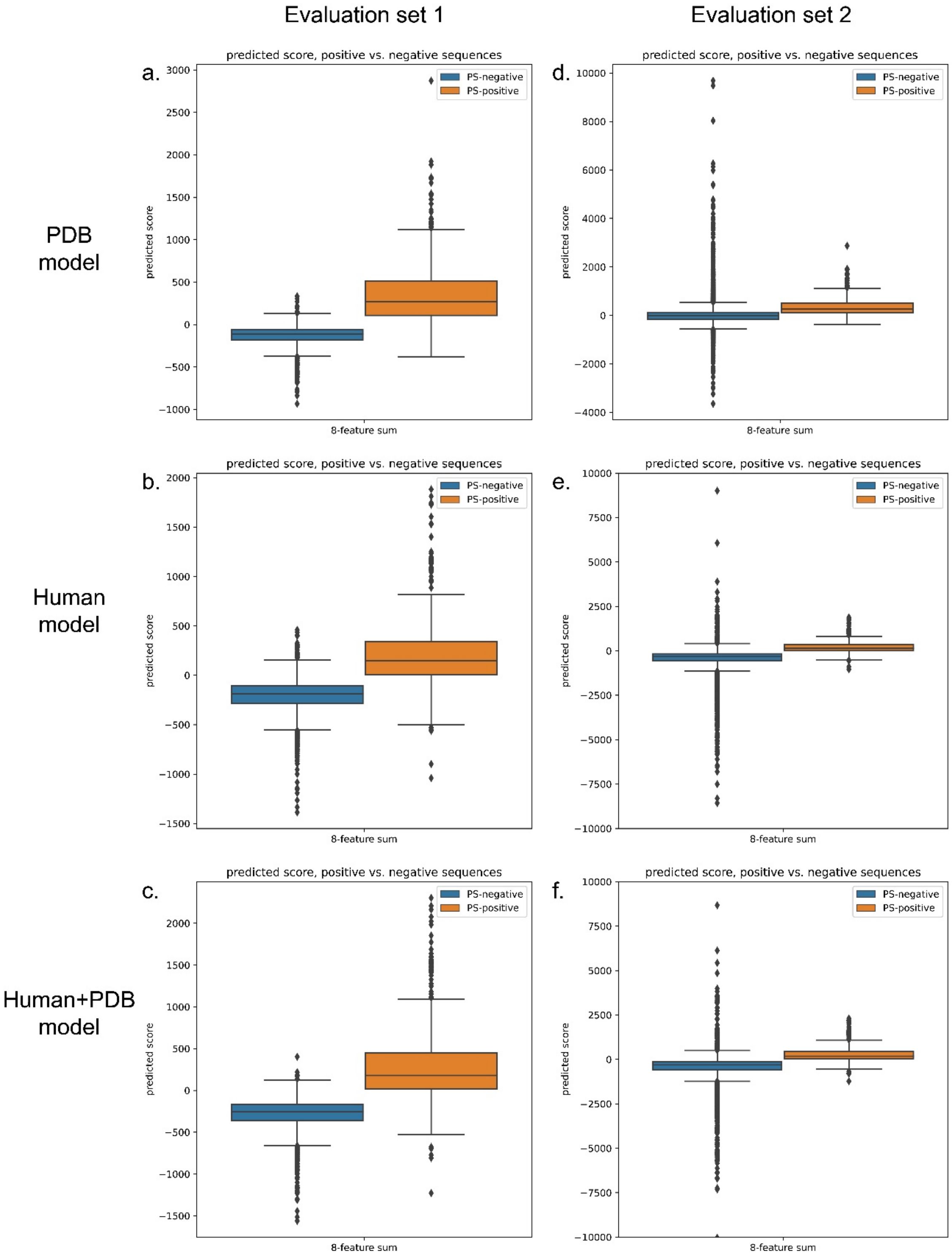


**Figure S4**. Final model performance via boxplots, for 3 models. Predicted score boxplots of positive vs. negative sequences on Evaluation set 1 (left) and Evaluation set 2 (right) for 3 different models: (a,d) PDB model, (b,e) Human model, and (c, f) Human+PDB model.

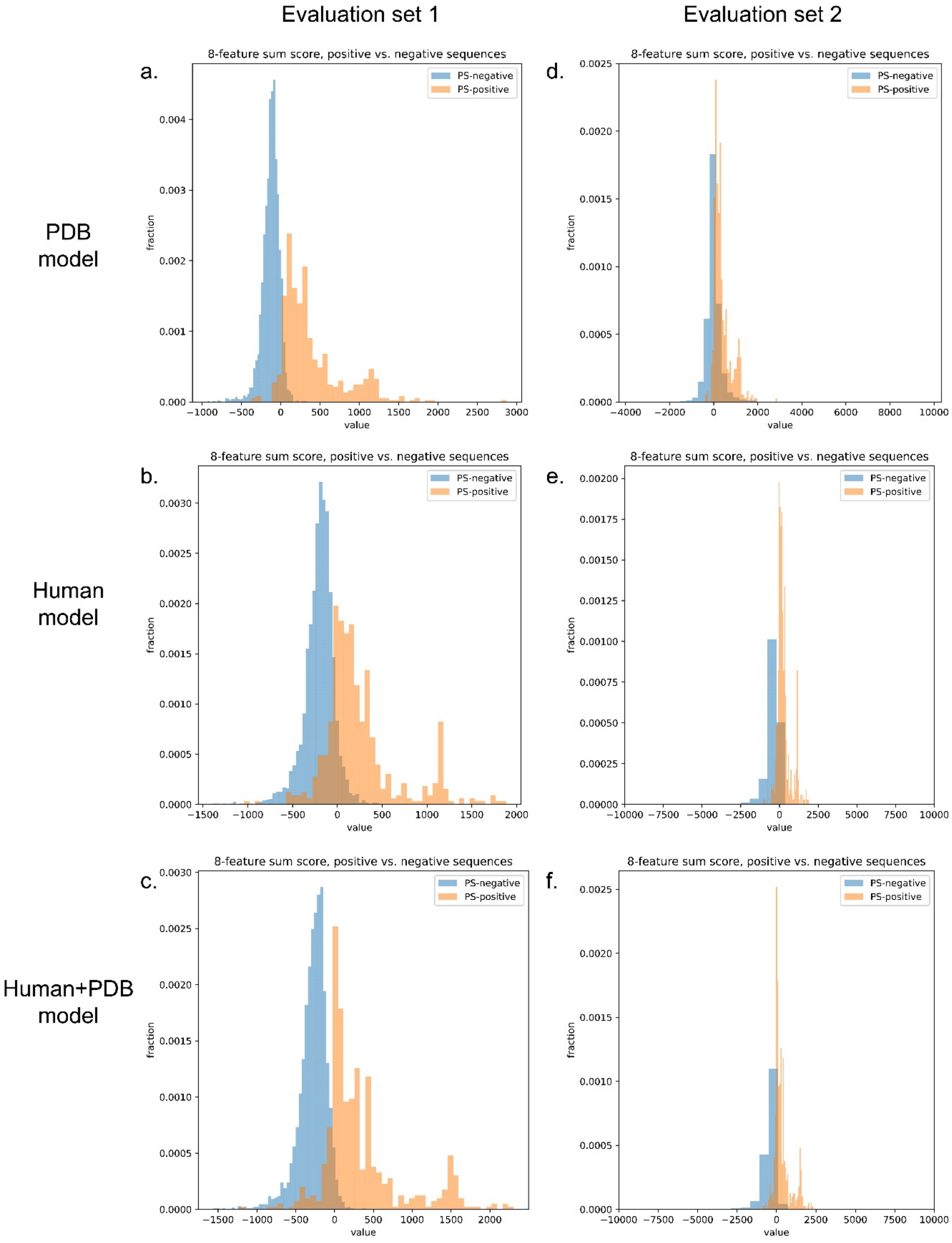


**Figure S5**. Final model performance via histograms, for 3 models. Distribution histograms of positive vs. negative sequences on Evaluation set 1 (left) and Evaluation set 2 (right) for 3 different models: (a,d) PDB model, (b,e) Human model, and (c,f) Human+PDB model.

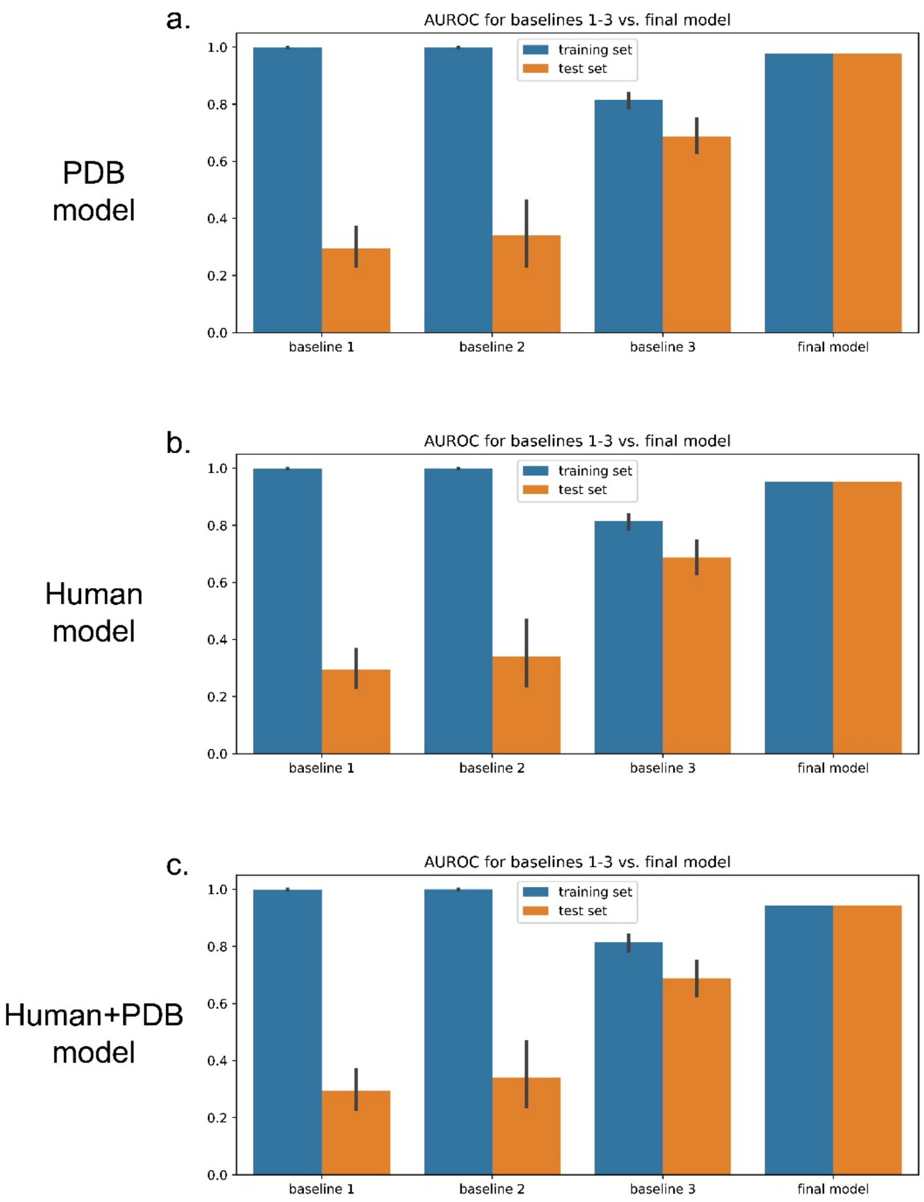


**Figure S6**. Comparison of three training baselines and the final predictor models for validation, for the 3 models. Performance comparisons are shown for (a) PDB model, (b) Human model, and (c) Human+PDB model. Baseline 1 was created by providing random values from a normal distribution N(0, 1) in the weight training step instead of providing PDB-based physical feature values into the genetic algorithm. Baseline 2 was created by providing random values from the distribution of residue-specific physical feature values instead of providing sequence-based physical feature values. Baseline 3 was created by optimizing 1 weight for 20 residue types for each physical feature (removing residue specificity) during training instead of optimizing 20 weights for 20 residue types for each physical feature.

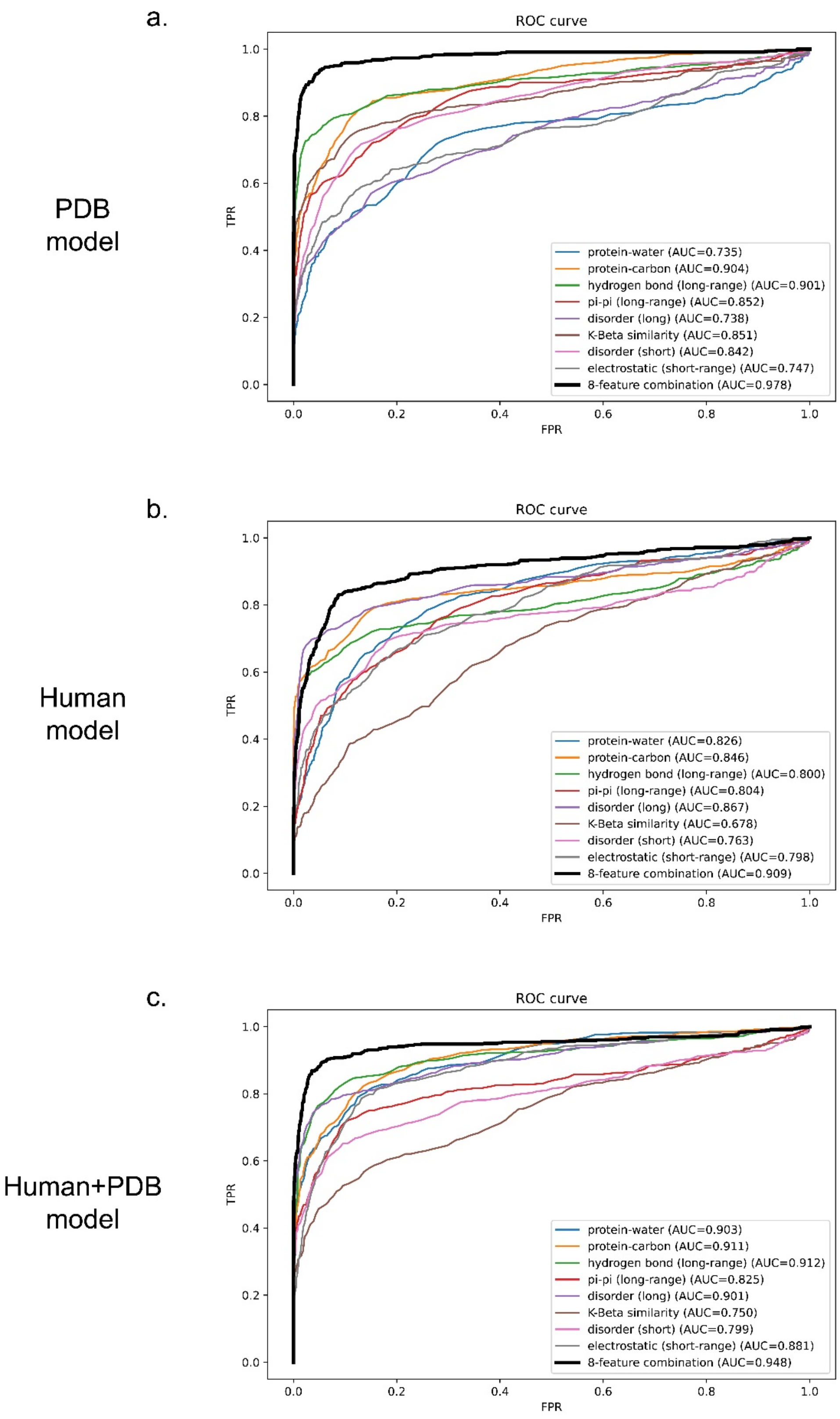
a.
PDB model
ROC curve
TPR
FPR
protein-water (AUC=0.735)
protein-carbon (AUC=0.904)
hydrogen bond (long-range) (AUC=0.901)
pi-pi (long-range) (AUC=0.852)
disorder (long) (AUC=0.738)
K-Beta similarity (AUC=0.851)
disorder (short) (AUC=0.842)
electrostatic (short-range) (AUC=0.747)
8-feature combination (AUC=0.978)
b.
Human model
ROC curve
TPR
FPR
protein-water (AUC=0.826)
protein-carbon (AUC=0.846)
hydrogen bond (long-range) (AUC=0.800)
pi-pi (long-range) (AUC=0.804)
disorder (long) (AUC=0.867)
K-Beta similarity (AUC=0.678)
disorder (short) (AUC=0.763)
electrostatic (short-range) (AUC=0.798)
8-feature combination (AUC=0.909)
c.
Human+PDB model
ROC curve
TPR
FPR
protein-water (AUC=0.903)
protein-carbon (AUC=0.911)
hydrogen bond (long-range) (AUC=0.912)
pi-pi (long-range) (AUC=0.825)
disorder (long) (AUC=0.901)
K-Beta similarity (AUC=0.750)
disorder (short) (AUC=0.799)
electrostatic (short-range) (AUC=0.881)
8-feature combination (AUC=0.948)

**Figure S7**. Comparison of performance via ROC curves of predictors trained on 8 features vs. 1 feature, for 3 models: (a) PDB model, (b) Human Model, and (c) Human+PDB model.

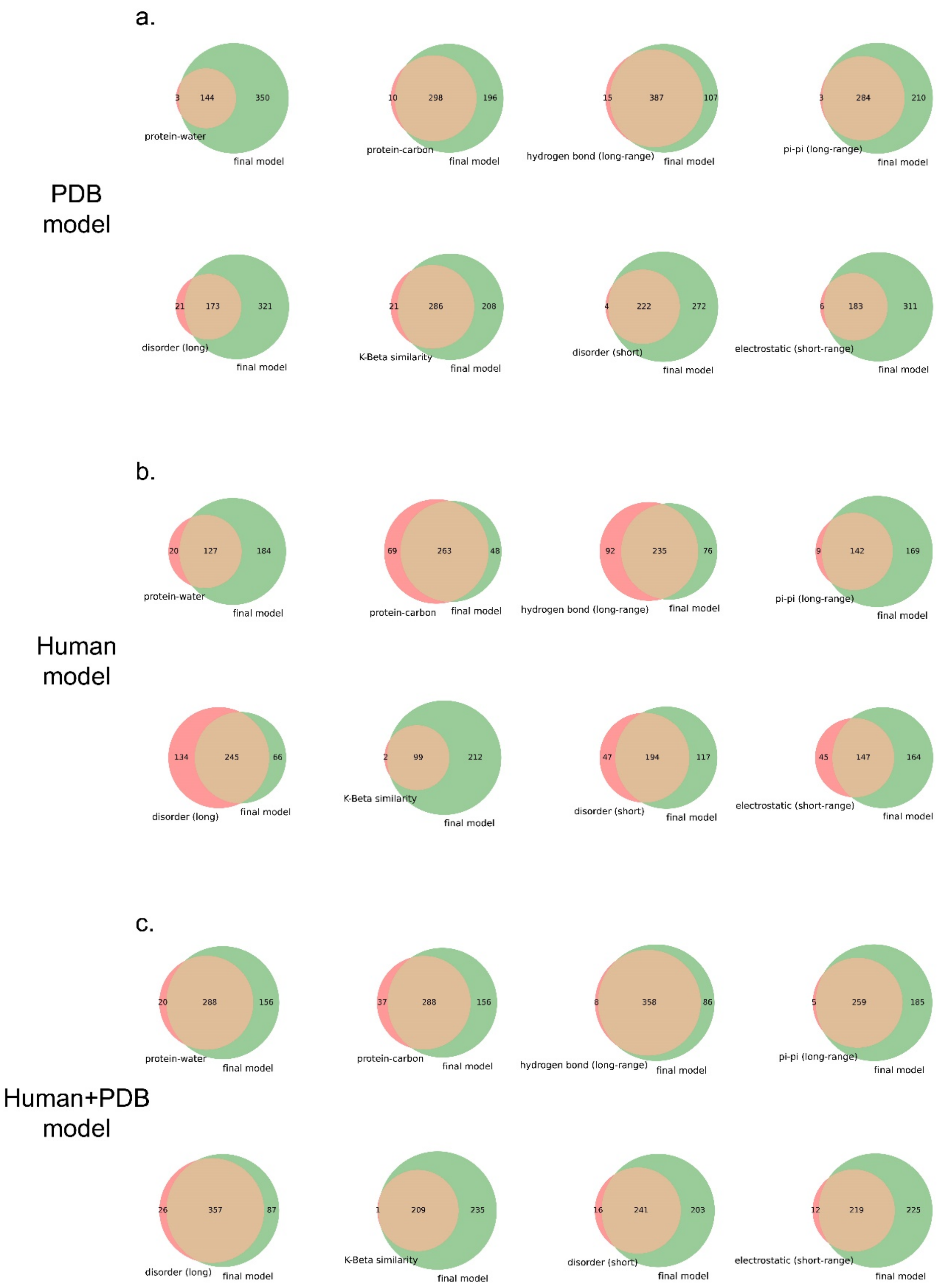


**Figure S8**. Comparison of performance via Venn diagrams of predictors trained on 8 features vs. 1 feature for the 3 models. Venn diagrams showing the coverage overlaps of PS-positive sequences by 1-feature predictors vs. the 8-feature predictor at a confidence threshold that returns 2% of the PDB, for 3 models: (a) PDB model, (b) Human model, and (c) Human+PDB model.

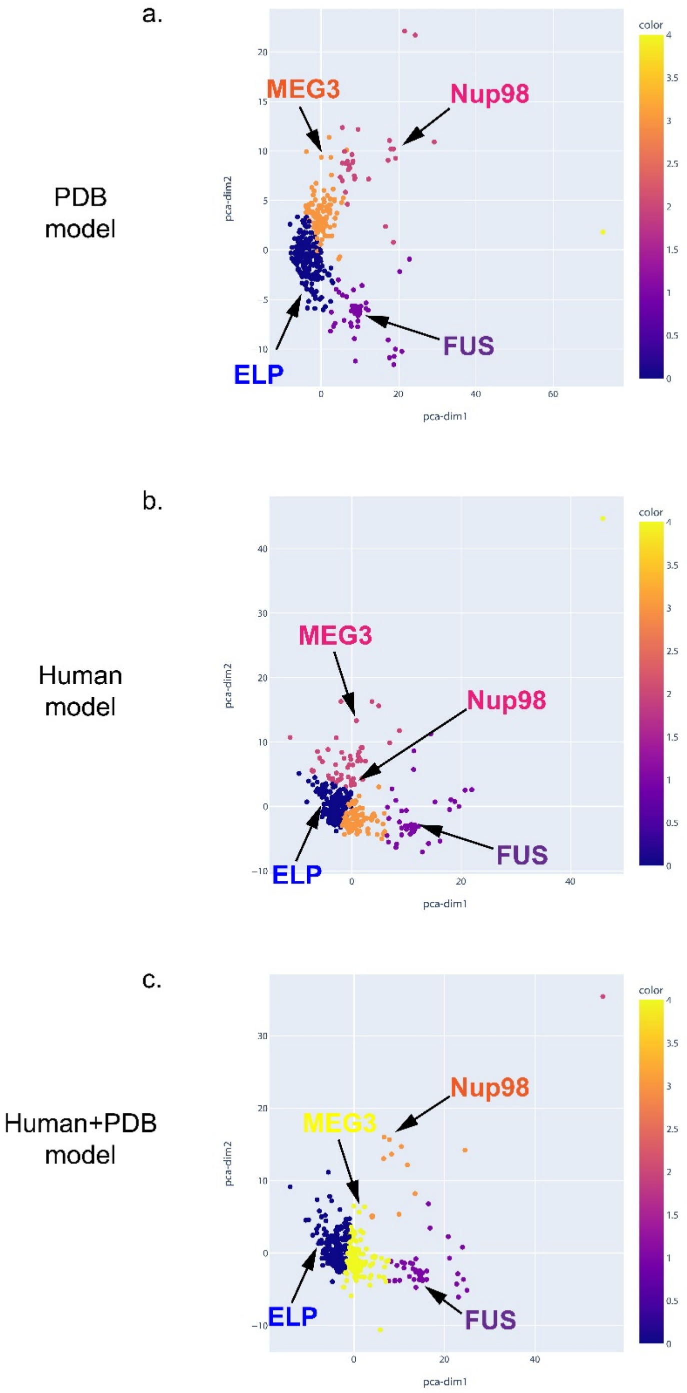


**Figure S9**. Feature score-based clustering for PS-positive proteins for the 3 models. Plots of 2 abstracted dimensions for clustering based on feature z-scores, showing the separation of different types of phase-separating sequences, for 3 models: (a) PDB model, (b) Human model, and (c) Human+PDB model.

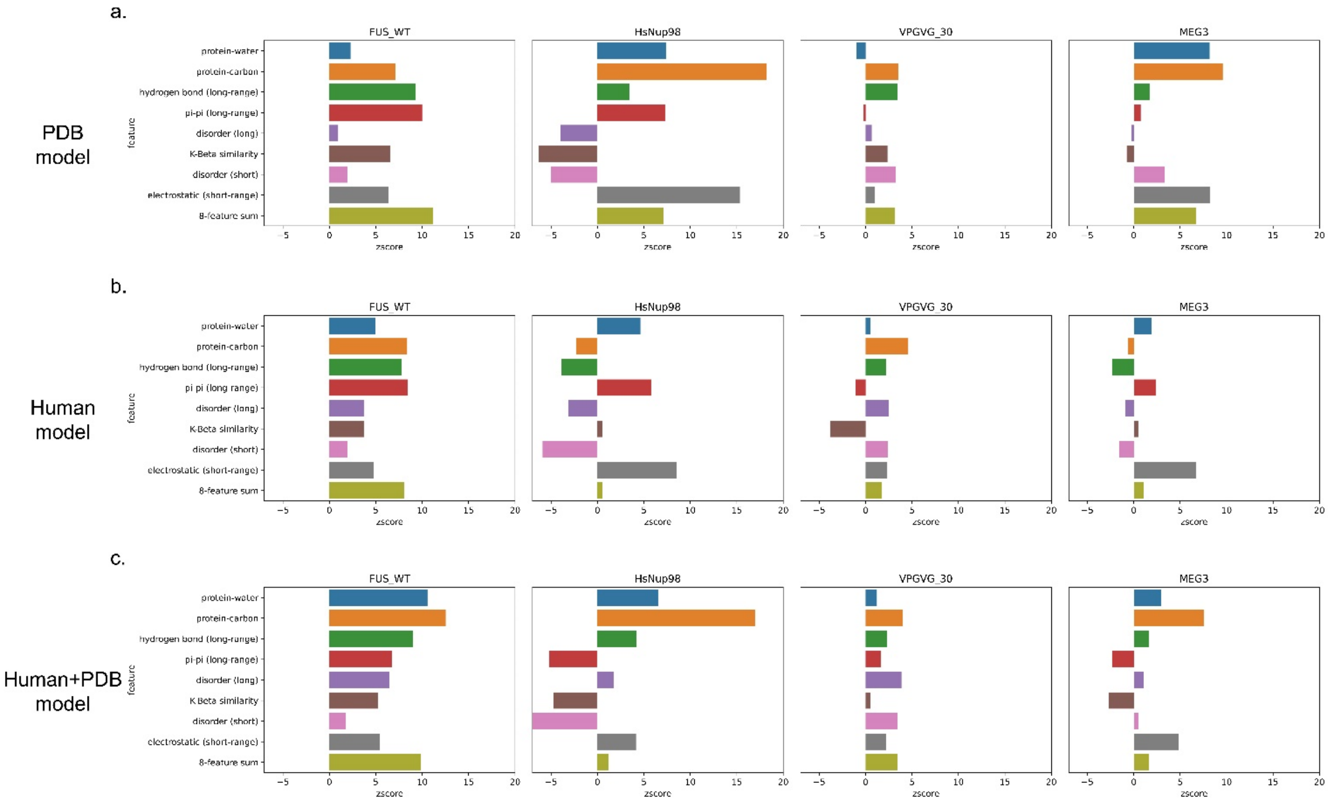


**Figure S10**. Feature score breakdown for example sequences from distinct clusters of PS-positive proteins for the 3 models. The score breakdown of 4 example sequences from 4 clusters in Fig S8 is shown for FUS (human), Nup98 (human), elastin-like peptide (ELP, VPGVG_30, 30 repeats of VPGVG) and MEG-3 (*C. elegans*) for 3 models: (a) PDB model, (b) Human model, and (c) Human+PDB model.

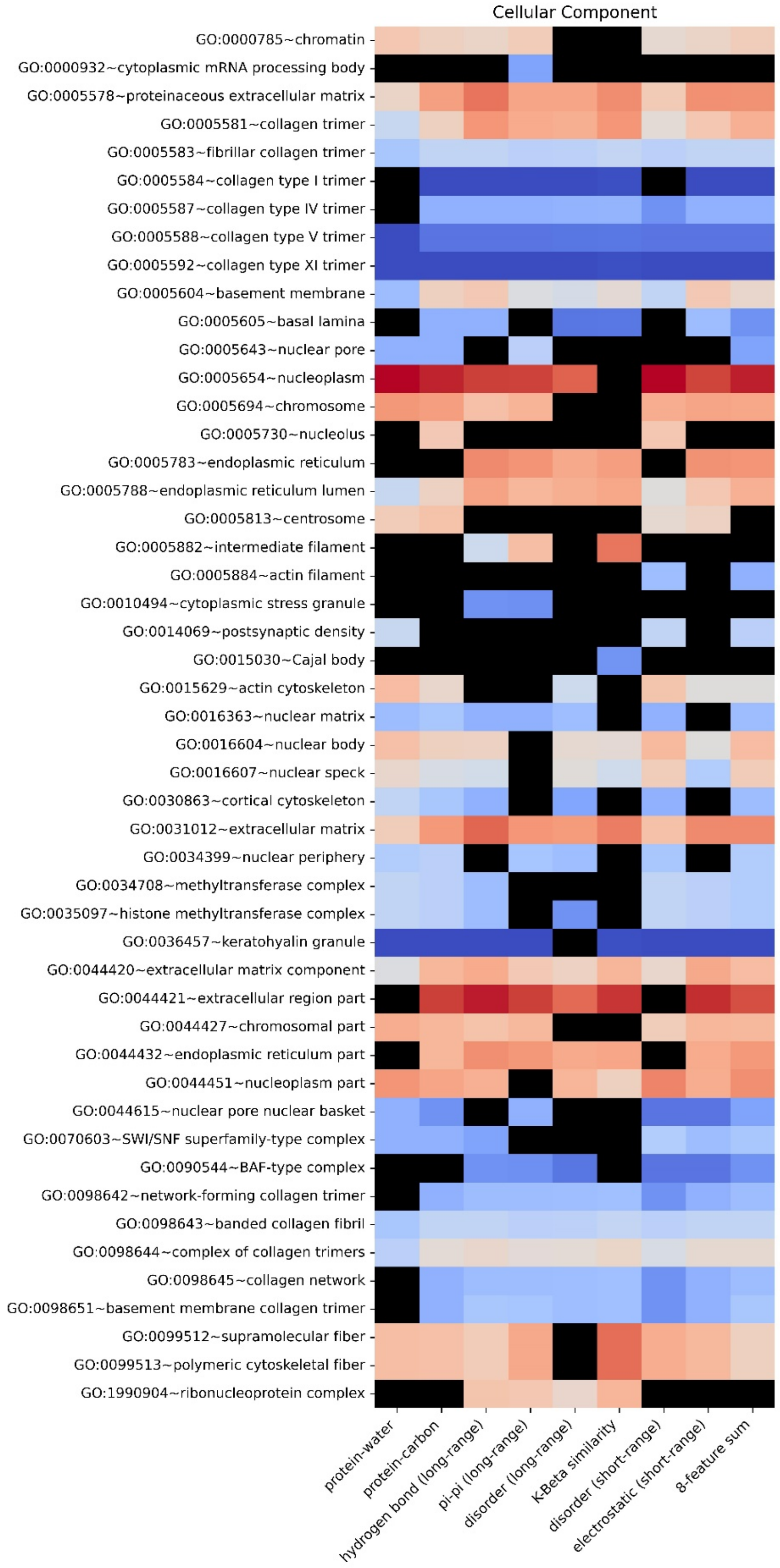


**Figure S11**. Enrichment heatmap by GO functional annotations for different features for the PDB model. Heatmap showing the enrichment of proteins with a given functional annotation that fall under a 2% confidence threshold for each single feature score and 8-feature sum score. The color gradient shows the natural logarithm of the enrichment percentage.

**Table S1**. Constructed Training/Test/Evaluation sets and number of sequences in each set.

| set type | set name | # positive samples | # negative samples |
|---|---|---|---|
| Training set | Training set 1 | 305 | 1703, PDB |
| | Training set 2 | 305 | 1703, Human |
| | Training set 3 | 305 | 1703, PDB + Human |
| Test set | Test set 1 | 260 | 1703, PDB |
| | Test set 2 | 260 | 1703, Human |
| | Test set 3 | 260 | 1703, PDB + Human |
| Evaluation set | Evaluation set 1 | 565 | 16794, PDB |
| | Evaluation set 2 | 565 | 20380, Human |

**Table S2**. 16 physical features with corresponding definitions.

<table>
<tr><th>Physical Feature</th><th>Definition</th></tr>
<tr><td>pi-pi (short-range)</td><td rowspan="2">Pi-Pi contacts were defined using the method in Vernon et al. [1], and then divided into short range and long range by sequence separation. Less than 5 residues apart is defined as short range, and greater than or equal to 5 is defined as long range.</td></tr>
<tr><td>pi-pi (long-range)</td></tr>
<tr><td>protein-water</td><td rowspan="2">Water and carbon counts were calculated only for the subset of proteins in our training set that have a total number of water molecules greater than the number of protein residues. This captures almost all models with resolution <= 1.8 but removes lower resolution models. Counts are measured for residues in their crystallographic context (measurement includes atoms from symmetry partners).</td></tr>
<tr><td>protein-carbon</td></tr>
<tr><td>sec. structure (helices)</td><td rowspan="2">DSSP letter code was used for secondary structure assignments, with H/G used for helix, E for strand, and all others binned to loop.</td></tr>
<tr><td>sec. structure (strands)</td></tr>
<tr><td>disorder (long)</td><td rowspan="2">For identifying disordered residues, a DSSP assignment of “not G/H/E” over a span of at least 3 residues was used to classify residues as loops. These loop residues were then assigned as short disorder if they fall within 3 residues of G/H/E and as long disorder if they do not.</td></tr>
<tr><td>disorder (short)</td></tr>
<tr><td>electrostatic (short-range)</td><td rowspan="2">Phenix.reduce [2] was used to complete PDB structures by adding hydrogen atoms and charge interactions were calculated using the following pseudocode, with partial charges taken from the Talaris2014 energy function [3].<br><br>Q1 = partial_charge for atom X of amino acid 1<br>q2 = partial_charge for atom Y of amino acid 2<br><br>absF = 330.0 * abs(q1*q2)/(distance**2)<br>if q1*q2 < 0.0: absF *= -1.0<br><br>if SequenceSeparation >= 10: add absF to electrostatic (long range)<br>if SequenceSeparation < 10: add absF to electrostatic (short range)<br><br>Final per-residue values were then binned as follows:<br>bin = np.clip(int( round( residue_value / 16.0 ) ), -9, 9)</td></tr>
<tr><td>electrostatic (long-range)</td></tr>
</table>

<table>
<tr><td>hydrogen bond (short-range)</td><td rowspan="2">Structures probed for hydrogen OH-N hydrogen bonds using phenix [2], with the following commands used to extract hydrogen bond information.<br>phenix.reduce -Quiet -FLIP [pdb file] > ./PHENIX_ALL/PHENIXL.pdb phenix.probe "NITROGEN,OXYGEN,HYDROGEN" -Quiet -ONEDOTeach -NOCLASHOUT -SUMMARY -NOVDWOUT ./PHENIX_ALL/PHENIXL.pdb | grep greentint > ./N17.PHENIX/HLIST.'+pdb<br><br>Bonds were than classified as short range and long range by sequence separation (short range < 5, long range ≥ 5).</td></tr>
<tr><td>hydrogen bond (long-range)</td></tr>
<tr><td>cation-pi (short-range)</td><td rowspan="2">We reran the electrostatic scores after adding arbitrary partial charges to the surfaces of aromatic rings, with a partial charge value of -0.05 added 0.85 Å above and below the plane of the ring for each atom, counterbalanced by a partial charge of 0.1 at the atom.<br><br>The cation pi-score is then taken from the difference between this modified score and the unmodified electrostatic score.</td></tr>
<tr><td>cation-pi (long-range)</td></tr>
<tr><td>K-Beta similarity</td><td rowspan="2">Superpositions to kinked beta fibrils were made for chain A in each of 5 structures, PDB IDs 6bwz, 6bxv, 6bxx, 6bzm, and 6bzp. The full chain of each was superimposed to every overlapping window (same number of residues as the chain with none missing) in our PDB training set and kinked beta similarity was measured for each individual PDB residue by taking the minimum CA-RMSD over all measurements the residue was involved in.<br><br>Residues were then classified as K-Beta similar if the minimum CA-RMSD was under 1.0 Å and as K-Beta dissimilar if it was over 2.0 Å.</td></tr>
<tr><td>K-Beta non-similarity</td></tr>
</table>

**Table S3**. The “signs” of 16 features determined by AUROC direction during individual feature training. Feature with a positive correlation to phase separation prediction are in black and those with a negative correlation to phase separation prediction are in red.

| feature name | initial AUROC | final AUROC | feature sign |
|---|---|---|---|
| pi-pi (short-range) | 0.713 | 0.952 | + |
| pi-pi (long-range) | 0.834 | 0.968 | + |
| protein-water | 0.806 | 0.966 | + |
| protein-carbon | 0.155 | 0.035 | - |
| sec. structure (helices) | 0.240 | 0.054 | - |
| sec. structure (strands) | 0.787 | 0.949 | + |
| disorder (long) | 0.904 | 0.961 | + |
| disorder (short) | 0.691 | 0.954 | + |
| electrostatic (short-range) | 0.854 | 0.959 | + |
| electrostatic (long-range) | 0.263 | 0.048 | - |
| hydrogen bond (short-range) | 0.696 | 0.955 | + |
| hydrogen bond (long-range) | 0.794 | 0.967 | + |
| cation-pi (short-range) | 0.355 | 0.039 | - |
| cation-pi (long-range) | 0.226 | 0.033 | - |
| K-Beta similarity | 0.833 | 0.938 | + |
| K-Beta non-similarity | 0.239 | 0.042 | - |

**Table S4**. The AUROC of different trained feature combinations.

| feature combination | AUROC (training set) | AUROC (test set) |
|---|---|---|
| 16 features | 0.959 | 0.935 |
| 12 features | 0.962 | 0.933 |
| final model (8 features) | 0.969 | 0.942 |
| final model (8 features) trained on training set 1+ test set 1 ("PDB model"), using Evaluation set 1 | 0.978 | |
| final model (8 features) trained on training set 1+ test set 1 ("PDB model"), using Evaluation set 2 | 0.824 | |
| final model (8 features) trained on training set 2 + test set 2 ("human model"), using Evaluation set 1 | 0.908 | |
| final model (8 features) trained on training set 2 + test set 2 ("human model"), using Evaluation set 2 | 0.941 | |
| final model (8 features) trained on training set 3+ test set 3 ("human + PDB model"), using Evaluation set 1 | 0.947 | |
| final model (8 features) trained on training set 3+ test set 3 ("human + PDB model"), using Evaluation set 2 | 0.933 | |

### Separate Supplementary Files

One attached Excel file contains, on separate tabs, **Tables S5-S10**.
**Table S5**. GO enrichment analysis.
**Table S6**. Uniprot IDs of 6102 sequences from human proteome using CRAPome as filtering method.
**Table S7**. Uniprot IDs of 3406 sequences from PDB base.
**Table S8**. Uniprot IDs of 3406 sequences randomly selected from human base in Table S6.
**Table S9**. Uniprot IDs of 6812 sequences from PDB+human base.
**Table S10**. Detailed information of 565 LLPS-positive sequences with PMID of each sequence's paper.

**File S1**. 565 PS-positive sequences (fasta file).
**File S2**. 16794 PDB sequences (fasta file).
**File S3**. 20380 Human sequences (fasta file).

### Supplementary Material References


1 Vernon, R. M. et al. Pi-Pi contacts are an overlooked protein feature relevant to phase separation. eLife 7, e31486, doi:10.7554/eLife.31486 (2018).
2 Adams, P. D. et al. PHENIX: a comprehensive Python-based system for macromolecular structure solution. Acta Crystallographica Section D: Biological Crystallography 66, 213-221 (2010).
3 O'Meara, M. J. et al. Combined covalent-electrostatic model of hydrogen bonding improves structure prediction with Rosetta. Journal of chemical theory and computation 11, 609-622 (2015).
